# Deciphering *Arabidopsis thaliana* Acclimation to Sublethal Combined Abiotic Stresses

**DOI:** 10.1101/2024.06.13.598895

**Authors:** Zhang Jiang, Ava Verhoeven, Yihong Li, Romy Geertsma, Rashmi Sasidharan, Martijn van Zanten

**Affiliations:** Plant Stress Resilience, Institute of Environmental Biology, Utrecht University, Padualaan 8, 3584CH Utrecht, the Netherlands

## Abstract

Plants are frequently exposed to environmental challenges. Responses to sub-lethal abiotic stress combinations are complex and often distinct from responses to individual stresses and remain poorly understood. Investigating traits and molecular factors mediating acclimation to stress combinations is essential for the development of climate change-resilient field crops. Here, we studied the morphological, physiological, and molecular responses of *Arabidopsis thaliana* to i) co-occurring high temperature and drought and ii) flooding sequentially followed by drought, both of which have increased in frequency due to climate change. A set of 15 physiological and morphological traits were assessed during single and combined stresses. By combining these comprehensive trait analyses with transcriptome characterization, we established the generally additive effects of simultaneous or sequential stresses on plant morphology and physiology compared to the corresponding individual stresses. Although drought had a mild effect in both stress combinations, a unique transcriptome signature emerged upon combination with high temperature simultaneously or flooding sequentially. Molecular processes identified as important for multi-stress resilience included plastid-nucleus communication, ABA signaling and photo-acclimation. Based on the RNA-seq data, a set of 39 genes was identified as potential multi-stress response regulators. Mutants were tested to validate their contribution to plant survival and phenotypic acclimation under combined stress. We confirmed the involvement of several genes in regulating phenotypic acclimation traits. Among the novel identified factors are EARLY FLOWERING 6 (ELF6) and ARABIDOPSIS TÓXICOS EN LEVADURA (ATL80), with significant effects on plant growth, leaf development and plant survival (wilting) during high-temperature drought and post-submergence drought respectively.

## Introduction

Increased occurrences of abiotic stresses such as heat, drought and flooding due to climate change are negatively impacting crop productivity (Kopecká et al., 2023; Shabbir et al., 2022) and thereby food security (Cushman et al., 2022; Kumar et al., 2022; Masson-Delmotte et al., 2021). Although the effect of such abiotic stresses on plant functioning has been extensively studied (Harb et al., 2010; Larkindale et al., 2005; van Veen et al., 2016) the focus has been predominantly on single stressors, often involving abrupt transfer of plants into harsh stressful conditions. In natural or agricultural settings, abiotic stresses seldom happen in isolation and usually occur at a sublethal severity (Bailey-Serres and Colmer, 2014; De Smet et al., 2021; Van Dooren et al., 2020; Zhang et al., 2020) with multiple stresses coinciding simultaneously or sequentially (Mittler, 2006; Suzuki et al., 2014).

Co-occurring stresses can elicit synergistic or antagonistic effects or are a blend of responses to individual stresses. Hence responsiveness cannot be simply deduced by taking the sum of the responses of individual stresses (Balfagón et al., 2019; Mittler, 2006; Rivero et al., 2022; Rizhsky et al., 2004; Suzuki et al., 2014; Zandalinas et al., 2020). For instance, high temperature induces stomatal opening to allow leaf cooling through transpiration, while under drought stomatal conductance is reduced to prevent unnecessary water loss. However, when high temperature and drought occur concurrently a signalling conflict occurs and the stomata tend to be closed (Mittler and Blumwald, 2010; Rizhsky et al., 2004; Zandalinas et al., 2016a, 2016b).

The precise acclimation strategy fitting a given stress combination is determined by various (sometimes limiting) parameters, such as plant genotype, developmental age, duration and severity of stresses (Rivero et al., 2022; Sinha et al., 2021; Zandalinas et al., 2021, 2016b; Zandalinas and Mittler, 2022; Zhou et al., 2017). Additionally, the sequence of events plays a pivotal role (Banti et al., 2008; Kong and Henry, 2016). For example, soil salinity pretreatment can alleviate effects of subsequent low-temperature damage in tomato plants, as salt stress-triggered signal cascades can activate downstream overlapping signal transduction pathways (*e.g.*, reactive oxygen species (ROS), abscisic acid (ABA) and low-temperature signaling) that enhance photosynthetic acclimation under low-temperatures (Yang et al., 2022).

Despite its importance, multi-stress acclimation mechanisms remain poorly understood (Rivero et al., 2022; Zandalinas et al., 2020; Zandalinas and Mittler, 2022). Here we characterized the effects of two stress combinations typical of climate change in *Arabidopsis thaliana.* These were: i) combined high temperature and drought and ii) submergence followed by drought, and their corresponding individual stresses. A comprehensive assessment of the impact of these stress combinations on plant morphology, physiology, and transcriptome revealed different, in general additive, effects on plant growth, development and physiology compared to the corresponding individual stresses. The unique phenotypic characteristics correlated with distinct transcriptomic signatures involving synchronous regulation of multiple generic processes such as plastid-nucleus communication (retrograde signaling), ABA signaling and photo-acclimation.

A reverse genetics approach was used to validate the contribution of a set of 44 candidate genes to multi-stress acclimation responses. Effects were confirmed for several factors, including FLOWERING 6 (ELF6) and ARABIDOPSIS TÓXICOS EN LEVADURA (ATL80) that impact on specific subsets of traits related to growth, development and water status maintenance during high temperature/drought or submergence/drought combinations. Our results reveal that acclimation to different sublethal combinatorial stresses is likely coordinated by distinct candidate regulatory genes acting as either positive or negative regulators of multi-stress acclimation traits.

## Results

### Combined and sequential abiotic stresses elicit additive responses

*Arabidopsis* Columbia-0 (Col-0) plants were subjected to two different stress combinations upon reaching the 10-leaf stage (LS10): i) high ambient temperature (27 °C) + progressive drought (HTD) and ii) submergence, followed by progressive drought after de-submergence (PSD; 21 °C) (Fig. 1A), control conditions (C; 21 °C) and corresponding single stresses: high temperature (HT; 27 °C), submergence (S; 21 °C), whereafter plants were well-watered during the post-submergence phase (PS) (Fig. 1A). Drought (D) was imposed progressively by withholding watering. After 5 days, the average soil water content (%SWC) dropped to 50–65% field water capacity (= 100% saturation), and to 30–40% after 10 days (Supplemental Fig. S1, A and B). Plants subjected to HTD and PSD displayed earlier wilting than when D was imposed in isolation (Supplemental Fig. S2, A-C) and produced fewer new leaves than those subjected to the corresponding single stresses (Supplemental Fig. S2, D and E). Drought-treated plants (D and PSD) showed increased chlorophyll content compared to the relative controls (C and PS) (Supplemental Figs. S3 and S4). Based on these responses, we focused on the 0, 5, 10 and 15 days following stress initiation time points in the remainder of this study (Fig. 1A).

**Figure 1.**
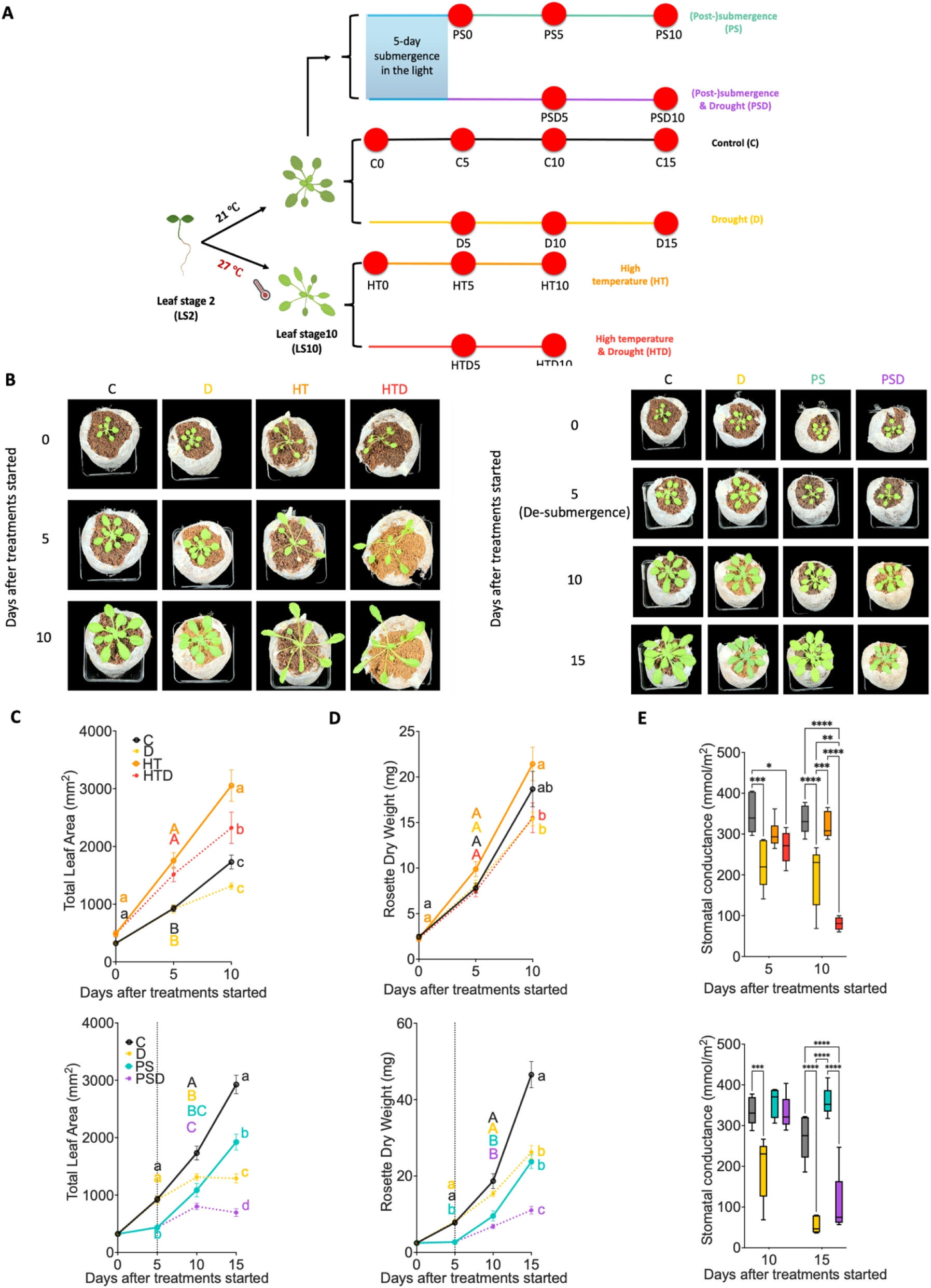
Effects of combined and sequential sublethal stresses and corresponding single stresses on phenotypic traits. **(A)** Experimental setup. Harvest moments and measurements (indicated by red dots) were typically at 0, 5, 10 and 15 (C and D) days after the drought (D, HTD and PSD) treatment started at average %SWC of 100%, 50–65%, 30– 40% and ∼20% respectively (Supplemental Fig. S1, A and B). For C, HT and PS the %SWC were maintained at the saturated level (95%-100%) across all timepoints. **(B)** Representative images of plants on Jiffy 7c coconut pellet growth substrate, subjected to combined (left) and sequential (right) stresses and the corresponding single stresses at 0, 5, 10 and 15 days, counted from the moment that the treatments started. **(C-E)** Total leaf area of the whole rosette (C, n = 14-20), rosette dry weight (D, n = 15-18) and stomatal conductance of young leaves (E, n = 5-6) of plants subjected to combined (upper row) and sequential (lower row) stresses and the relevant single stresses at the corresponding harvesting time points. Error bars indicate means ± SEM. Letters denote significant differences between different treatments at the same time points (p < 0.05, 2-way ANOVA with Tukey’s Post-hoc test). The dashed vertical lines indicate the moment plants were de-submerged. Boxes indicate boundaries of the second and third quartiles (Q) of the data distribution. Black horizontal bars indicate median and whiskers Q1 and Q4 values within 1.5 times the interquartile range. Asterisks represent significant differences (*p < 0.05, **p < 0.01, ***p < 0.001, ****p < 0.001, 2-way ANOVA with Tukey’s Post-hoc test). Abbreviations; C; control (black lines and symbols), D; drought (yellow), HT; high temperature (orange), HTD; high temperature & drought (red), S/PS; 5-day submergence / (post-)submergence & recovery (blue), PSD; (post-)submergence & drought (purple).

#### High temperature and drought

D did not significantly hamper rosette growth nor dry weight accumulation, but triggered primary root lengthening (Fig. 1, B-D; Supplemental Fig. S5D). Plants in HT showed the expected typical thermomorphogenic phenotype (Quint et al., 2016), including elongated petioles and hyponastic leaves (Fig. 1B; Supplemental Figs. S5, A-C and S6). HTD treatment resulted in smaller rosettes and shorter petioles and leaf blades compared with plants exposed to HT, but rosette size, petiole length and leaf blade length under HTD exceeded the values observed under D (Fig. 1C; Supplemental Figs. S5, B-C and S6). Plants grown in HT or D did not differ in dry weight compared to plants in C (Fig. 1D). However, HTD resulted in significant dry-weight reduction after 10 days compared to HT, but not C (Fig. 1D). Primary root length was significantly promoted in HTD compared to the individual stresses (Supplemental Fig. S5D). Together, this indicates that during the subjection to HTD, D and HT stresses interact.

Plants contained relatively more water under high temperature conditions (HT and HTD) than at control temperatures (C and D) (Supplemental Fig. S7A). Moreover, the differences in leaf surface temperature between ‘control’ and drought-treated plants at 27 °C (HT and HTD) were greater than at 21 °C (C and D) (Supplemental Fig. S7, B and C). Together with an accelerated drought-induced temperature increase at 27 °C compared to 21 °C (Supplemental Fig. S7, B and C), these results indicate that the drought effect on leaf temperature was more pronounced when combined with high temperature. However, stomatal conductance of plants in C and HT was not significantly different, whereas C and HTD did differ significantly, with HTD having lower stomatal conductance (Fig. 1E). D caused, as expected, a reduction in relative water content corresponding with a significant decline in stomatal conductance compared to C (Fig. 1E; Supplemental Fig. S7A).

No differences in Malondialdehyde (MDA) content were observed across treatments up to day 10 (Supplemental Fig. S7D), suggesting that no significant oxidative stress occurs. However enhanced ion leakage in HT-treated plants (Supplemental Fig. S7E) points to membrane damage at high temperatures (Nielsen et al., 1997; Sharma et al., 2012). The addition of 10-days drought (HTD) tempered this effect, while no effect was observed when D was applied in isolation (Supplemental Fig. S7E).

#### Submergence and drought

5-days complete submergence (S) led to severely inhibited rosette growth compared to non-submerged plants (Fig. 1, B-D; Supplemental Figs. S8, S9 and S10). During post-submergence plants could recover if irrigation was applied (PS) (Fig. 1, B-D; Supplemental Figs. S8, S9 and S10). D had a significant negative effect on total rosette area and dry weight, compared to C after 10 days (Fig. 1, C and D). When D was imposed following de-submergence (PSD), total rosette area and dry weight accumulation were further reduced compared to PS (Fig. 1, C and D; Supplemental Fig. S10, A and E). In contrast to D, S restricted primary root lengthening whereas root elongation was enhanced under PSD (Supplemental Figs. S8D and S10F).

D negatively affected relative water content after 15 days (Supplemental Fig. S11A), which was enhanced under PSD (Supplemental Fig. S11A and S12A). Both D and PSD treatments resulted in a reduction of stomatal conductance (Fig. 1E; Supplemental Fig. S12B). Accordingly, leaf temperature was raised under these conditions (Supplemental Fig. S11, B-C).

Drought increased MDA accumulation, especially when combined with prior submergence (PSD) (Supplemental Figs. S11D and S12C). Yet, no clear effect of PSD, D or PS was observed on ion leakage (Supplemental Figs. S11E and S12D).

### Transcriptomic responses elicited by combined and sequential stresses

Next, a comparative RNA-sequencing approach was taken using young leaves sampled 0, 5, and 10 days after stress initiation (Fig. 1A) to uncover molecular processes underlying responses to combined and sequential stresses. In total, 58 samples, distributed over 15 different harvesting timepoints were sequenced, resulting in 3,628,334,664 detected reads (Supplemental Fig. S13A). On average, 72% of the mapped reads aligned to the Arabidopsis transcriptome (5.4 to 61.8 million reads per sample). For approximately one-third of sequenced samples less than 50% of the reads mapped to the Arabidopsis transcriptome (Supplemental Fig. S13A). 98.2% of the unmapped reads were identified as RNA segments of *Arabidopsis Latent Virus 1* (ArLV1 RNA1 and RNA2; GenBank accessions MH899120.1 and MH899121.1, respectively) (Verhoeven et al., 2023). Although this virus does not evoke visible symptoms in plants (Verhoeven et al., 2023), viral infections could impact the host transcriptome (Shates et al., 2019). We therefore compared samples with reads that mapped more than 50% with those mapping less than 50% to the Arabidopsis transcriptome on day 5 C treatment (Supplemental Fig. S13B). No differentially expressed genes between both fractions were detected, implying that ArLV1 does not significantly impact the plant transcriptome.

PCA analysis revealed significant changes across time (PCO1, explaining 29,88% of observed variation) and treatments (separated over PCO1 and PCO2) (Supplemental Fig. S14A). As expected, individual biological replicates of the same timepoint and stress treatment clustered together (Supplemental Figs. S14 and S15). Superimposing the fraction of reads mapping to the Arabidopsis genome per sample on the PCA revealed no clear pattern (Supplemental Fig. S14B), confirming that ArLV1 contamination did not significantly affect the transcriptome. We therefore decided to continue processing our data using all samples except two (day 5 control) of which the mapping percentages to Arabidopsis transcriptome were below 11% (Supplemental Figs. S13).

### High temperature and submergence alter transcriptome responses to drought

#### High temperature and drought

In response to D, only 3 differentially expressed genes (DEGs; |Log_2_FC| > 0, FDR < 0.05) were detected; *GUARD-CELL-ENRICHED GDSL LIPASE 3* (*GGL3*), *PHOSPHOFRUCTOKINASE 1* (*PFK1*) and *AT5G16990* (Fig. 2A). Under HTD, the drought effect became more apparent (590 and 1090 DEGs after 5 and 10 days of HTD, respectively) (Fig. 2A). Moreover, HTD resulted in 955 unique DEGs that were not detected under D or HT (Fig. 2B). Thus, combined stress evokes a unique transcriptomic response and has greater impact than individual stresses. HT and HTD shared 469 DEGs, suggesting that HT has a more dominant effect on the transcriptome than D in the combined stress (HTD) situation (Fig. 2B). Only 2 DEGs; *PFK1* and *AT5G16990,* were commonly regulated by D, HT and HTD (Fig. 2B), suggesting that these genes might be involved in early stress responses in young leaves.

**Figure 2.**
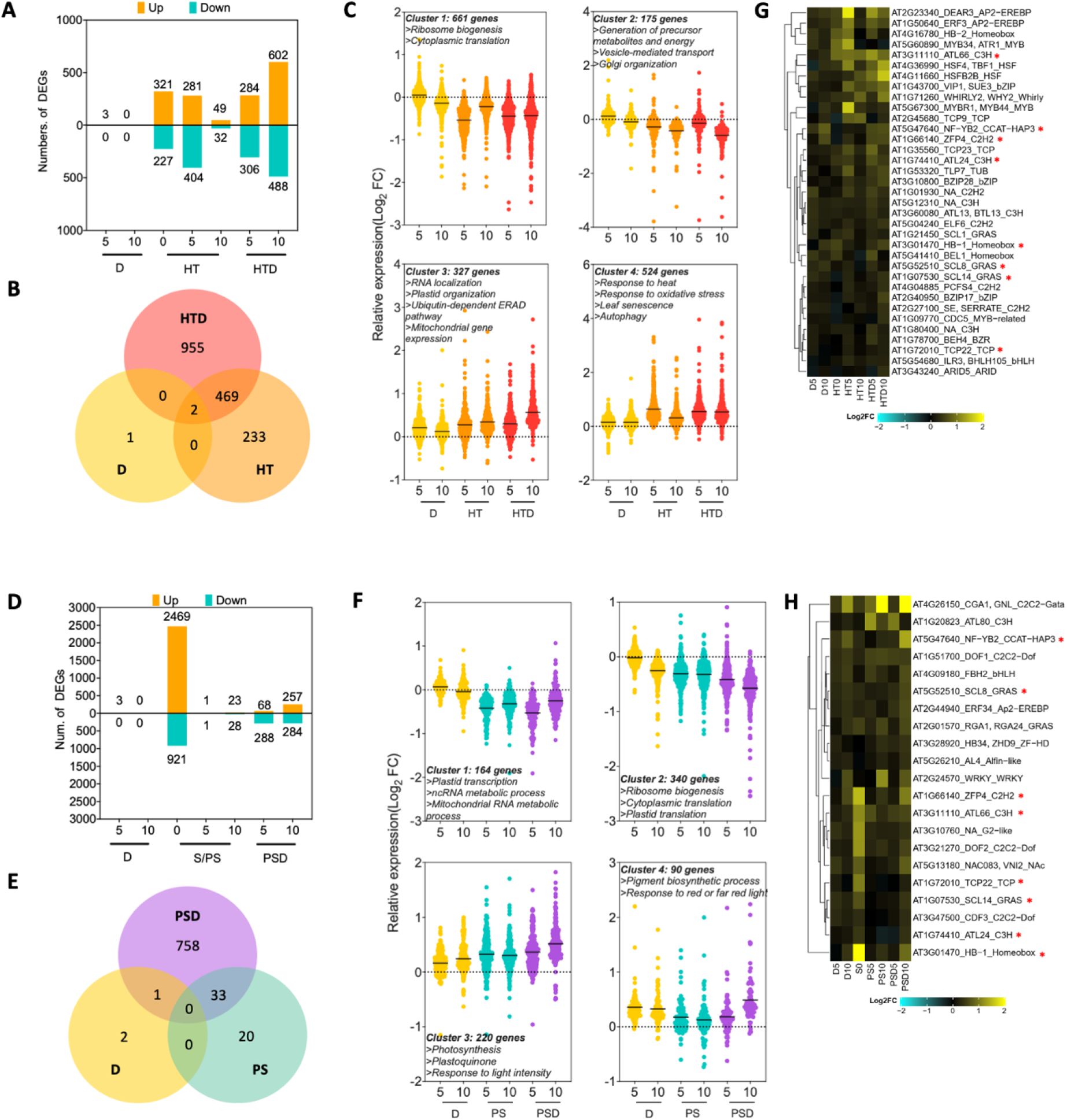
Effect of combined and sequential stresses and corresponding single stresses on the transcriptome. **(A, D)** Number of up-(orange) and down-(green) regulated DEGs (|Log_2_FC| >0, p < 0.05) at day 0 (the pre-growth effect of HT before the drought treatment started), 5, and 10 days after the treatments (D) started (note HT was prolonged), compared to control (C) conditions. **(B, E)** Venn diagrams showing the number of DEGs commonly or differently detected in the combined (HTD), sequential (PSD) and single (D, HT, PS) stresses. **(C, F)** *k-means* clustering presenting expression patterns of identified DEGs at timepoints 5 and 10 days during combined or sequential stresses and the corresponding individual stresses. Violin plots indicate the distributions of relative expression (Log_2_FC) of identified DEGs. Mean values of each violin plot are indicated by a solid black line. Dashed horizontal lines indicate Log_2_FC = 0. Key enriched biological processes identified by Gene Ontology analysis and number of genes assigned to each cluster are listed above or below the graphs. **(G, H)** Hierarchically clustered heatmaps showing relative Log_2_FC levels of transcription factors (TFs) (Supplemental Tables S3 and S4) identified among the upregulated DEGs in the HTD and PSD datasets. For each identified TF, the AGI gene locus ID, the commonly used abbreviation (if available) and TF family the factor belonged to, are listed. Color scales indicate Log_2_FC values (relative to control (C) conditions). Red asterisks indicate overlapping TFs between HTD and PSD. For treatment abbreviations and used colors, see legend of Figure 1.

*K-means* clustering of the 1687 detected DEGs (day 5 and 10) across individual and combined stress treatments revealed 4 groups with distinct expression patterns (Fig. 2C). Cluster 1 and 2 consisted of DEGs downregulated under HT and HTD conditions and unchanged under D. Gene Ontology (GO) annotation revealed that these clusters were dominated by genes associated with ribosome activities and protein processing (Fig. 2C; Supplemental Table S1). DEGs that were upregulated in HT and HTD were in Cluster 3 and 4 (Fig. 2C). These DEGs are associated with RNA processing (particularly alternative splicing), organelle activities including ubiquitin-depended endoplasmic-reticulum-associated protein degradation (ERAD) pathway, mitochondrial activities and stress responsiveness such as response to heat, starvation or oxidative stress and leaf senescence (Fig. 2C; Supplemental Table S1).

#### Submergence and drought

To characterize the effects of submergence followed by drought (PSD), the following comparisons (Log_2_FC in gene expression) were made: i) 5-day submergence (S) compared to pre-submergence (submergence effect), ii) 5 or 10 days of D compared to C (drought effect), iii) 5- or 10-day PS or PSD compared to C respectively, to determine effects of S followed by irrigation (PS) or drought (PSD). This approach ensured comparisons between samples at similar developmental stages.

5-days submergence resulted in 3390 DEGs (2469 up- and 921 downregulated) (Fig. 2D), reflecting considerable transcriptome reconfiguration. Following de-submergence (PS) only a limited number of DEGs were detected after 5 and 10 days (2 and 51 DEGs respectively; Fig. 2D). Upregulation of several core-hypoxia responsive genes (Mustroph et al., 2009) during 5 days S pointed to occurrence of tissue hypoxia (Supplemental Fig. S16). As expected, the upregulation was reversed by re-oxygenation following de-submergence (PS and PSD at day 5 and 10) (Supplemental Fig. S16). D during de-submergence (PSD) resulted in 356 and 541 DEGs at 5 and 10 days of PSD, respectively (Fig. 2D). Like HTD, PSD also led to a unique transcriptomic signature, as the majority of DEGs identified after PSD treatment (758 DEGs) were not affected by PS nor D (Fig. 2E). Only 33 DEGs were shared between PSD and PS (Supplemental Fig. S17).

*k-means* clustering including all 814 DEGs (day 5 and 10 across D, PS and PSD treatments; Fig. 2F), revealed four clusters. Two clusters contained genes downregulated by PS and PSD (Cluster 1 and 2) and two contained genes upregulated in these conditions (Cluster 3 and 4) (Fig. 2F; Supplemental Table S2). GO analysis revealed that Cluster 1 and 2 were dominated by genes associated with nucleic acid, ribosome and translation-related processes. Several of the downregulated GO terms in clusters 1 and 2 were associated with plastid functions, indicating impeded chloroplast activities during post-submergence (PS and PSD). In Cluster 3 photosynthesis-related GO terms were significantly enriched, while in Cluster 4 only two GO categories were enriched (Fig. 2F; Supplemental Table S2). Of note, *CYTOKININ-RESPONSIVE GATA FACTOR 1* (*CGA1*/*GNL*) was shared between these two GO groups.

### Transcription factors mediating acclimation to combined and sequential stresses

In total, 48 different transcription factors (TFs) from 23 TF families were upregulated by HTD (35) or PSD (21), with 8 TFs overlapping between the two groups (Fig. 2, G and H; Supplemental Fig. S18). Only 7 out of 35 HTD-upregulated TFs were also upregulated under HT (Supplemental Table S3) and only 2 out of 21 PSD-upregulated TFs, were induced under 5- or 10-day PS (Supplemental Table S4). This underscores the unique response that is evoked by combined stress application. Considering PSD, of the 19 uniquely induced TFs, 12 were significantly upregulated by 5-day S (Supplemental Table S4). This points to a large effect of prior submergence on the sequential-stress transcriptome signature.

Gene regulatory networks (GRNs) were constructed with the HTD and PSD upregulated gene clusters (Clusters 3 and 4) to identify putative regulatory nodes (Supplemental Fig. S19, A and B). The obtained HTD network consisted of 12913 interactions between 847 regulators (nodes). Only 4169 interactions between 340 regulators were formed in the PSD network (Supplemental Fig. S19, A and B). To identify putative ‘master regulators’, 11 nodes with the highest number of connections with the other regulators in the same GRN were selected. Four of these were shared between the HTD- and PSD-networks: *G-BOX BINDING FACTOR 3* (*GBF3*), *ABSCISIC ACID RESPONSIVE ELEMENT-BINDING FACTOR 1* (*ABF1*), *ABA INSENTITIVE 5* (*ABI5*) and *NF-YB2* (Supplemental Fig. S20, Table). Note that *THYLAKOID LUMEN PROTEIN* (*TLP18.3*) ranked 12^th^ in the degree analysis but is associated with chloroplast function (Järvi et al., 2016; Wu et al., 2011). *GENOMES UNCOUPLED 4* (*GUN4*) and *AT1G66130* had the same number of nodes in GRN, therefore in total 11 overrepresented genes for HTD and PSD are indicated in Supplemental Fig. S19.

### Validation of identified candidate genes by mutant analysis

Next, a reverse genetics approach was taken to validate the effects of identified factors on multi-stress acclimation (Supplemental Table S5). For this, TFs were selected that were upregulated during either HTD or PSD, supplemented with some of the overrepresented genes from the GRNs. In total, 41 confirmed homozygous T-DNA insertion mutants, covering in total 39 candidate genes (28 for HTD and 19 for PSD), were obtained (Supplemental Tables S6 and S7).

#### High temperature and drought

Traits indicative of plant growth and development were measured at 0 and 10 days after stress initiation. Subsequently, the stress treatment was extended until the plants showed wilting symptoms (Fig. 3A). Of the 29 mutants with putative effect on HTD traits (Fig. 3A; Supplemental Table S6), *aba insensitive 5* (*abi5-7*), *early flowering 6* (*elf6-3*), *at5g12310*, *VIRE2-interacting protein 1 (vip1)* and *zinc finger protein 4* (*zfp4*) displayed early wilting phenotypes (Fig. 3B; Supplemental Table S8). As expected, the ABA signaling mutant *abi5-7* (Bi et al., 2017; Dekkers et al., 2016) showed substantially deviating phenotypes under HTD compared to the wildtype (WT) in addition to clear wilting phenotypes (Fig. 3B; Supplemental Table S8). This points to a pivotal role for ABA signaling in modulating plant growth and development under HTD. The *elf6-3* mutant showed accelerated wilting (Fig. 3B; Supplemental Table S8) and a considerable delay in leaf initiation rate.

**Figure 3.**
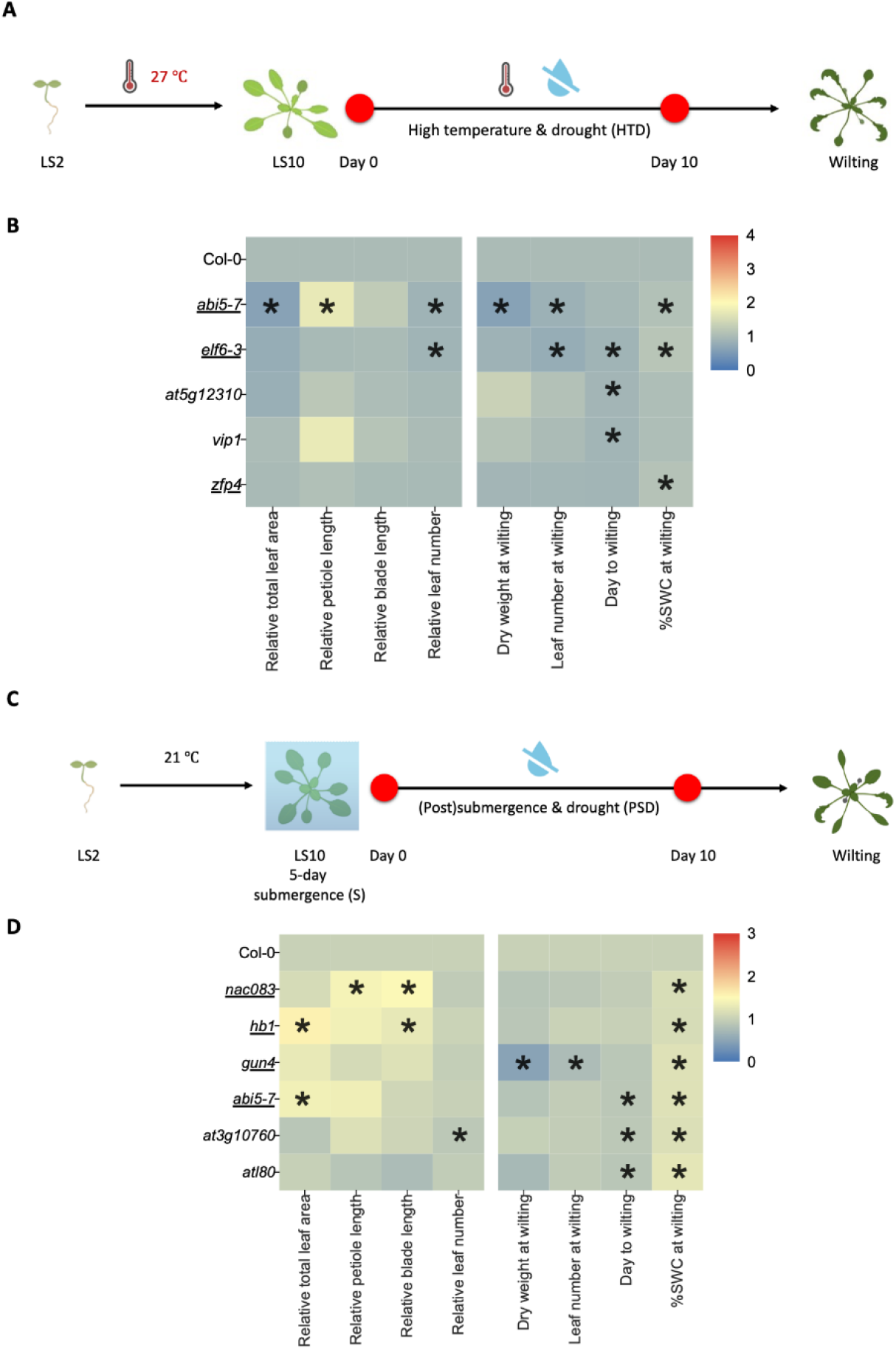
Plant growth, development and wilting traits of selected mutants under combined or sequential stresses. **(A, C)** Experimental scheme for the used reverse genetic analyses. 2-leaf stage (LS2) Arabidopsis mutants and corresponding wild-type (WT) plants were pre-grown at 27 °C (HTD; A) or 21 °C (PSD; C) until they reached the 10-leaf stage (LS10). Subsequently, the plants were subjected to 10 days of combined high temperature and drought conditions (HTD), or 5-day submergence in the light, followed by 10-day drought treatment (PSD) and thereafter were both left unwatered until wilting occurred. Measurements of phenotypic traits were carried out at day 0, day 10 and at the moment of wilting (*i.e.*, timepoint differed per plant). **(B, D)** Heatmaps indicating phenotypic traits of Arabidopsis mutants with either significantly advanced wilting time and/or high %SWC at wilting, relative to the corresponding wild-type plants (Col-0) during the exposure to HTD (B) or PSD (D). The relative values of total leaf area, petiole/blade length and leaf number were calculated by normalizing the data obtained at day 10 by that of day 0 (left four columns in each panel). Dry weight and leaf number at wilting, days to wilting and %SWC at wilting are indicated in the right four columns per panel). The abbreviated names of tested mutants and the wild type plants are indicated per row. Color scales indicate relative values of measured traits (relative to the corresponding wild-type Col-0 plants). Underlined mutants indicate confirmed knockout of the corresponding genes based on previous studies (Supplemental Tables, S6 and S7). Asterisks represent significant differences between mutants and wild-type plants for the particular trait (Supplemental Tables S8 and S9) (p < 0.05, unpaired t-test). n=8-12.

#### Submergence and drought

Of the 20 mutants tested (Fig. 3C; Supplemental Table S7), six (*NAC domain containing protein 83* (*nac083*), *homeobox1* (*hb1*), *gun4*, *at3g10760*, *abi5-7* and *arabidopsis tóxicos en levadura* (*atl80*)) exhibited early wilting phenotypes under PSD (Fig. 3D; Supplemental Table S9). The *nac083*, *hb1* and *abi5-7* mutants exhibited a significant increase in total leaf area and/or leaf length (Fig. 3D; Supplemental Table S9). Both *gun4* and *at3g10760* showed reduced leaf initiation rates. Notably, *gun4* weighed only ∼50% of the (dry) weight at the moment of wilting compared to WT (Supplemental Table S9). Despite wilting at a high %SWC level, *atl80* mutant phenotypic traits were comparable to WT (Fig. 3D; Supplemental Table S9).

For mutants exhibiting early wilting during HTD (*elf6-3*, *at5g12310*, *vip1* and *zfp4*) or PSD (*atl80, nac083, at3g10760, hb1, gun4*), experiments were extended with inclusion of the corresponding individual stresses (Fig. 4 and Fig. 5; Supplemental Figs. S20, S21, S22 and S23). The *abi5-7* mutant was omitted as ABA signaling is already studied in great detail in plant acclimation to diverse stresses (Skubacz et al., 2016; Wang et al., 2021).

**Figure 4.**
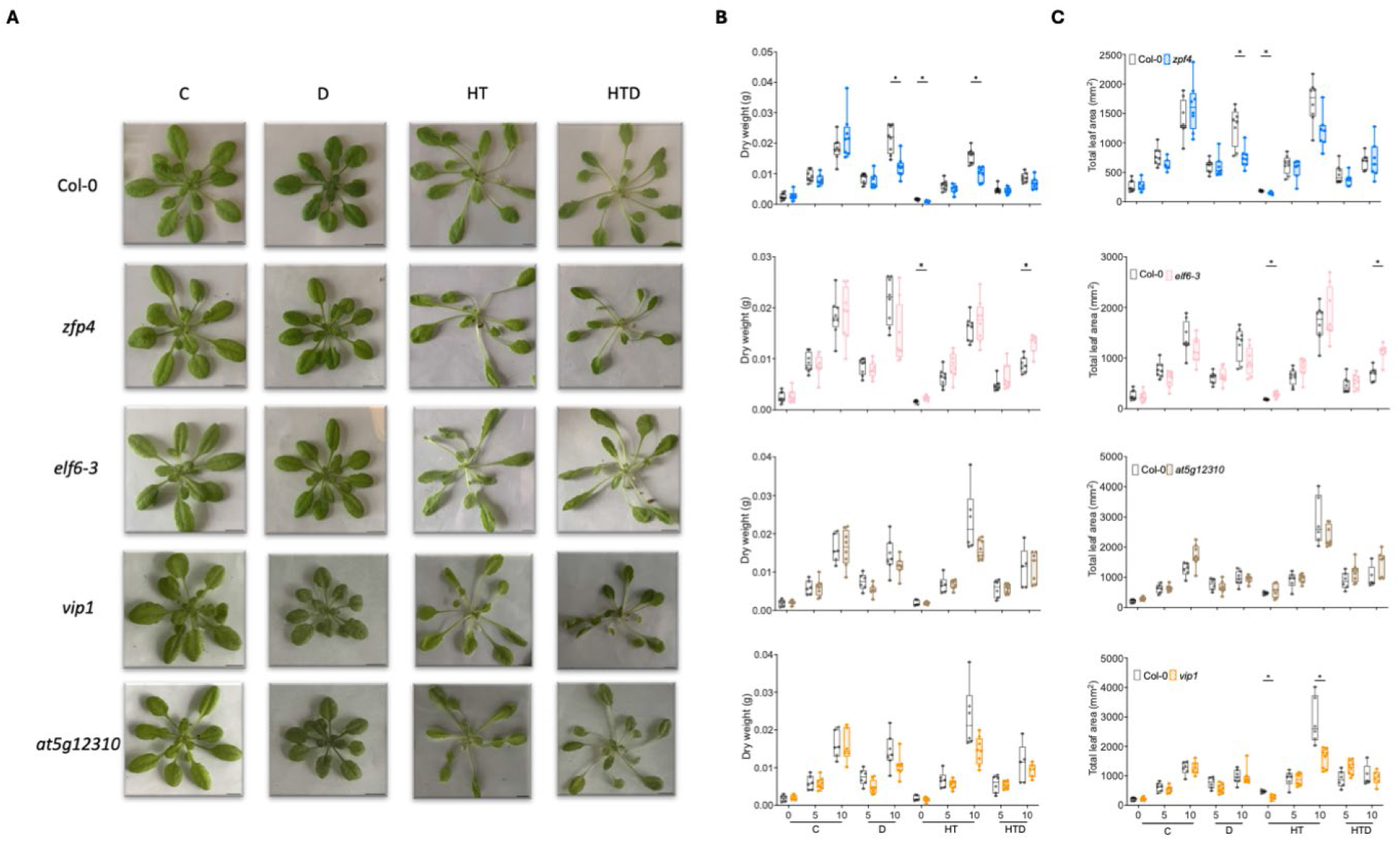
Effect of combined and individual stresses on phenotypic traits of selected Arabidopsis mutants. **(A)** Representative images of whole rosettes of selected mutants and corresponding wild-type plants (Col-0) grown on Jiffy 7c coconut pellet substrate, subjected to single and combined stresses at 10 days after the treatments initiated. Scale bars indicate 1 cm. **(B, C)** Rosette dry weight (B) and total leaf area (C) of Arabidopsis mutants (*zfp4* (blue), *elf6-3* (pink), *at5g12310* (brown), and *vip1* (orange)) and the wild-type plants (Col-0; gray). For plants that wilted before the harvesting time points, traits were measured at the day of wilting onset. Boxes indicate boundaries of the second and third quartiles (Q) of the data distribution. Horizontal bars indicate median and whiskers Q1 and Q4 values within 1.5 times the interquartile range. Asterisks represent significant differences between the mutant and the corresponding wild-type plants within the same time point (p < 0.05, multiple unpaired t-test with Holm-Šídák correction). Numbers indicate days after treatments started. For treatment abbreviations and used colors, see legend of Figure 1. n= 5-10.

**Figure 5.**
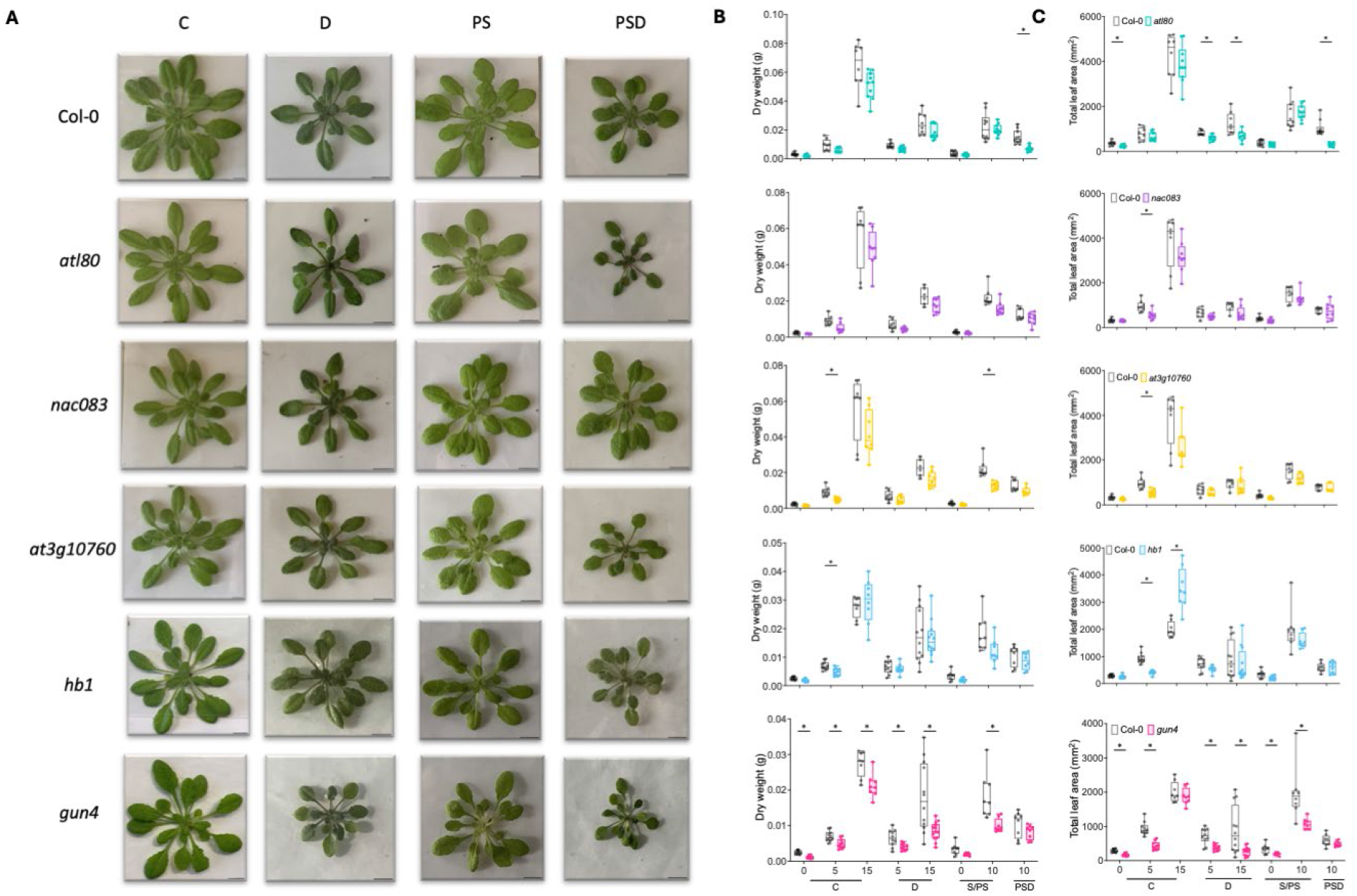
Effect of sequential and individual stresses on phenotypic traits of selected Arabidopsis mutants. **(A)** Representative images of whole rosettes of selected mutants and the corresponding wild-type plants (Col-0) grown on Jiffy 7c coconut pellet substrate, subjected to single and sequential stresses 15 days after the treatments started. Scale bars indicate 1 cm. **(B, C)** Rosette dry weight (B) and total leaf area (C) of Arabidopsis mutants (*atl80* (aqua), *nac083* (purple), *at3g10760* (yellow), *hb1* (azure) and *gun4* (magenta)) and the wild-type plants (Col-0; gray). For plants that wilted before the harvesting time points, the traits were measured on the day of wilting onset. Boxes indicate boundaries of the second and third quartiles (Q) of the data distribution. Horizontal bars indicate median and whiskers Q1 and Q4 values within 1.5 times the interquartile range. Asterisks represent significant differences between the mutant and the corresponding wild-type plants within the same time point (p < 0.05, multiple unpaired t-test with Holm-Šídák correction). Numbers indicate days after treatment started. For treatment abbreviations and used colors, see legend of Figure 1. n=6-13.

#### Validation of high temperature and drought mutants

Compared to WT, the *elf6-3* mutant had higher dry weight and rosette area under HTD treatment (Fig. 4, B and C). This mutant also displayed enhanced dry weight and had a larger rosette area at 0-day (pre-growth) at HT, but neither at later HT time points nor under C and D treatments (Fig. 4, B and C). This indicates that ELF6 is a negative regulator of plant growth and rosette expansion under HTD. The *zfp4* mutant showed reduced dry weight and total leaf area under 0-day HT and 10- day D, whereas *vip1* only showed a smaller total leaf area than the WT under 0- or 10-day HT (Fig. 4, B and C). This suggests that both ZFP4 and VIP1 mediate plant growth during D and/or HT but not HTD.

In HTD, all mutants with accelerated wilting displayed a significant decrease in chlorophyll content (Supplemental Fig. S21A). In contrast, both *vip1* and *at5g12310* displayed higher chlorophyll content under 10-day D (Supplemental Fig. S21A). No significant differences in chlorophyll abundance were observed between mutants and WT during HT single treatment (Supplemental Fig. S21A). Under HT and HTD, stomatal conductance of all mutants was comparable to WT (Supplemental Fig. S21B). Notably, 5-day D significantly reduced stomatal conductance in both *vip1* and *at5g12310* mutants compared to WT (Supplemental Fig. S21B).

#### Validation of post-submergence and drought mutants

The *atl80* mutant exhibited decreased dry weight and total leaf area compared to the WT when subjected to PSD and developed smaller rosettes and shorter leaves during C and D treatment (Fig. 5, A-C; Supplemental Fig. S22, A and B). Both *nac083* and *hb1* mutants had similar dry weights and total leaf area as the WT during individual (D, PS) and sequential stresses (PSD) (Fig. 5, B and C). However, prolonged PS treatment resulted in decreased blade length and leaf number in *nac083* (Supplemental Fig. S22, B and C). As reflected by reduced dry weight accumulation, blade length and leaf number during 10-day PS, the *at3g10760* mutant was unable to recover to the same extent as WT after submergence (Fig. 5B; Supplemental Fig. S22, B and C). The *gun4* mutant appeared highly sensitive to D and PS, as dry weight, total leaf area, leaf length and leaf initiation rate were significantly reduced (Fig. 5, B and C; Supplemental Fig. S22, B and C). However, *gun4* also exhibited a significant reduction in these traits under C (Fig. 5, B and C; Supplemental Fig. S22, B and C). Chlorophyll content and stomatal conductance remained unchanged for most mutants (Supplemental Fig. S23, A and B), except *gun4,* which had higher chlorophyll and stomatal conductance under 10-day C (Supplemental Fig. S23A).

### Principal component analysis reveals trait correlations and validates mutant effects

A PCA analysis incorporating all measured traits of all mutants and WTs revealed strong correlation between leaf development traits (including relative petiole (RPL) and blade (RBL) length and total leaf area (RLA)) (Fig. 6, A and B). Likewise, relative leaf number (RLN), leaf number at wilting (LNW) and relative dry weight at wilting (DWW) were positively correlated (Fig. 6, A and B). As expected, days to wilting (DTW) and %SWC at wilting (%SWCW) negatively correlated (Fig. 6, A and B), indicating that fast-wilting mutants indeed also generally contained a high %SWC at the moment of wilting.

**Figure 6.**
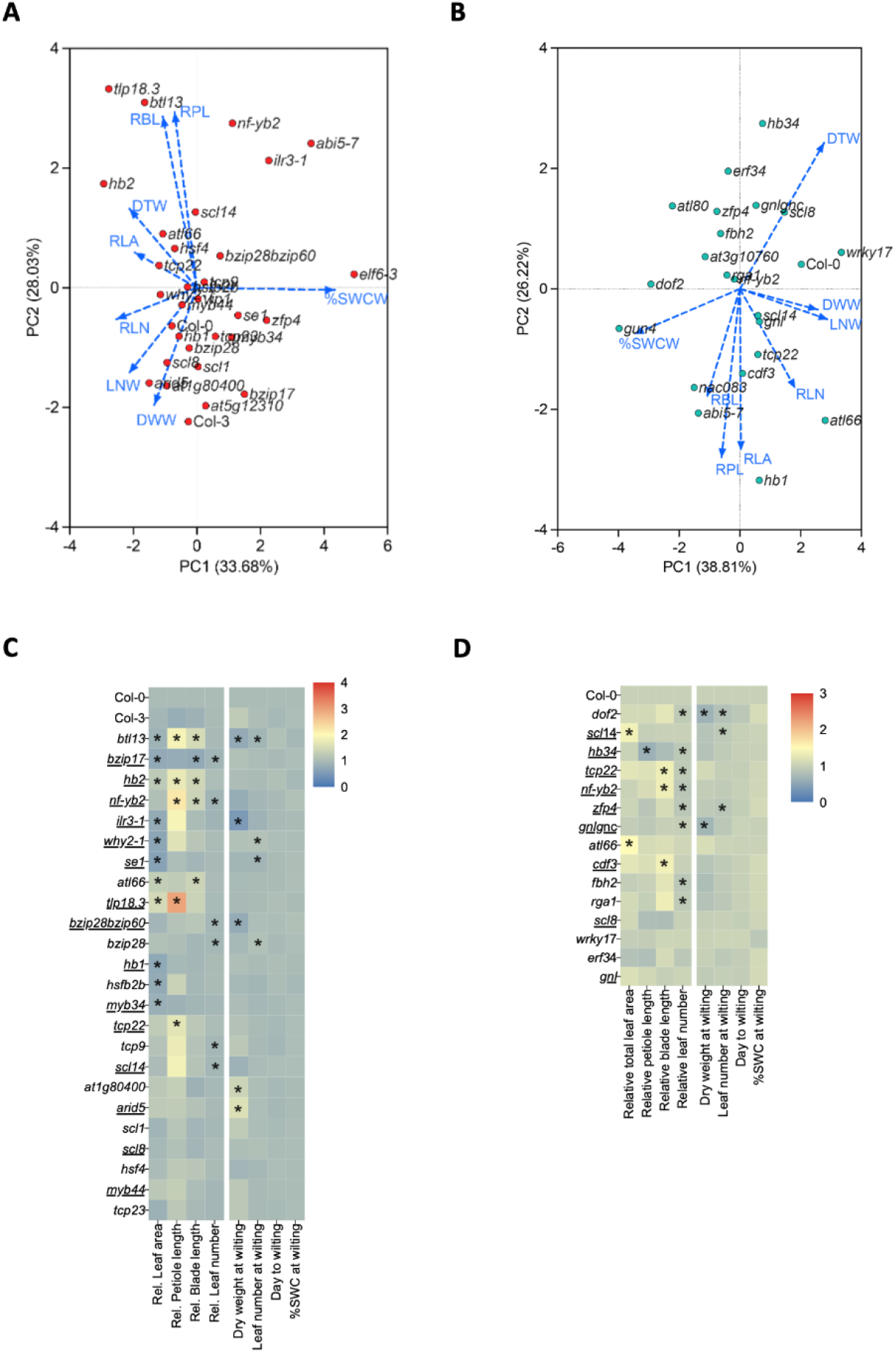
Plant growth, development and wilting trait analyses of selected mutants under combined or sequential stresses. **(A, B)** Principal component analysis (PCA) of all measured traits of all tested mutants and corresponding wild-type plants subjected to HTD (A) or PSD (B). Visualized are the distributions of mutants and wild-type plants indicated by symbols and correlations between measured traits by arrows. Abbreviations of traits: RLA: relative total leaf area, RPL: relative petiole length, RBL: relative blade length, RLN: relative leaf number, DWW: dry weight at wilting, LNW: leaf number at wilting, DTW: days to wilting, %SWCW: percentage of soil water content (%SWC) at wilting. **(C, D)** Heatmaps indicating phenotypic traits of mutants without early wilting phenotypes (opposed to those indicated in Fig. 3 and Fig. 4) along with the corresponding wild-type plants (Col-0 and Col-3) in response to HTD (C) or PSD (D). Relative values of total leaf area, petiole/blade length and leaf number were calculated by normalizing the data obtained at day 10 by that of day 0 (left four columns). Dry weight and leaf number at wilting, days to wilting and %SWC at wilting are indicated in the (right four panels). Abbreviated names of mutants and the wild-type plants are indicated per row. Color scales indicate relative values of measured traits (relative to the corresponding wild-type plants, with Col-3 relative to Col-0). Underlined mutants indicate confirmed knockout mutants of the corresponding genes based on previous studies (Supplemental Tables S6 and S7). Asterisks represent significant differences between mutant and the corresponding wild-type plants for the particular trait (Supplemental Table S8 and S9) (p < 0.05, unpaired t-test), n=8-12. For treatment abbreviations and used colors, see legend of Figure 1.

#### High temperature and drought

The *tlp18.3* and *bca2a zinc finger atl13* (*btl13*) mutants, correlated with leaf lengthening traits under HTD treatment and displayed significantly increased (young) leaf lengthening compared to WT (Col-0 or Col-3) (Fig. 6, A and C; Supplemental Table S8). For example, *tlp18.3* recorded a relative petiole length that was approximately 3-fold higher than Col-0 (Fig. 6C; Supplemental Table S8). A similar trend was observed for *homeobox 2* (*hb2*), *nf-yb2*, *arabidopsis tóxicos en levadura* (*atl66*) and *teosinte branched1* /*cycloidea*/*proliferating cell factor 22* (*tcp22*) mutants, exhibiting enhanced relative petiole and/or blade elongation (RPL and RBL) under HTD (Fig. 6, A and C; Supplemental Table S8). The mutants *btl13*, *why2-1* and *serrate* (*se-1*) showed reduced total leaf area and leaf number at wilting compared to the WT (Fig. 6C; Supplemental Table S8).

The basic region/leucine zipper motif (bZIP) TFs, bZIP17 and bZIP28 proteins have been implicated in heat stress responses (Gao et al., 2022; Kataoka et al., 2017). Compared to the WT, *bzip17* displayed significantly reduced total rosette area, blade lengthening and leaf formation under HTD, while *bzip28* had reduced leaf number (Fig. 6C; Supplemental Table S8). The *bzip28 bzip60* double mutant (Samperna et al., 2021), showed a significant delay in both leaf formation and dry weight accumulation (Fig. 6C; Supplemental Table S8). *ilr3-1* mutant plants exhibited strongly enhanced hyponastic growth when grown at elevated ambient temperature (27 °C), whereas additional drought during high temperature (HTD) did not further alter this phenotype (Supplemental Fig. S24, A and B). Next to these observations, several mutants displayed significant changes in only one of the measured traits (Fig. 6C; Supplemental Table S8). Notably, *ilr3-1* phenotypically resembled *abi5-7* (Fig. 6, A and C).

#### Post-submergence and drought

The *DNA binding with one finger 2* (*dof2*) mutant showed reduced leaf formation rate and dry weight accumulation in PSD (Fig. 6D; Supplemental Table S9). The *scl14* mutant showed increased total leaf area, whereas leaf number at wilting was reduced (Fig. 6D; Supplemental Table S9). Interestingly, *hb34*, which displayed decreased petiole length and leaf formation rate, exhibited a typical leaf-rolling phenotype in PSD, but not under C (Fig. 6D; Supplemental Fig. S25, A and B). Additionally, *tcp22* and *nf-yb2* mutants had significantly longer leaf blades but reduced leaf initiation rates (Fig. 6D; Supplemental Table S9). The *zfp4* mutant only showed a lag in leaf formation in PSD (Fig. 6D; Supplemental Table S9). The single *gnl* mutant (Richter et al., 2013) had no clear phenotype (Fig. 6D, Supplemental Table S9). However, in combination with a knockout of the *GNL* homolog, *GATA NITRATE-INDUCIBLE CARBON-METABOLISM INVOLVED* (*GNC*) (*gnlgnc*) reduced dry weight and impaired leaf initiation were noted (Fig. 6D; Supplemental Table S9). Additionally, both *atl66* and *cycling dof factor 3* (*cdf3*) mutants had larger leaves (Fig. 6D; Supplemental Table S9). The *flowering bhlh 2* (*@h2*) and *repressor of ga1-3 1* (*rga1*) mutants showed a significant reduction in relative leaf number compared to the WT when exposed to PSD (Fig. 6D; Supplemental Table S9).

## Discussion

Co-occurring abiotic stresses often have distinct effects on plants and elicit different acclimation responses compared to individual stresses (Choudhury et al., 2017; Nadeem et al., 2023; Zandalinas and Mittler, 2022; Zhang and Sonnewald, 2017). In addition, (combined) abiotic stresses encountered in natural or agricultural conditions are often at a gradual or sublethal severity, and hence relatively mild, compared to those reported in experimental laboratory studies (Morales et al., 2022; Zhang et al., 2020)

The experimental system that we present in this work departed from standard protocols of abrupt stress exposures. Arabidopsis plants were exposed to sublethal single and combined stresses, that typify weather events exacerbated by climate change. Our comprehensive dataset reveals a suite of unique phenotypic and molecular signatures of plant responses that emerge only when these sub-lethal stresses are applied sequentially or simultaneously (Fig. 1, C-E, Fig. 2, A and D; Supplemental Figs. S5, S7, S8, S10 S11 and S12). These responses are also highly dependent on the duration and sequence of stress imposition (Guo et al., 2021). For instance, many studies have explored the combined effects of heat and drought, but with varying results. For example, a predominant drought signature was noted on plant growth and development when heat and drought were applied concurrently (Zhou et al., 2017). In addition, in wheat (*Triticum aestivum*), episodes of prolonged drought in combination with heat waves exacerbate biomass reduction and loss of grain yield, when compared to individually applied drought or heat (Perdomo et al., 2015; Pradhan et al., 2012; Tricker et al., 2018). This is attributed to the negative interactions between heat and drought, with the effects becoming additive when combined (Suzuki et al., 2014). Additionally, a meta-analysis using >120 published cases studying crop responses to combined heat and drought revealed that it caused on average twice the decrease in relative yield (to control) compared to exposure to heat stress alone (Cohen et al., 2021). In apparent contrast, in our study, HT had a dominant effect over D on plant rosette growth and leaf development (Fig. 1, B-D; Supplemental Figs. S5 and S6). This is likely because in our experiments HT was applied from germination onwards, well before D was applied.

Submergence significantly hampered plant growth but did not cause plant death (Fig. 1, B-D; Supplemental Figs. S8, S9 and S10). This is likely because our flooding treatment occurred under the regular day/night regime, which differs from studies where lethality was noted using submergence in darkness (Van Veen et al., 2013; Vashisht et al., 2011). Plants were able to steadily recover if optimal irrigation was applied following de-submergence (Fig. 1, B-D; Supplemental Figs. S8, S9 and S10). However, when experiencing drought following de-submergence (PSD), leaf lengthening, rosette area increase and dry mass accumulation were significantly impacted (Fig. 1, B-D; Supplemental Figs. S8, S9 and S10). This is in line with Luo et al. (2009) who highlighted the essential role of photosynthetic acclimation during submergence recovery (Luo et al., 2009). As drought can be a major impediment to photosynthesis (Chaves et al., 2009; Cornic, 2000), the observed inhibition of growth and development could be explained by hampered photosynthetic acclimation. Interestingly, phenotypic differences between PS and PSD were greater than those between C and D after 10 days (Supplemental Fig. S10). This implies that PS exerted a negative synergic effect on D in regulating plant growth and development in the PSD treatment.

Although both HT and PS interact with D and consequently elicit distinct effects on plant phenotypes under the combined stress conditions, HT and PS seemingly affect physiological responsiveness to HTD and PSD to different degrees. For example, when combined with HT, drought-promoted leaf temperature increase occurred substantially earlier than under individual drought conditions (Supplemental Fig. S7, B and C). In contrast, a submergence pre-treatment (S) did not accelerate drought-induced leaf temperature elevation (Supplemental Fig. S11, B and C). Furthermore, stomatal conductance decreased 5 days after HTD (Fig. 1E), but not during 5-day PSD (Fig. 1E; Supplemental Fig. S12B). A likely explanation is that stomata are highly susceptible to high temperature, resulting in a pronounced stomatal response when combined with drought (De Smet et al., 2023).

Drought can have a major impact on the transcriptome (Bouzid et al., 2019; Chen et al., 2021). However, while our plants clearly experienced drought (Fig.1, B-D; Supplemental Fig.S1 and S2), D did not elicit significant transcriptomic changes (3 DEGs, Fig.2, A and D) nor evoked strong effects on phenotypic traits shortly after stress initiation (Fig. 1, B-D; Supplemental Figs. S5, S6, S8, S9 and S10). This is consistent with a study by Prasch & Sonnewald in which only 41 DEGs were identified after 5-day progressive drought treatment (Prasch and Sonnewald, 2013). The relatively mild drought intensity likely explains the limited number of DEGs (Clauw et al., 2015). In addition, the young leaves that were sampled may be less responsive to drought than for instance roots tissues, as plants might evoke protective mechanisms to prioritize stress protection of younger leaves and meristematic tissues (Rankenberg et al., 2024, 2021; Verelst et al., 2013).

In line with observed unique responses at the phenotypic level, combinatorial stresses imposed distinct and more profound effects on the transcriptome compared to single stress application (Fig. 2; Supplemental Fig. S14). For both stress combinations we observed a considerable number of unique multi-stress specific DEGs. Notably, the majority of PSD-triggered DEGs were not affected by PS nor D (Fig. 2, D and E). This difference in transcriptome reconfiguration is consistent with the concept that prior exposure to one stress can have consequences for the response to the second stress (Rejeb et al., 2014).

Our RNA-seq dataset permitted the identification of genes putatively contributing to morpho-physiological responses to combinatorial and/or individual stresses. To identify these candidate genes, we focused on upregulated TFs and regulatory hubs identified from constructed GRNs (Fig. 2, G and H; Supplemental Fig. S19; Supplemental Table S5). 39 candidate genes identified in this way were experimentally validated to probe their functions in combinatorial stress acclimation. The aim was to unveil genetic effects based on testing a broad range of candidate factors, rather than focusing on deciphering the precise molecular mechanisms underlying selected genes. This approach admittedly neglects downregulated genes, which might also play important roles in mediating responses to combined and/or sequential stresses and are worthwhile to test in future experiments.

We identified *ELF6* as a negative regulator of phenotypic acclimation to HTD, while *ATL80* is a positive regulator for PSD (Fig. 3, B and D, Figs. 4 and 5; Supplemental Figs. S20 and S22). *ELF6* and its close homolog *RELATIVE OF EARLY FLOWERING 6* (*REF6*), function as histone H3 at Lys27 trimethylation (H3K27me3) demethylases in Arabidopsis (Crevillén et al., 2014; He et al., 2022; Lu et al., 2011). H3K27me3 is crucial for the regulation of plant thermomorphogenesis and heat stress memory (Casal and Balasubramanian, 2019; Perrella et al., 2022; Yamaguchi et al., 2021). Therefore, the involvement of *ELF6* in regulating plant growth and development under HTD may connect to H3K27me3-mediated high temperature responses. ATL80 is a plasma membrane-localized RING E3 ubiquitin ligase that negatively regulates cold stress response and phosphate mobilization (Suh and Kim, 2015). ATL80 was recently found to potentially target components within the retrograde signaling pathways for degradation (Méndez-Gómez et al., 2024). Additionally, we identified regulators such as *ZFP4* and *GUN4*, as factors in controlling plant wilting under HTD or PSD. *ZFP4* and *GUN4* appear to participate in rosette expansion and/or dry weight accumulation under individually applied stresses such as D, HT and PS, but not combinatorial stresses (Fig. 3, B and D, Figs. 4 and 5; Supplemental Figs. S20 and S22). Further investigations are needed to decipher their precise roles in controlling (combinatorial) stress acclimation.

## Conclusions

From the observed diversity in affected phenotypic traits across the tested mutants, treatments and timepoints (Fig 3, B and D, Fig. 6), we conclude that different aspects of acclimation strategies of plants to deal with HTD or PSD, are coordinated by a multitude of genetic factors. These factors together determine certain phenotypic outcomes depending on developmental (st)age and relative stress levels. For example, in response to HTD, TFs such as TLP18.3, BTL13, TCP22 and HB2 are likely involved in orchestrating leaf elongation, possibly as negative regulators, as their corresponding mutant lines displayed induced relative leaf lengthening (Fig. 6, A and C). Upon PSD treatment, HB1 and CDF3 seemingly inhibit the lengthening of petiole and blade (Fig. 6, B and D).

Altogether, our work contributes to a better understanding of plant response to sublethal combinatorial stresses. Generated insights can contribute to the establishment of mechanistic models for predicting trait responsiveness under the given stress condition. In addition, it should be investigated how sublethal combinatorial stresses affect productivity in agronomical crops, and whether the multitude of identified multi-stress regulators also contribute to resilience of commercial crops. Such investigations are critical considering the increased co-occurrence of these stresses in the context of climate change. This work has therefore the potential to contribute to breeding and/or engineering of field crops that maintain optimal yields in future climate conditions.

## Materials and methods

### Plant materials, growth conditions and stress treatments

Arabidopsis Col-0 seeds were sown on Primasta potting soil (Asten, The Netherlands) and stratified 4 days in darkness. Plants were grown in MD1400 (Snijders, The Netherlands) climate chambers under 8h photoperiod, 21 °C or 27 °C (day and night), 130-150 μmol m^-2^ s^-1^ PAR with fluorescence tube or LED lightening and 70% RH. Plants that reached the 2 true-leaf stage were transplanted to Jiffy 7c coconut pellet substrate (Jiffy Products International BV, Zwijndrecht, The Netherlands) that were pre-soaked in water and 50 mL Hoagland solution (Millenaar et al., 2005) (saturated weight = 250±20 g), contained in 9×9 cm pots for stability. Additional Hoagland solution was supplied 2, 5 and 7 days after transplantation. Every second day, water was added and pots were randomized to account for position effects.

Plants containing 10 true leaves (LS10) were subjected to sublethal stress conditions two hours after the photoperiod started or remained in control conditions (C/HT). Progressive drought (HTD/D) consisted of withholding watering. Drought severity, expressed as relative soil water content (%SWC), refers to the ratio between the measured pellet weight and weight at full field capacity (Supplemental Fig. S1, A and B). Plants subjected to submergence (PS and PSD) were pre-grown and treated at 21 °C. One day before treatment initiation, plastic tubs (54 x 27 x 37 cm) were disinfected with a chlorine tablet (Diversey Inc., Racine, USA) and thereafter rinsed. The tubs were filled with deionized water and allowed to equilibrate to the climate room’s temperature. Plants were completely submerged (S) for 5 days while receiving the regular day/night treatment. Thereafter, the pellets containing the plant were covered by absorbent papers to remove excess water until the weight was comparable to non-submerged control (C) and were then either subjected to the regular regime of irrigation (PS), or to progressive drought conditions by withholding watering (PSD).

### Morphological analyses

To assess leaf angles the two most hyponastic intermediate-aged leaves were imaged after placement perpendicular to a camera. Plant shoots were thereafter detached, flattened using transparent paper, and imaged from the top. ImageJ software (National Institutes of Health, USA) was used to determine leaf traits. For hyponasty, leaf angle relative to the horizontal was measured by taking the average angle of the captured opposing leaves.

To quantify dry weight, plants were dried in an oven set at 80 °C and weighed on an analytic balance (Sartorius BP221S, Göttingen, Germany). To quantify leaf initiation (rate), the number of all visible leaves (excluding cotyledons), was scored daily. Primary root length was determined using a ruler, after the pellet substrate was rinsed from the roots with tap water.

### Physiological trait analyses

Chlorophyll content was determined using a CCM-300 Chlorophyll Content Meter (Opti-Sciences lnc., Hudson, USA). After calibration, the detector was placed at 5 mm distance from the leaf and chlorophyll content was calculated based on the fluorescence intensity. For destructive biochemical detection, chlorophyll was extracted from detached leaves using 96% DMSO. Absorption was determined at 664, 647 and 750 nm wavelengths using a spectrophotometer plate reader (Synergy HT Multi-Detection Microplate Reader; BioTek Instruments Inc., USA). Dry weight of the remaining plant material was used to calculate relative chlorophyll a, b, and total chlorophyll content.

Relative water content was calculated as (Fresh weight–dry weight)/(turgor weight–dry weight))x100%. To this aim, fresh weight of excised rosettes was determined using an analytic balance (Sartorius BP221S, Göttingen, Germany). The rosettes were then immediately allowed to saturate with water in petri dishes (24 hours in darkness). Excess water was removed and turgor weight of plants was measured. Plants were then dried at 80°C and dry weight was recorded.

Stomatal conductance was measured of the abaxial leaf side using an SC-1 Leaf Porometer (Decagon Devices, Inc., Pullman, USA) equilibrated to ambient temperature.

Leaf surface temperatures were recorded using a FLIR A655sc LWIR thermal imaging camera (Teledyne FLIR, USA) with a 13.1 mm FoV 45° x 33.7° hawkeye IR lens. Thermal images were captured every 15 minutes using FLIR ResearchIR Max 4 software (Teledyne FLIR, USA) from the start of the stress treatments till wilting occurred. Temperatures of 6 leaves per plant were measured at ZT = 0, 6, 12, 18 h and every 4^th^ image, equaling every 24 hours, the regions of interest was adjusted to correct for leaf growth and movement.

To determine Malondialdehyde (MDA) content, detached rosettes were grinded in ethanol after recording fresh weight and cold-centrifuged. Part of the supernatant was mixed with an equal volume of TCA/TBA solution (0.65% (w/v) thiobarbituric acid in 20% (w/v) trichloroacetic acid). The mixture was shaking-incubated at 95°C for 30 minutes and cooled in an ice bath. After cold-centrifugation, absorbance was measured using a spectrophotometer plate reader (Synergy HT Multi-Detection Microplate Reader; BioTek Instruments Inc., USA) at 532 and 600 nm. MDA concentration was calculated and normalized by fresh weight: MDA (nmol/mL) = ((A532-A600)/155000)x1000000

For quantification of ion leakage, rosettes were detached, placed in a tube containing deionized water and shaken by rolling (1 hour at RT). Initial conductivity was measured using a pre-calibrated EC-33 conductivity meter (HORIBA, Japan). The samples were then incubated at 95°C for 30 minutes and cooled down to RT, after which final conductivity was measured. Ion leakage was calculated as: Ion leakage=(Initial conductivity/final conductivity)x100%.

### Transcriptomics sample preparation and analysis

The 7^th^-10^th^ leaves counted from the earliest emerged true leaves at LS10 were flash-frozen in liquid nitrogen and kept at -80 °C. Samples were grinded using a cryogenic grinding mill (Retsch, Haan, Germany) and plant total RNA was extracted using the RNeasy kit (Qiagen, Germany) following the manufacturers protocol.

RNA quality control, library construction and sequencing were performed by Macrogen (Amsterdam, The Netherlands). RNA integrity and purity were determined using an Agilent Technologies 2100 Bioanalyzer (Agilent Technologies, Palo Alto, USA) and only samples with Integrity Number of 7 or above were used for library construction by the TruSeq stranded mRNA protocol (Illumina, USA). Libraries were sequenced using an Illumina Novaseq6000 sequencer (Illumina, USA) with 150 bp pair-end reading. Generated datasets were deposited at the NCBI Gene Expression Omnibus (Project ID: PRJNA863409).

Data processing (trimming, read filtering and mapping) and differential gene expression analysis were conducted as previously described (Verhoeven et al., 2023). Two samples with a total mapped Arabidopsis read count below 7 million due to *ArLV1* infection were removed. For the determination of *ArLV1* effects on plant transcriptome, control (C) samples at timepoint 5 days were selected, as three of these samples contained relatively few reads mapping to *ArLV1* (0.01-9.56%) and four samples contained high amounts of *ArLV1* (78.94-90.08%). Only for this analysis, the complete dataset including the two low-quality samples was used.

Statistical analysis was done in R using the log_2_-normalized TPM values in linear models ran for 5 and 10 days timepoints separately, with different stress treatments as variables compared to the control treatment at the same timepoint. Obtained significances were corrected using a Benjamini-Hochberg adjustment (provided by the *prcomp* function). Expressed genes were visualized by ggplot2 (Wickham, 2009).

*k-means* clustering was used to arrange the significantly regulated genes in clusters with similar expression patterns. The optimal number of clusters was determined visually by plotting the within-cluster sums of squares and the average silhouette with the fviz_nbclust function from the factoextra package in R (Lê et al., 2008). Groups with a high variation in expression between the stress treatments were chosen for additional enrichment analysis.

For the GO enrichment analysis, Metascape (Metascape, http://Metascape.org/) (Zhou et al., 2019) was used with settings of p value cutoff = 0.01, minimal number of overlapping genes = 3 and a minimal enrichment value = 1.5.

The locus IDs of Arabidopsis were inputted into Arabidopsis transcription factor database (AtTFDB, https://agris-knowledgebase.org/AtTFDB/) and TFs were automatically identified by annotating the candidates to TFs from 50 different TF families. Candidate gene IDs were first imported into TF2Network (Kulkarni et al., 2018) to search for promoter binding sites and thereafter putative upstream TFs were identified. These TFs were then analyzed for co-expression and protein-DNA interactions with the predicted targets and the networks thereafter generated using Cytoscape (Shannon et al., 2003).

### Confirmation of T-DNA insertion lines

gDNA was extracted from leaves using the Direct PCR-Phire and Phusion kit (Thermo Scientific, lnc, USA) following the manufacturers protocol. Presence of T-DNA insertions and homozygosity were determined by PCR (primers see: Supplemental Tables S6 and S7). We were unable to obtain homozygous lines for 12 genes, mainly because of issues with seed germination of obtained seed stocks.

### Statistical analysis

If not explicitly mentioned otherwise, figures were generated using GraphPad Prism 9 (GraphPad Software, La Jolla, USA) or Biorender.com. Statistical analyses were performed using GraphPad Prism 9 or R software. Significance was calculated using an alpha of 0.05.

## Acknowledgements

We thank Rens Voesenek and Sjef Smeekens for their support in the early stages of the project. Evelien Stouten, Alejandro Morales and Basten Snoek are thanked for their technical support and advice.

## Author contributions

ZJ, AV, RS, MvZ, designed the research; ZJ, AV, YL, RG performed research. ZJ, AV, RS, MvZ, analyzed data; ZJ, RS, MvZ wrote the paper.

## Accession numbers

Generated sequence datasets from this article were deposited at the NCBI Gene Expression Omnibus (Project ID: PRJNA863409).

