## Supplemental data for "Deciphering *Arabidopsis thaliana* Acclimation to Sublethal Combined Abiotic Stresses"

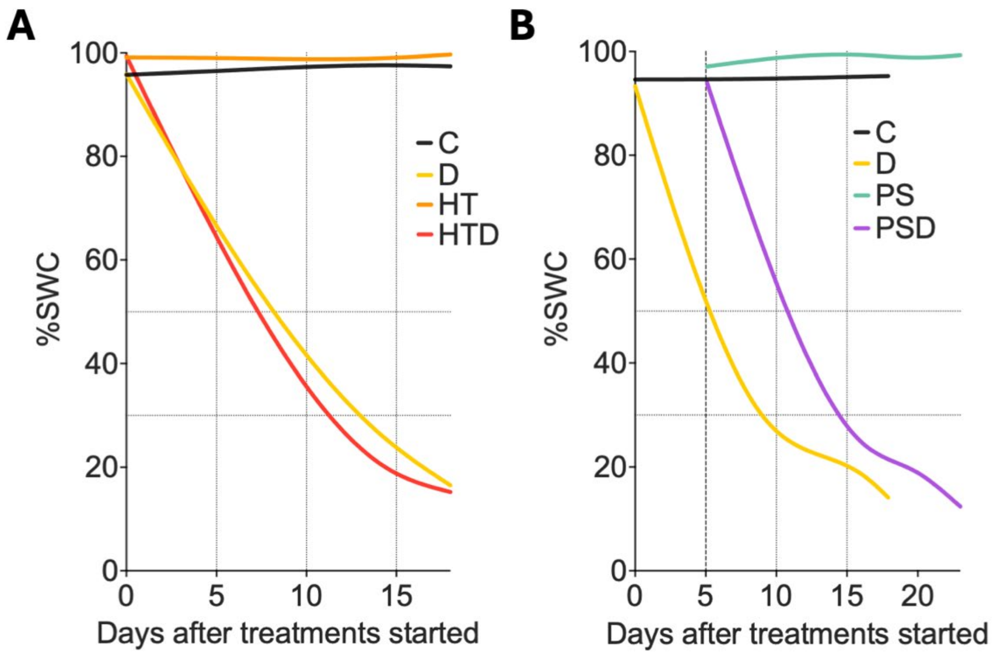


**Supplemental Figure S1.** Progressive decline in soil water content (%SWC) during combined high temperature and drought **(A)** and post submergence followed by drought **(B)**, and the composite single stresses and control. (A) n = 3-18. (B) n = 8-34. Abbreviations; C: control (black lines), D: drought (yellow), HT: high temperature (orange), HTD: high temperature & drought (red), S/PS: 5-day submergence / (post-)submergence & recovery (blue), PSD: (post-)submergence & drought (purple).

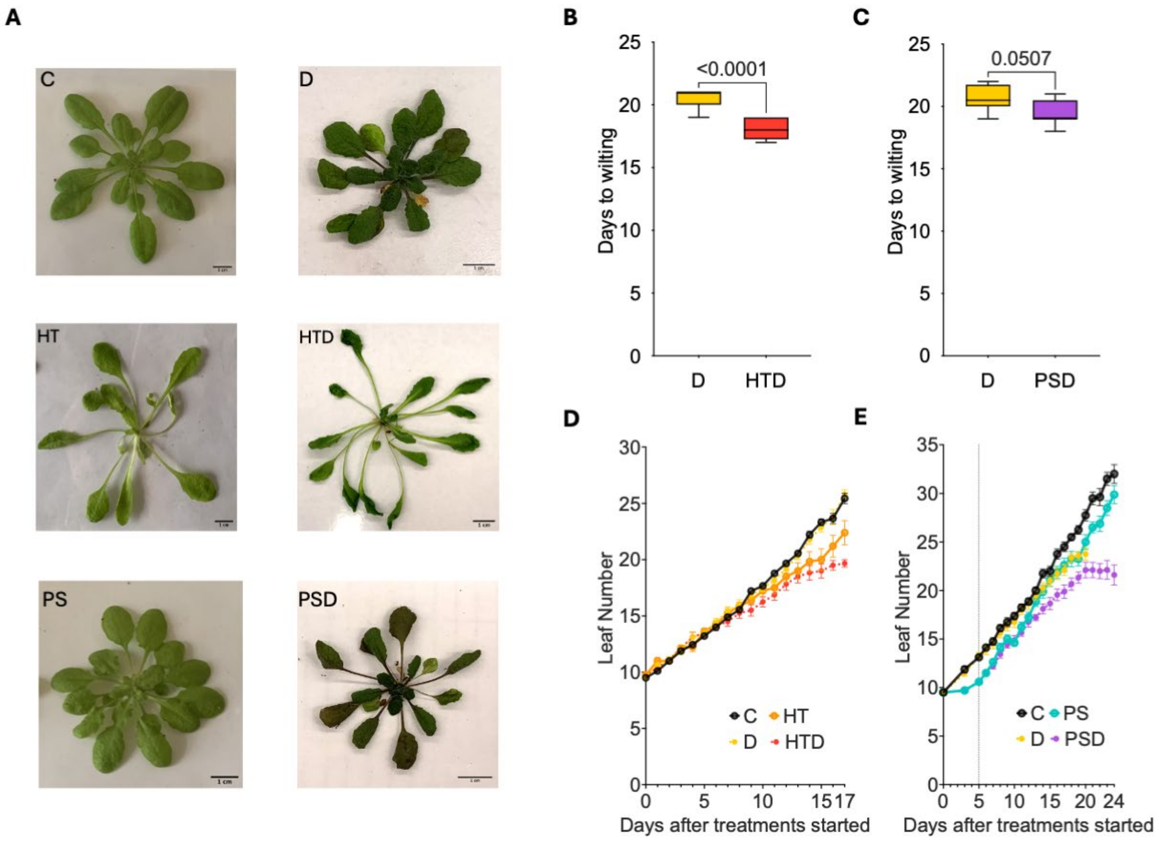


**Supplemental Figure S2.** Effect of combined and sequential stresses on leaf initiation and wilting. **(A)** Representative images showing Arabidopsis Col-0 rosettes at the wilting stage (right column; non-turgid) after progressive drought and the relative controls (left column; turgid non-wilted plants) at control temperature (21 °C, C and D, upper row), high temperature (27 °C) combined with drought (HT and HTD, middle row) and post submergence followed by drought (PS and PSD, bottom row). **(B, C)** Number of days until wilting occurred in plants subjected to (B) drought at control temperature (21 °C; D) and at high temperature (27 °C; HTD) and (C) drought at control temperature (21°C; D) and post-submergence followed by drought (PSD). Days were counted after water was withdrawn. (B) n = 8-10. (C) n = 8-9. Boxes indicate boundaries of the second and third quartiles (Q) of the data distribution. Black horizontal bars indicate median and whiskers Q1 and Q4 values within 1.5 times the interquartile range. Numbers above the bars indicate p values (unpaired t-test). **(D, E)** Leaf number of plants exposed to combined high temperature and drought (D) (HTD) or post-submergence followed by drought (E) (PSD) and the associated single stresses (HT, HTD, D, PS, PSD) and controls (C). Error bars indicate means ± SEM, (D) n = 6-18. (E) n = 5-33. The dashed vertical line in panel (E) indicates the moment plants were de-submerged. For treatment abbreviations and used colors, see legend of Supplemental Figure S1.

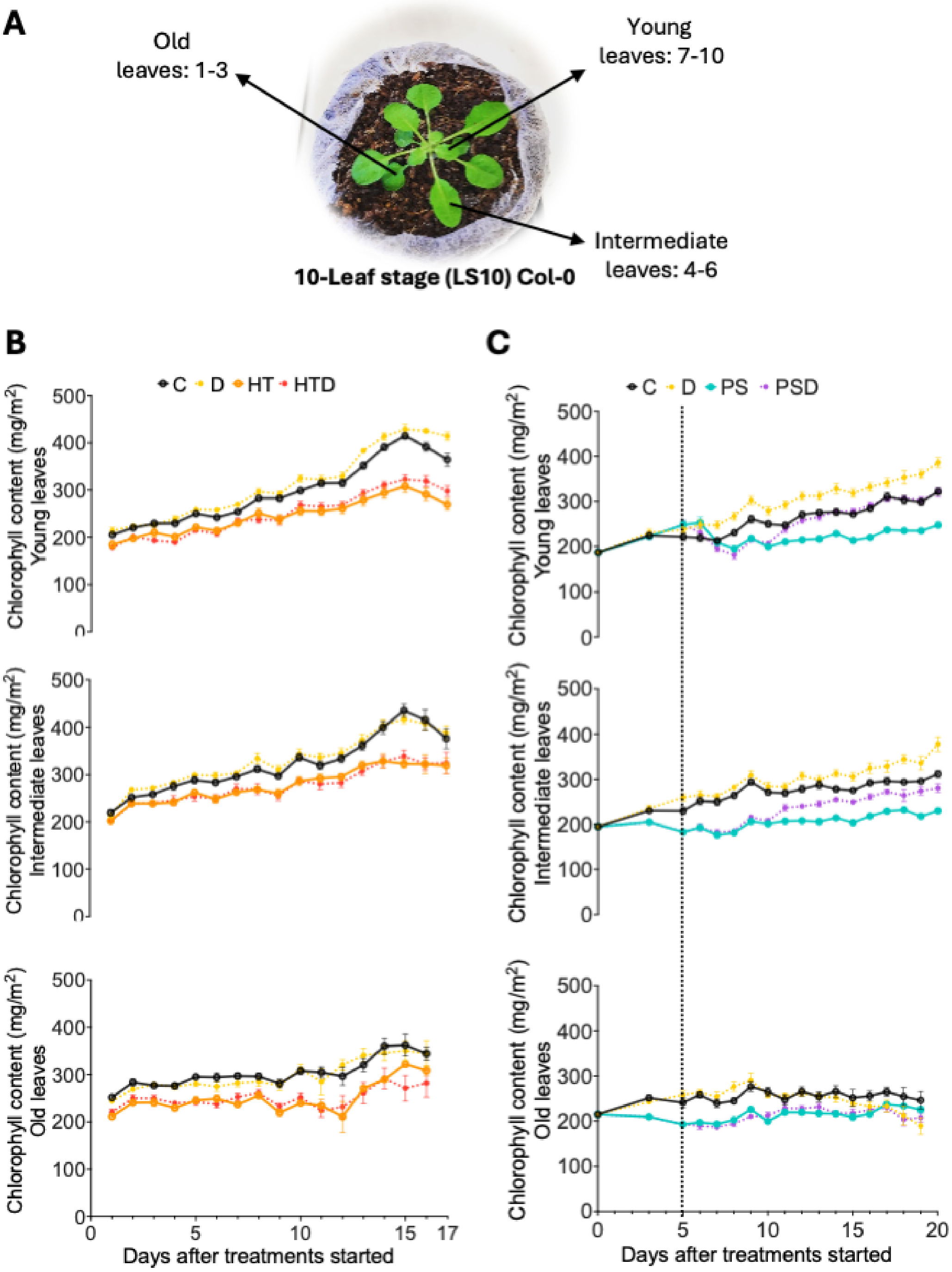


**Supplemental Figure S3.** Effect of combined and sequential stresses on chlorophyll content obtained with a CCM-300 chlorophyll content meter. **(A)** Representative image of a 10-leaf stage (LS10) Col-0 plant on Jiffy 7c coconut pellet growth substrate, with young, intermediate, and old leaves indicated. Associated leaf numbers are counted starting from the first true leaves (thus excluding the cotyledons). **(B, C)** Chlorophyll content of young (upper row), intermediate (middle row), and old leaves (lower row) exposed to high temperature & drought (HTD) (B) or post-submergence & drought (PSD) (C), as well as the associated single stresses (HT, PS, D). Error bars indicate means ± SEM, (A) n = n = 4-9. (B) n = n = 5-33. The dashed vertical line in panel (C) indicates the moment plants were de-submerged. For treatment abbreviations and used colors, see legend of Supplemental Figure S1.

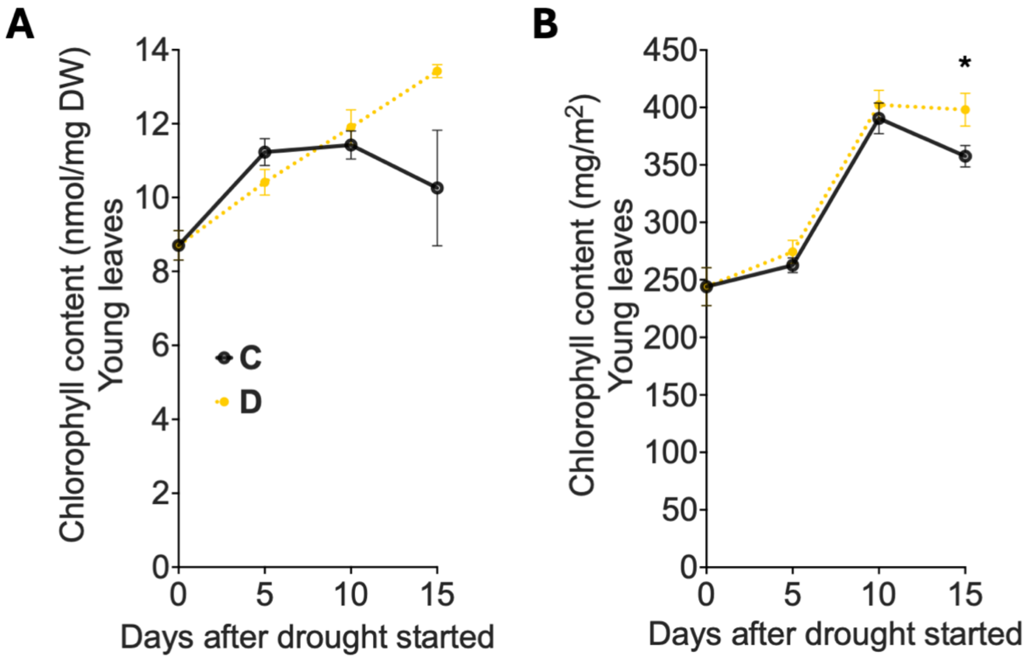


**Supplemental Figure S4.** Chlorophyll content of young leaves in control (C) and drought (D) conditions, obtained by destructive biochemical assay **(A)** and by using a CM-300 chlorophyll-meter **(B)**. Harvests and measurements were conducted at 0, 5, 10, and 15 days after drought started. Error bars indicate means ± SEM. n = n = 4-5. Asterisks represent significant differences between drought and control plants within the same time point ($p < 0.05$, unpaired t-test). For treatment abbreviations and used colors, see legend of Supplemental Figure S1.

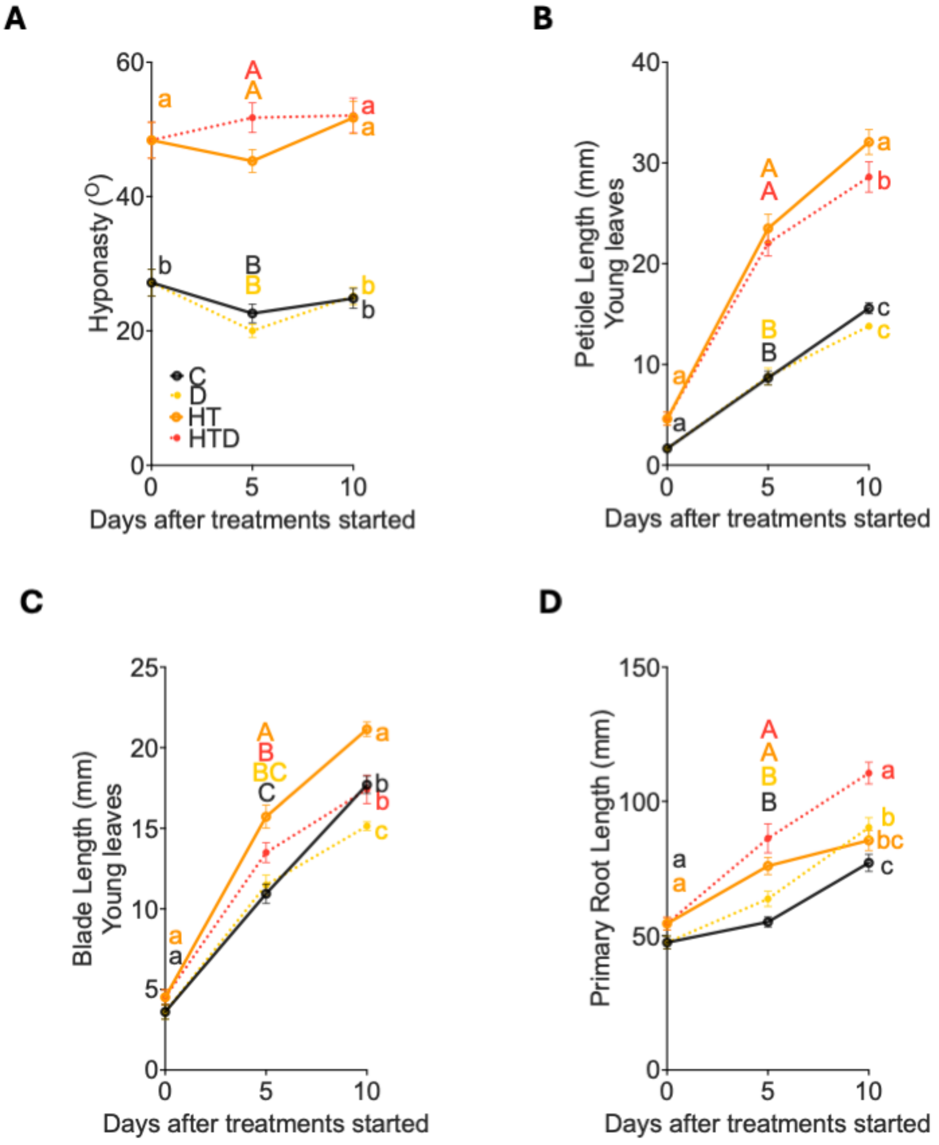


**Supplemental Figure S5.** Effects of combined high temperature and drought and the corresponding single stresses on morphological traits. **(A)** Average angles of the two most hyponastic leaves of individual plants, relative to the horizontal. n = n = 14-21. **(B, C)** Average length of petiole (B) and blade (C) of young leaves. n = 15-21. **(D)** Primary root length. n = 14-21. Error bars indicate means ± SEM. Letters denote significant differences between different treatments at the same time points ($p < 0.05$, 2-way ANOVA with Tukey's Post-hoc test). For treatment abbreviations and used colors, see legend of Supplemental Figure S1.

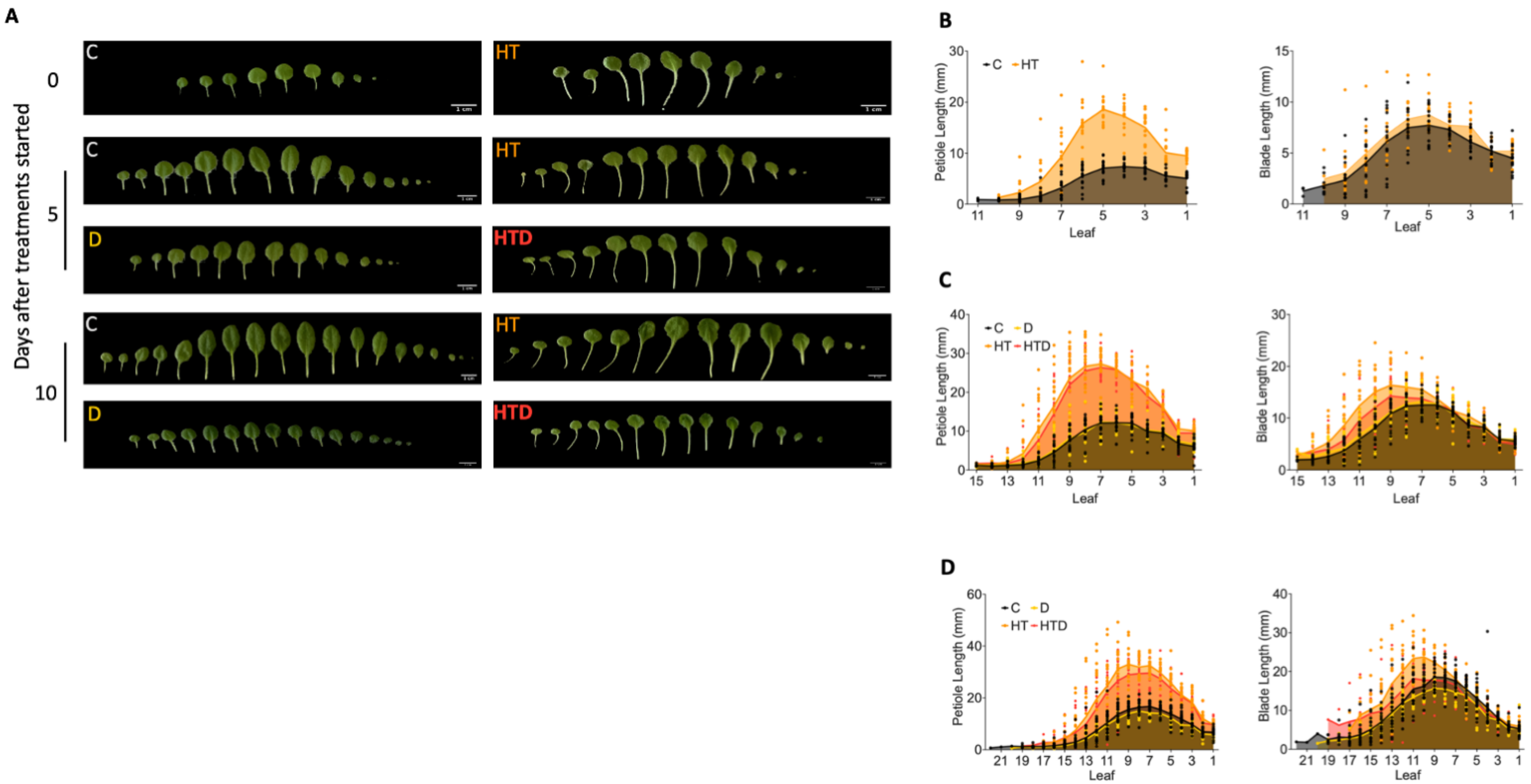


**Supplemental Figure S6.** Dynamics in plant petiole and blade length in response to combined high temperature and drought and the corresponding single stresses and controls. **(A)** Representative images of dissected leaves (ordered from old to young, with the cotyledons on the far right side) from plants subjected to single and combined stresses at 0, 5, and 10 days, counted from the start of stress treatments. Scale bars indicate 1 cm. **(B, C, D)** Petiole (left) and blade (right) lengths of all leaves from plants subjected to high temperature combined with drought and related single stresses and control at 0 (B), 5 (C) and 10 (D) days after the start of stress treatments. Dots and lines indicate individual plants and averaged data, respectively. n = 15-21. For treatment abbreviations and used colors, see legend of Supplemental Figure S1.

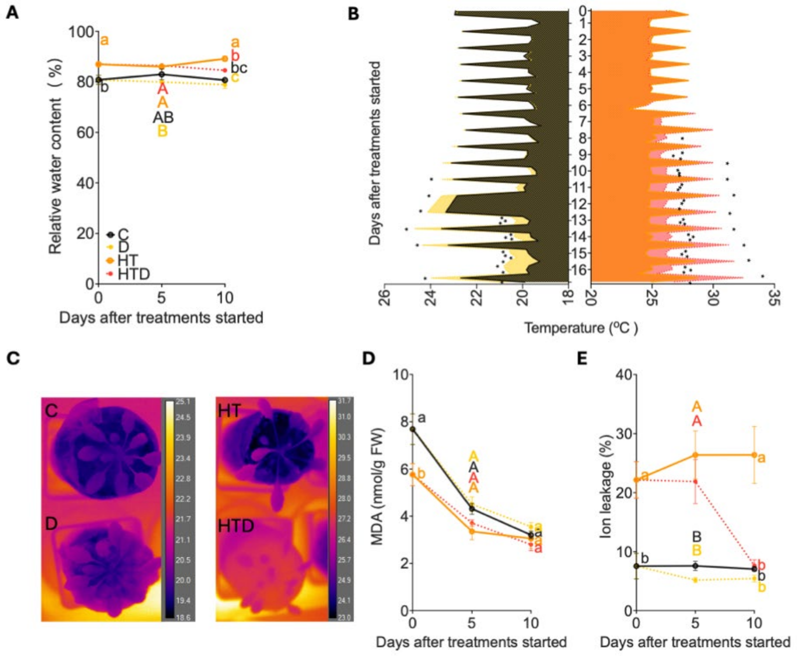


**Supplemental Figure S7.** Effects of combined high temperature and drought and corresponding single stresses on physiological traits. **(A)** Rosette water content relative to the maximal water content that the leaves can hold (100%, turgid water - dry weight). n = 8-11. **(B)** Dynamic changes in leaf surface temperature at 21 °C (C and D, left) and 27 °C (HT and HTD, right). Lines represent the average leaf temperature as measured every 6 hours (ZT = 0, 6, 12, and 18 h). n=3. Asterisks represent significant differences between measured leaf temperature between drought (D and HTD) and well-watered plants (C and HT) within the same timepoints (p < 0.05, One-way ANOVA with Tukey's Post-hoc test). Note that temperature fluctuates between the photoperiod (peak) and dark period. Data from three time points (ZT = 18 h on day 11 and ZT = 0 and 6 h on day 12 in 21 °C) were not recorded due to camera failure. **(C)** Representative thermal images of plants at 21 °C (C and D-treated, left) and 27 °C (HT-treated and HTD-treated, right) 10 days after the treatments started. **(D)** Rosette Malondialdehyde (MDA) content. n = 5-10. **(E)** Rosette ion leakage relative to the maximal electrolyte conductivity (100%). n = 6. (A, D, E) Error bars indicate means ± SEM. Letters denote significant differences between treatments within the same time points (p < 0.05, 2-way ANOVA with Tukey's Post-hoc test). For treatment abbreviations and used colors, see legend of Supplemental Figure S1.

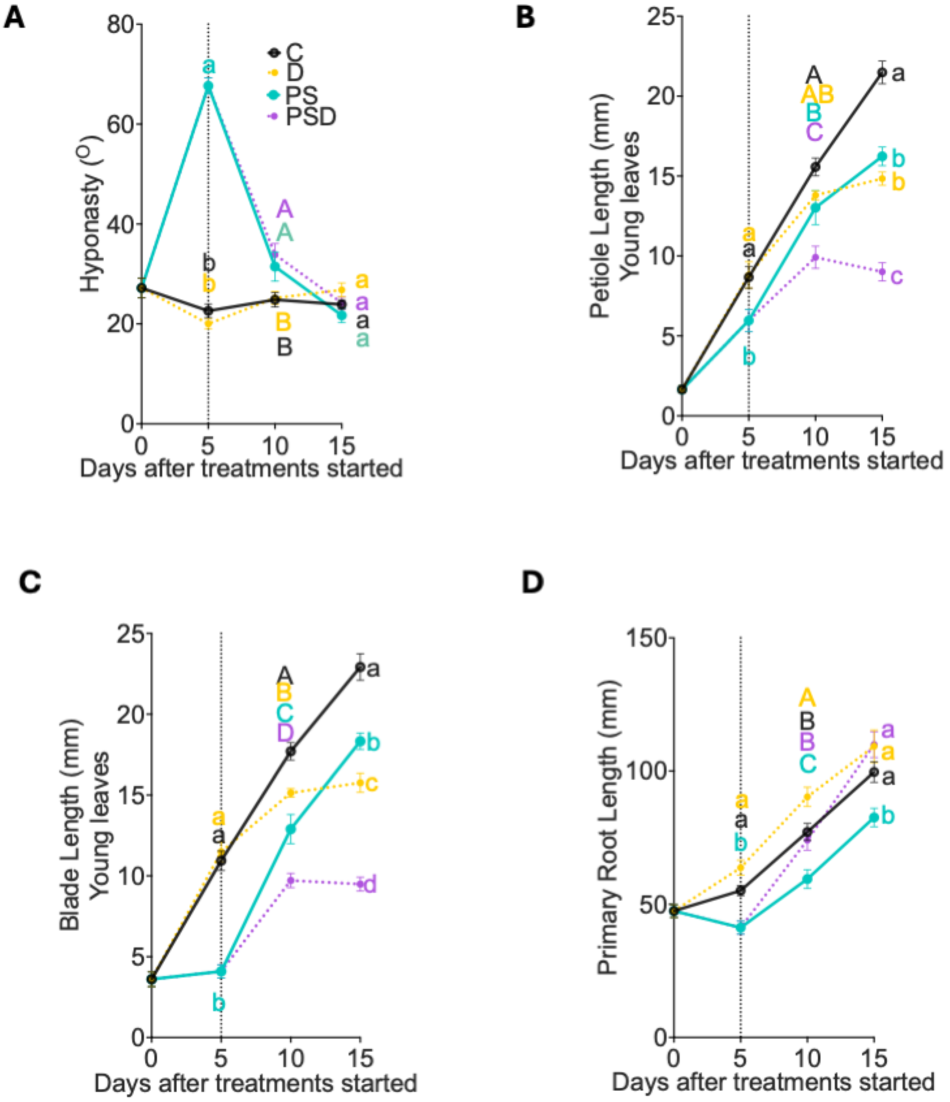


**Supplemental Figure S8.** Effects of sequential submergence and drought and the corresponding single stresses on various morphological traits. **(A)** Average angles of the two most hyponastic leaves of individual plants, relative to the horizontal. n = 14-21. **(B, C)** Average length of petiole (B) and blade (C) of young leaves. n = 15-21. **(D)** Primary root length. n = 14-21. Error bars indicate means ± SEM. Letters denote significant differences between different treatments at the same time points ($p < 0.05$, 2-way ANOVA with Tukey's Post-hoc test). For treatment abbreviations and used colors, see legend of Supplemental Figure S1.

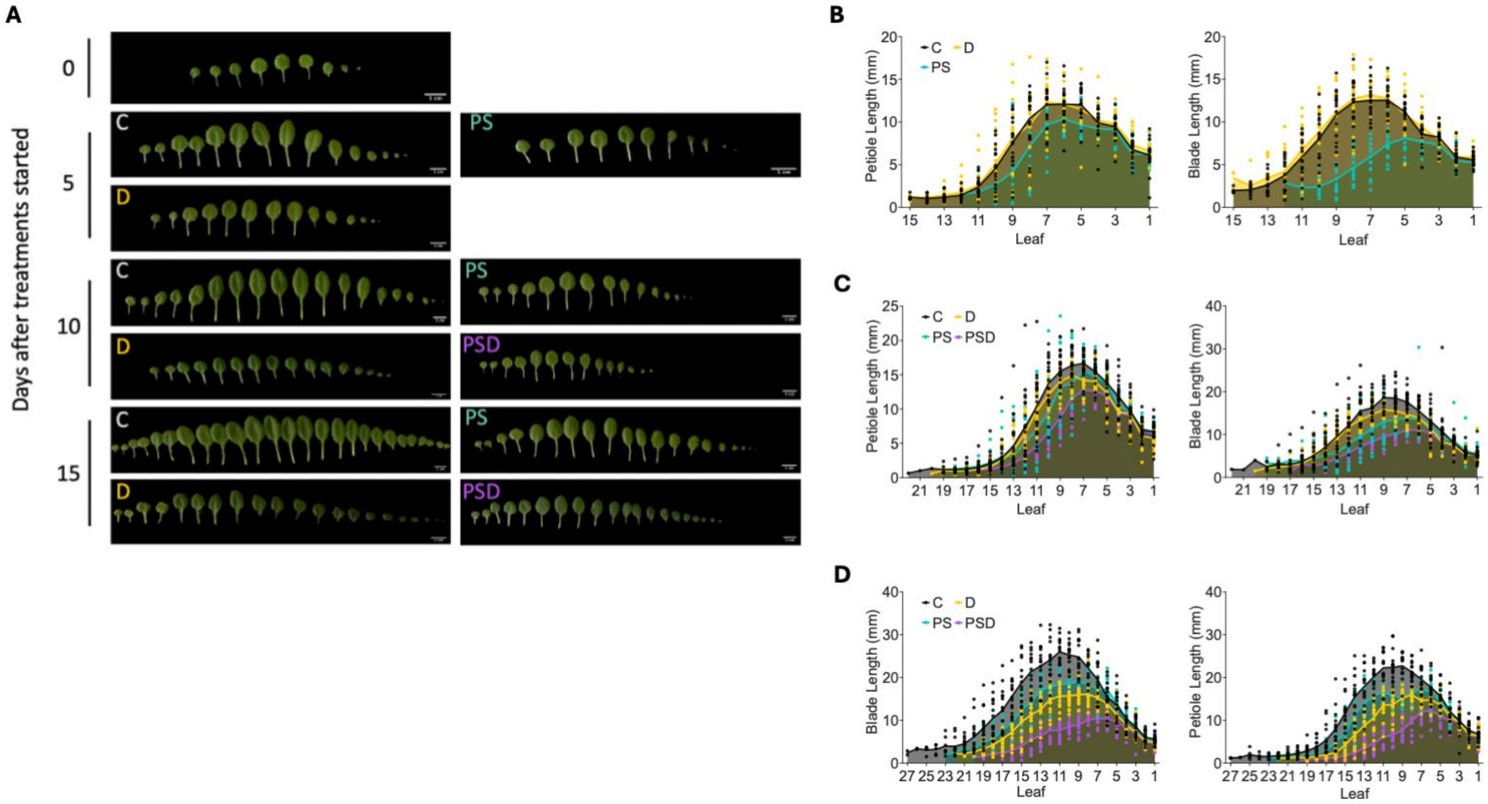


**Supplemental Figure S9.** Dynamics in plant petiole and blade length during post-submergence combined with drought and the relevant single stresses and controls. **(A)** Representative images of dissected leaves (ordered from old to young, with the cotyledons on the far right side) from plants subjected to single and sequential stresses at 0, 5, 10 and 15 (C and D) days, counted from the start of the stress treatments. Scale bars indicate 1 cm. (B, C, D) Petiole (left) and blade (right) lengths of all leaves from plants subjected to post-submergence followed by drought and related single stresses and control at 0 (B), 5 (C) and 10 (D) days after the start of de-submergence phase. Dots and lines indicate individual plants and averaged data, respectively. n = 15-21. For treatment abbreviations and used colors, see legend of Supplemental Figure S1.

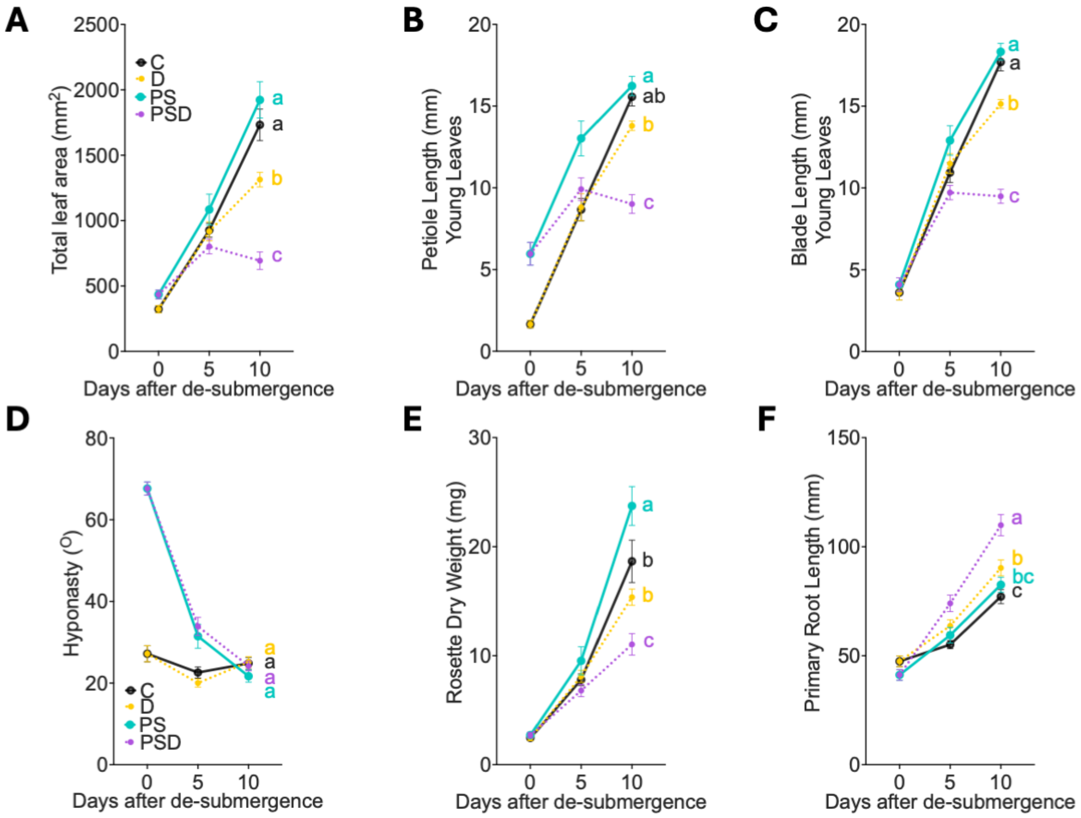


**Supplemental Figure S10.** Phenotypic traits as an effect of drought in presence or absence of prior submergence. **(A)** Total leaf area of the whole rosette. n = 15-20. **(B, C)** Averaged length of the petiole (B) and blade (C) of young leaves. n = 15-21. **(D)** Averaged angles of the 2 most hyponastic leaves of individual plants, relative to the horizontal. n = 14-21. **(E)** Rosette dry weight. n = 15-18. **(F)** Primary root length n = 14-21. Error bars indicate means ± SEM. Letters denote significant differences between different treatments the stress treatments ($p < 0.05$, 2-way ANOVA with Tukey's Post-hoc test). Data presented here is derived from Figure 1 and Supplemental Figure S8. For treatment abbreviations and used colors, see legend of Supplemental Figure S1.

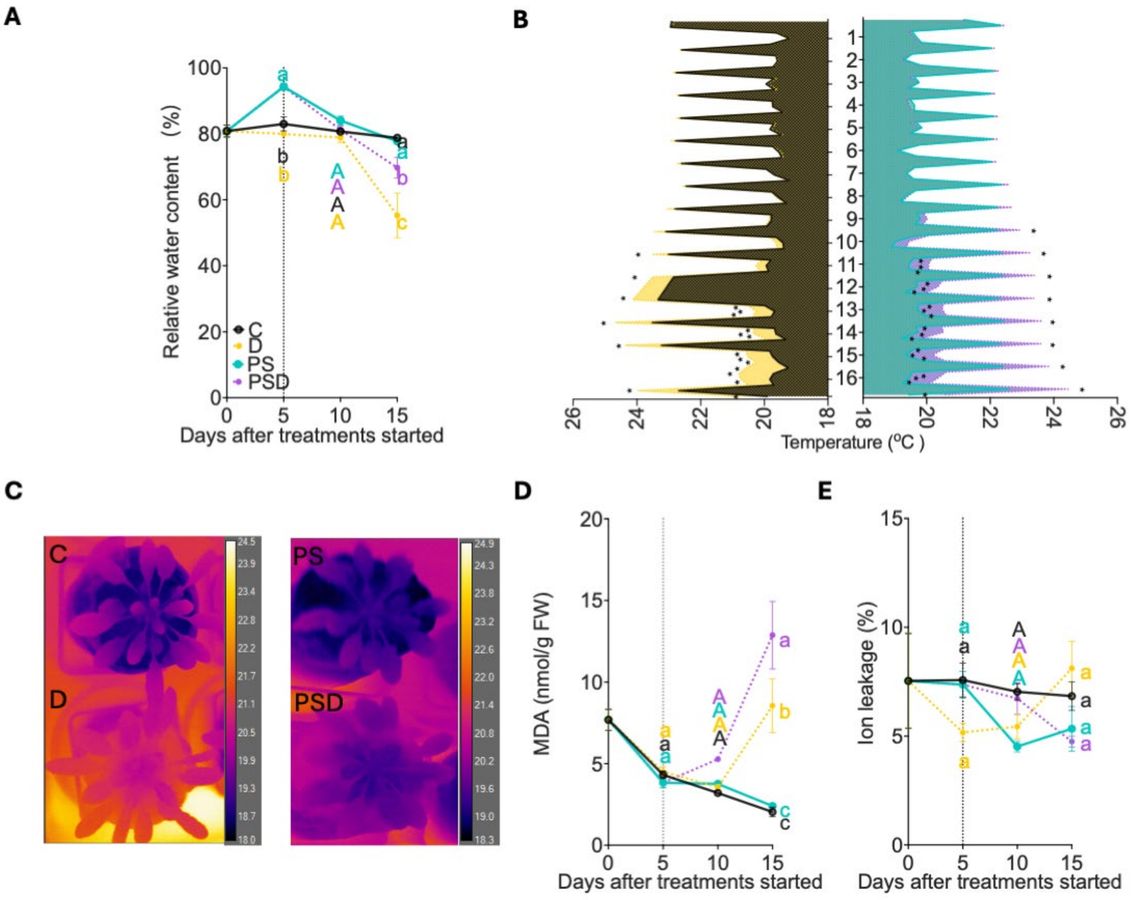


**Supplemental Figure S11.** Effects of submergence followed by recovery under drought or well-watered conditions and corresponding single stresses on physiological traits. **(A)** Rosette water content relative to the maximal water content that the leaves can hold (100%, turgid water - dry weight). n = 15-16. **(B)** Dynamic changes in leaf surface temperature of plants with (PS & PSD, right) and without (C & D, left) pre-submergence treatment. Lines represent the average leaf temperature measured every 6 hours (ZT = 0, 6, 12, and 18 h). n = 3. Asterisks represent significant differences between measured leaf temperature within the same timepoints ($p < 0.05$, One-way ANOVA with Tukey's Post-hoc test). Note that temperature fluctuates between the photoperiod (peak) and dark period. Data from three time points (ZT = 18 h on day 11 and ZT = 0 h and 6 on day 12 in 21 °C) were not recorded due to the camera failure. **(C)** Representative thermal images of plants in absence of prior submergence subjected to control and drought treatments for 15 days (C and D, left), or subjection to submergence followed by either well-watered condition or drought for 10 days (PS and PSD, right). **(D)** Rosette malondialdehyde (MDA) content. n = 5-10. **(E)** Rosette ion leakage relative to the maximal electrolyte conductivity (100%). n = 6. (A, D, E) Error bars indicate means ± SEM. Letters denote significant differences between different treatments within the same time points ($p < 0.05$, 2-way ANOVA with Tukey's Post-hoc test). The dashed vertical lines indicate the moment plants were de-submerged. For treatment abbreviations and used colors, see legend of Supplemental Figure S1.

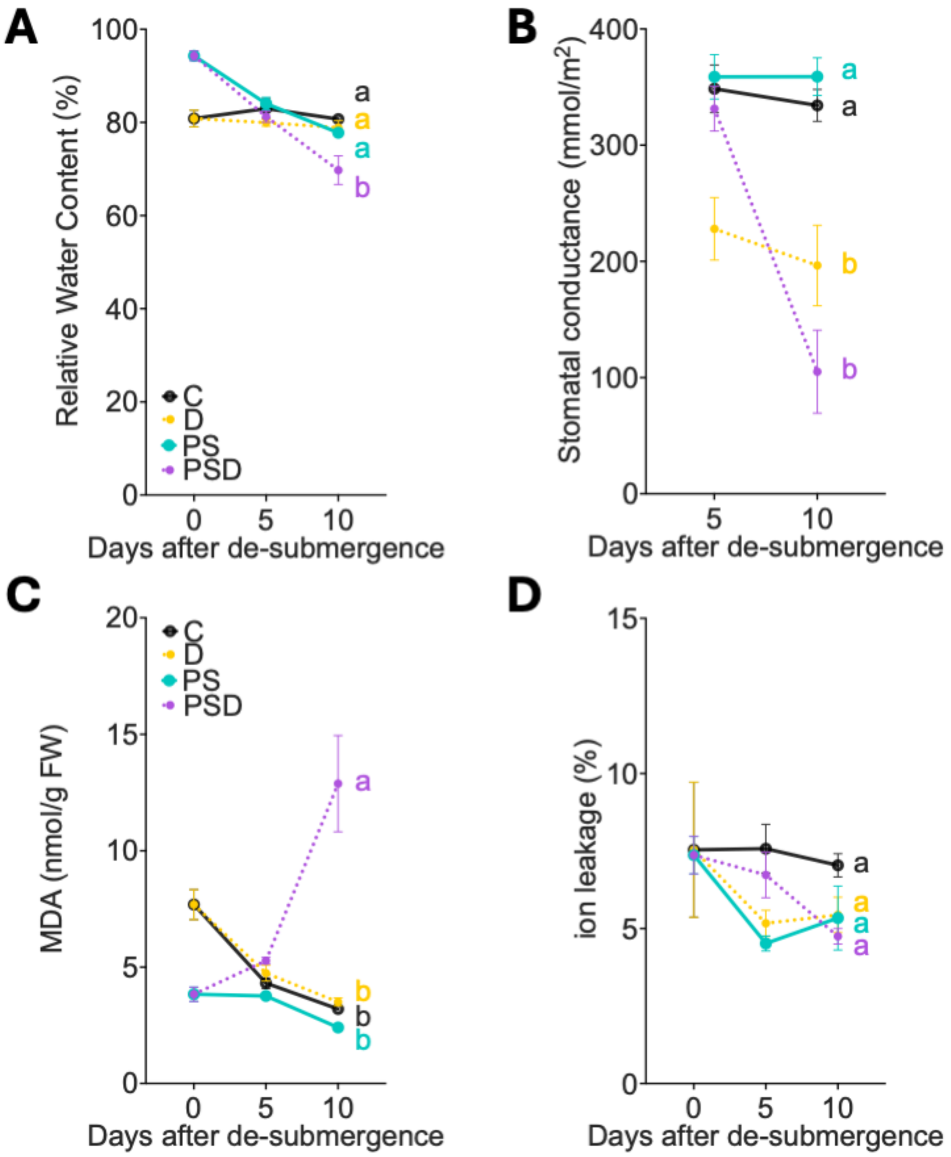


**Supplemental Figure S12.** Physiological traits as effect of drought in presence or absent of prior submergence. **(A)** Rosette water content relative to the maximal water content that the leaves can hold (100%, turgid water - dry weight). n = 15-16. **(B)** Stomatal conductance of young leaves. n = 5-6. **(C)** Rosette Malondialdehyde (MDA) content. n = 5-10. **(D)** Rosette ion leakage relative to the maximal electrolyte conductivity (100%). n = 6. Error bars indicate means ± SEM. Letters denote significant differences between different treatments at the last timepoint of the stress treatments ($p < 0.05$, 2-way ANOVA with Tukey's Post-hoc test). Data presented here is derived from Figure 1 and Supplemental Figure S11. For treatment abbreviations and used colors, see legend of Supplemental Figure S1.

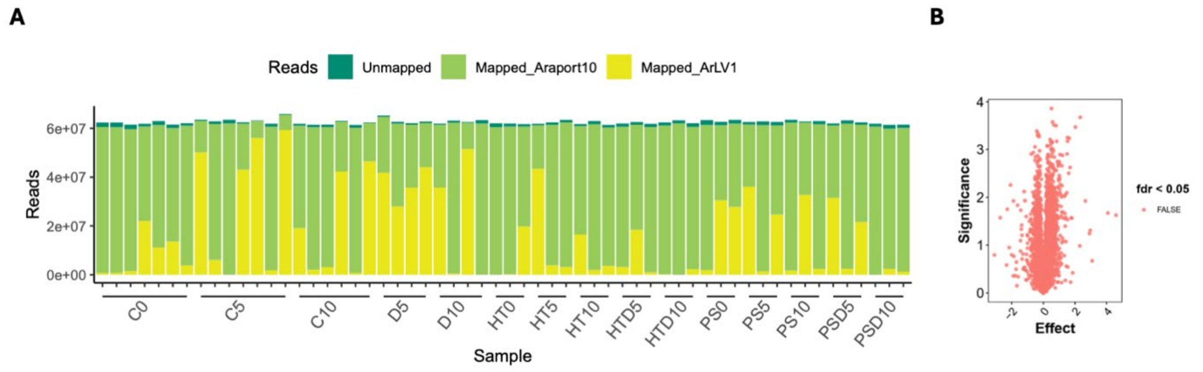


**Supplemental Figure S13.** Sequence coverage of RNA-seq dataset and transcriptomic effects of *ArLV1*. **(A)** Number of sequenced reads mapping to known Arabidopsis genes part of the transcriptome (Mapped_Araport10, green), RNA1 and RNA2 of the *ArLV1* virus (Mapped_ArLV1, yellow) or neither of the two (Unmapped, dark green). **(B)** Volcano plot indicating transcriptomic differences between samples mapping above and below 50% to the *A. thaliana* transcriptome in Control (C) plants at t = 5 days. Genes with FDR (false detection rate) < 0.05 are considered significant.

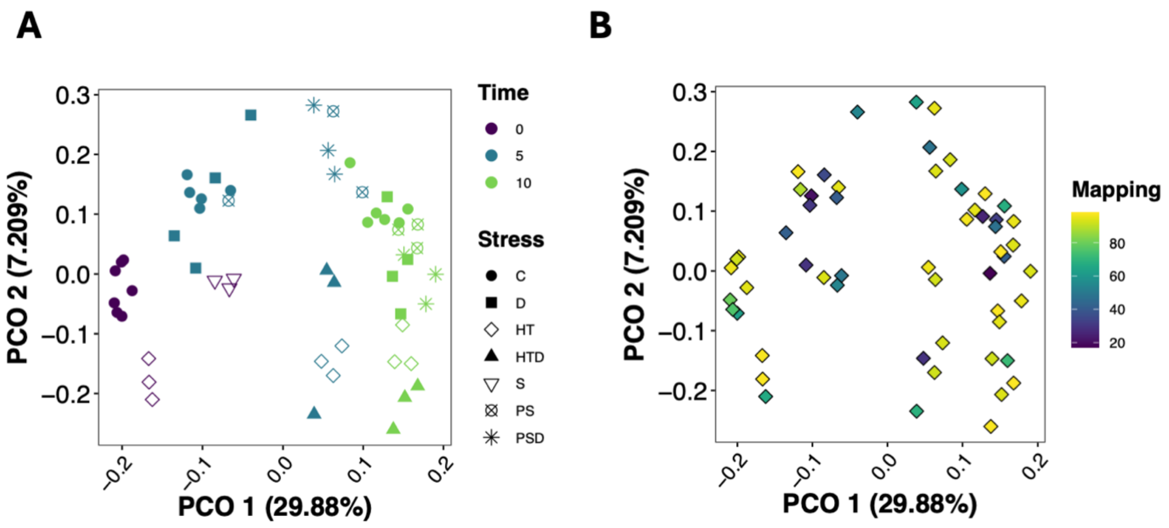


**Supplemental Figure S14. PCA of transcriptomes of young leaves exposed to different stress conditions. (A, B)** PCA analysis visualizing distributions of samples categorized by time (0 = purple, 5 days = blue and 10 days = green) and the different stresses, indicated by symbols (A) and the fraction of reads per sample mapping to the Arabidopsis transcriptome (B). Fraction of reads mapping to the Arabidopsis reference transcriptome is presented by a color scale. For treatment abbreviations see legend of Supplemental Figure S1.

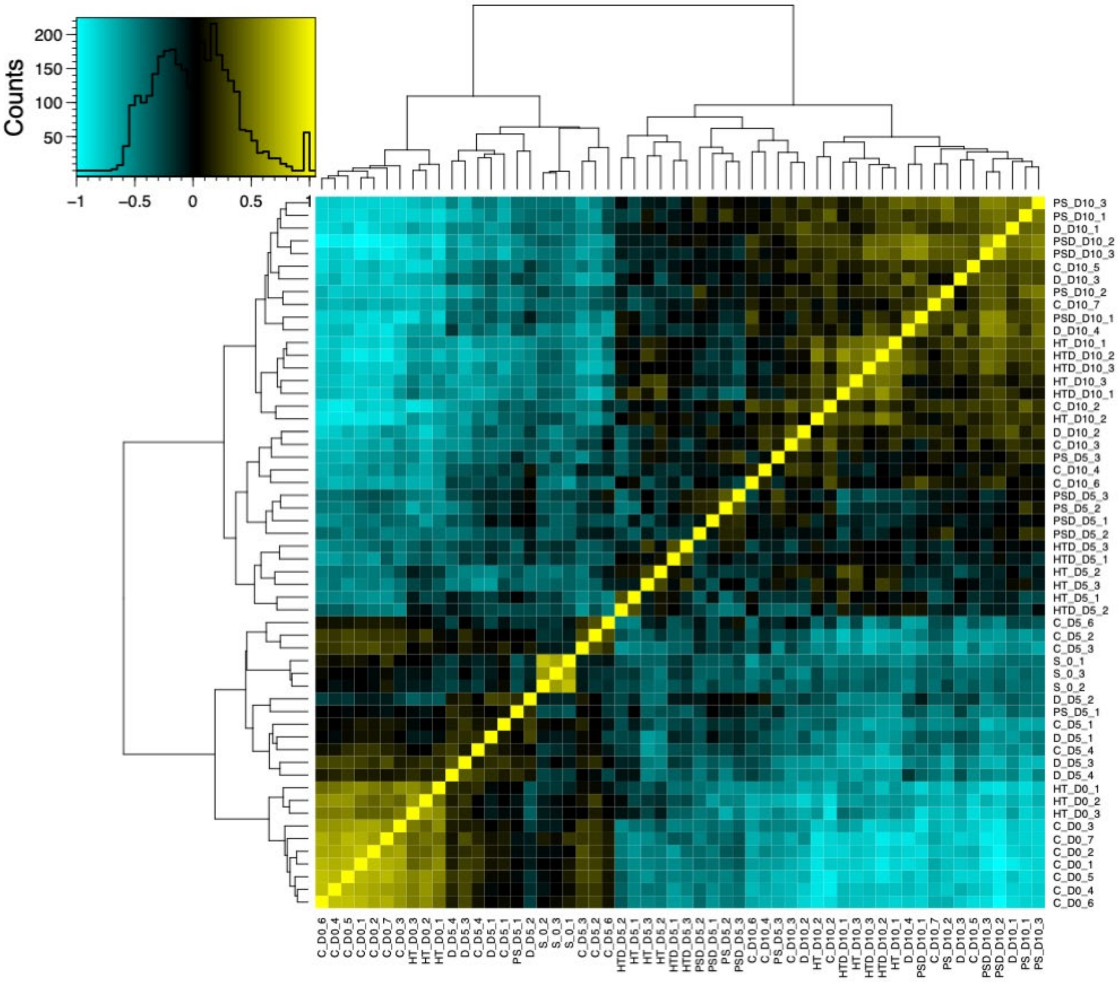


**Supplemental Figure S15.** Correlation matrix of samples in the RNA-seq dataset visualizing the distribution of transcripts of all 56 samples that passed the quality control. The color scale indicates the strength of the correlation (negative: blue, positive: yellow) and the distribution of the matrix values. The relatedness of individual samples are indicated by the hierarchal clustering trees. Samples are named as; treatment abbreviation_timepoint_biological replicate number. For treatment abbreviations see legend of Supplemental Figure S1.

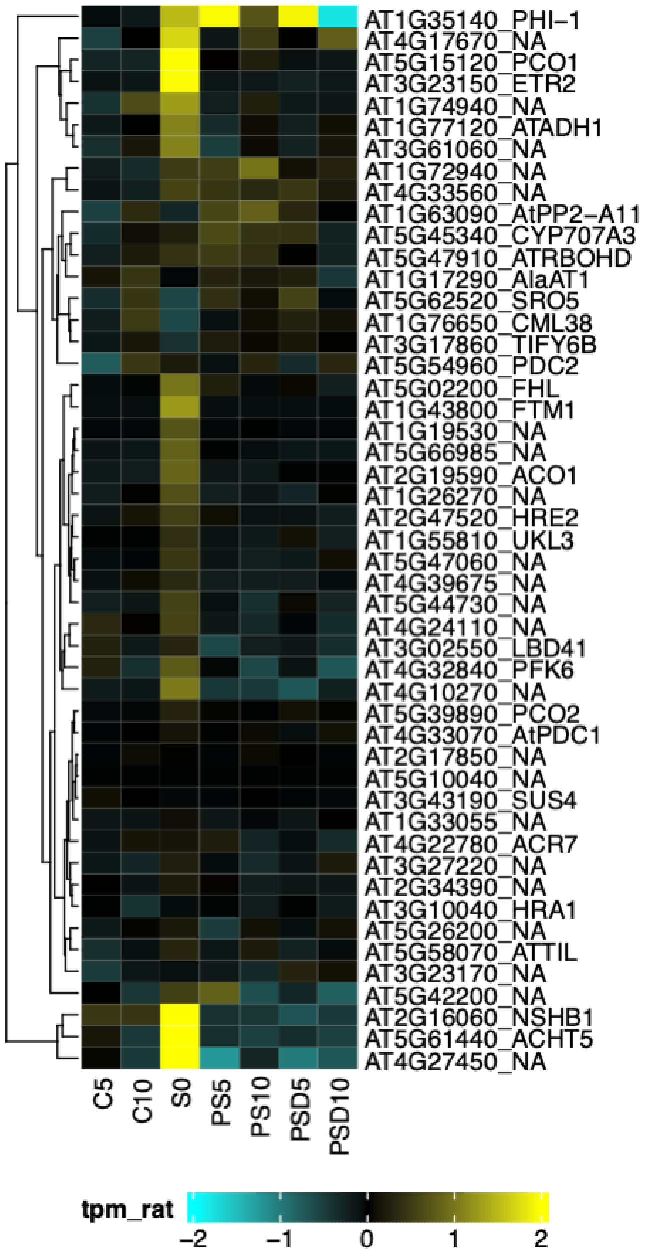


**Supplemental Figure S16.** Relative expression values of 49 known core-hypoxia responsive genes (Mustroph et al., 2009). Heatmap represents the expression ratio in transcripts per million (TPM) of hypoxia responsive genes in response to submergence (S) followed by drought (PSD) and the relevant individual (D and PS) stresses and control (C). Indicated are the AGI gene locus ID and the commonly used abbreviation if available (otherwise indicates as NA). The color scale indicates the expression levels; yellow represents up- and blue represents down-regulation.

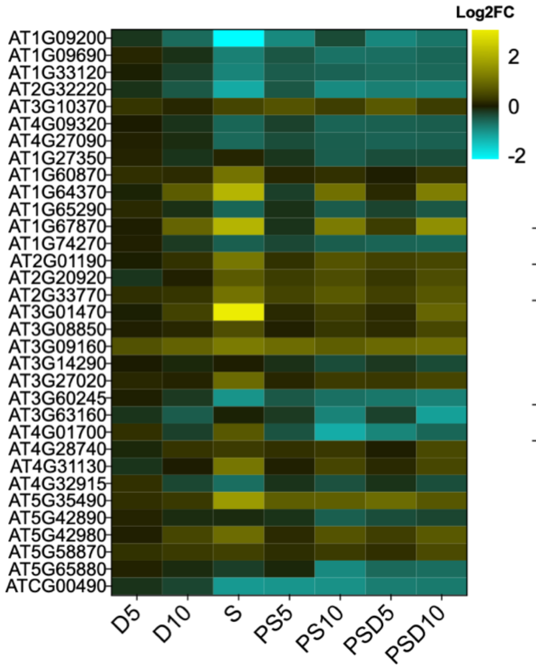


| Category | KEGG |
|---|---|
| Term | ath03010 |
| Description | Ribosome - Arabidopsis thaliana (thale cress) |
| -Log$_{10}$P | 5.13612 |

**Supplemental Figure S17.** Relative expression values of 33 DEGs regulated by both PS and PSD. For each DEG, the AGI gene locus ID is indicated. Color scales indicate Log$_2$FC value (relative to control (C) conditions); yellow and blue indicate up- and down- regulations, respectively. The GO and KEGG enrichment analysis of the 33 DEGs are indicated in the table.

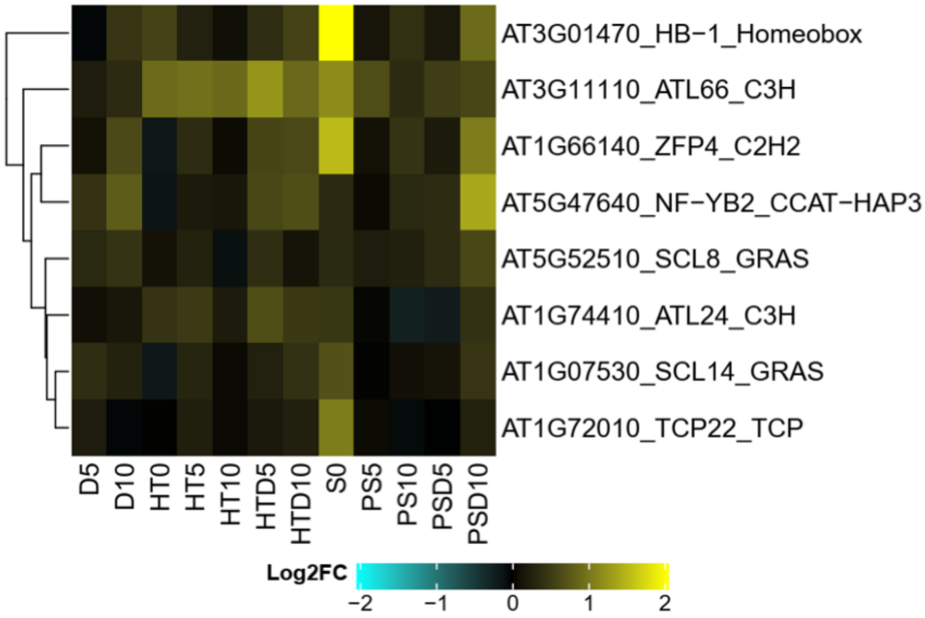


**Supplemental Figure S18.** Relative expression values of 8 TFs that were upregulated in both combined (HTD) and sequential (PSD) stresses, compared to controls (C). Indicated are the AGI gene locus ID and the commonly used abbreviation. Color scales indicate $Log_2FC$ value (relative to control (C) conditions); yellow and blue indicate up- and down- regulations, respectively.

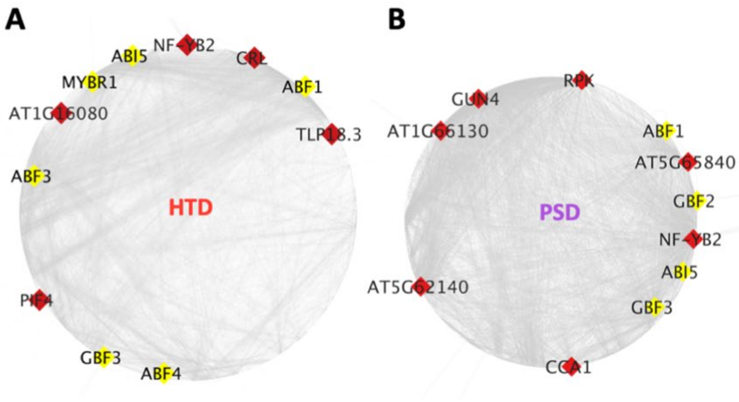


| Gene Name | Locus ID |
|---|---|
| GBF3 | AT2G46270 |
| ABF1 | AT1G49720 |
| ABI5 | AT2G36270 |
| NF-YB2 | AT5G47640 |

**Supplemental Figure S19.(A, B)** Gene regulatory network (GRNs) for DEGs from the upregulated gene clusters in combined high temperature and drought (HTD) (A) and submergence followed by drought (PSD) (B). For each GRN, the putative upstream regulators (11) with the greatest number of connections with the others in the same network are indicated. Genes associated with ABA responses are highlighted with yellow nodes and the common highlighted regulators shared by the two GRNs are indicated in the table on the right (Gene name and AGI locus ID).

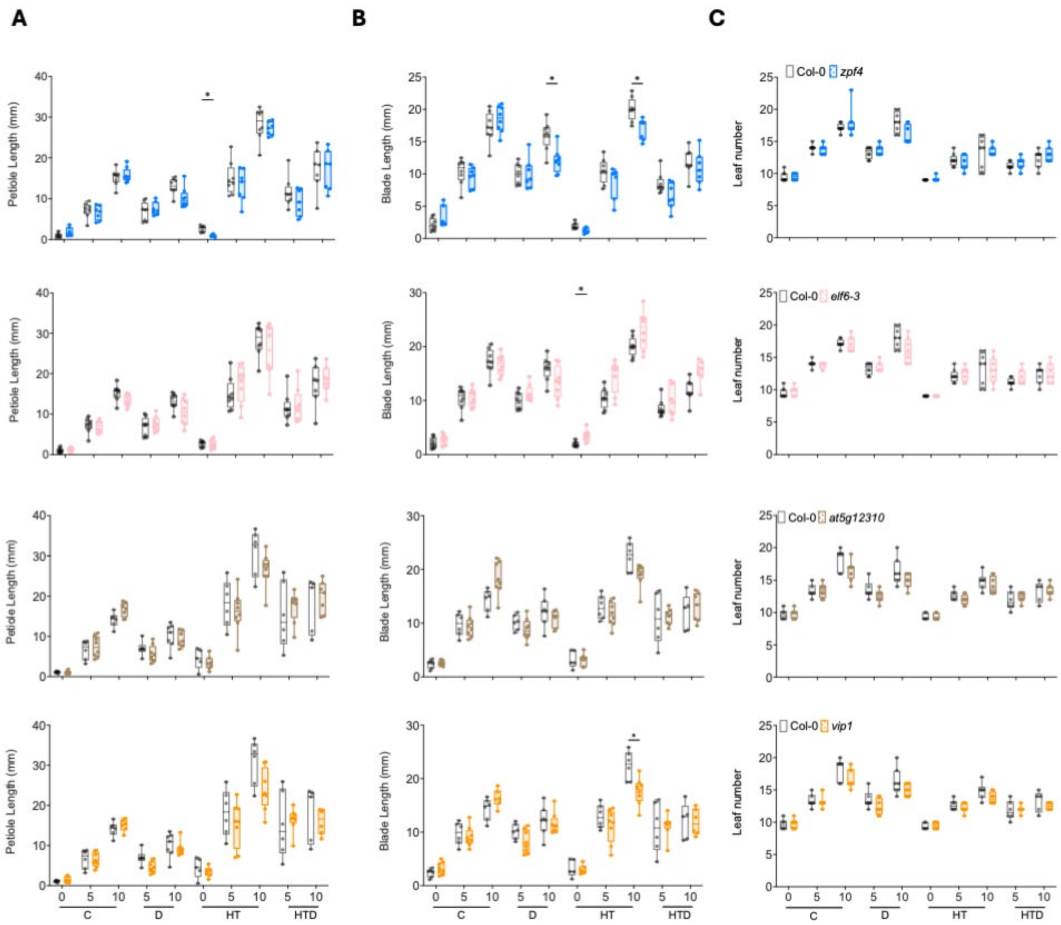


**Supplemental Figure S20.** Effect of combined and individual stresses on leaf development traits of selected mutants. **(A, B, C)** Average lengths of petiole (A) and blade (B) of young leaves, and leaf number (C) of Arabidopsis mutants (*zfp4* (blue), *elf6-3* (pink), *at5g12310* (brown), *vip1* (orange) and wild-type plants (Col-0; grey). Box plots show the median and boxes indicate boundaries of the second and third quartiles (Q1 and Q3) of the data distribution. Whiskers indicate Q1 and Q4 values within 1.5 times the interquartile range. Asterisks represent significant differences between the mutant and the corresponding wild-type plants within the same timepoint ($p < 0.05$, multiple unpaired t-test with Holm-Šídák correction). Numbers indicates days after treatment started. For treatment abbreviations see legend of Supplemental Figure S1. n = 5-10.

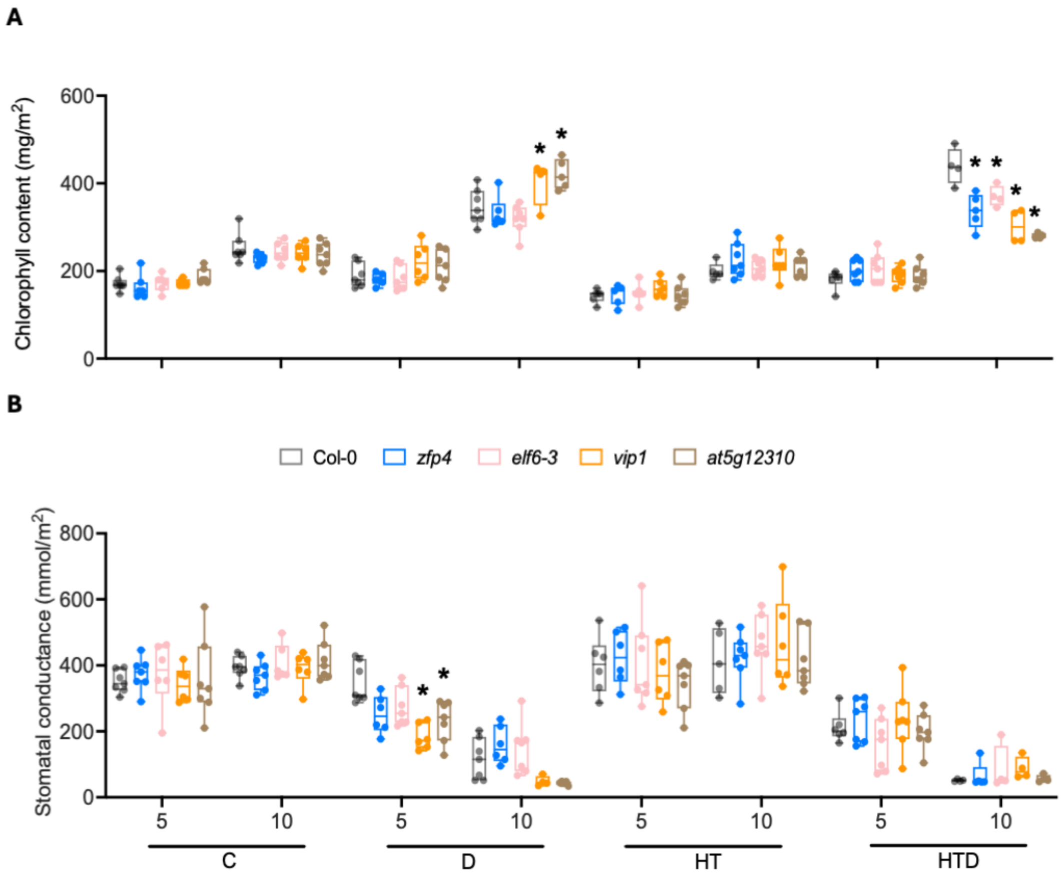


**Supplemental Figure S21.** Effect of combined and related individual stresses on chlorophyll content and stomatal conductance of selected mutants and corresponding wild-type. **(A, B)** chlorophyll content (A) and stomatal conductance (B) of Arabidopsis mutants (*zfp4* (blue), *elf6-3* (pink), *at5g12310* (brown), *vip1* (orange) and *pif4-2* (red)) and the wild-type (Col-0). Boxes indicate boundaries of the second and third quartiles (Q) of the data distribution. Horizontal bars indicate median and whiskers Q1 and Q4 values within 1.5 times the interquartile range. Asterisks represent significant differences between mutant and the corresponding wild-type within the same timepoint ($p < 0.05$, one-way ANOVA with Dunnett test). Numbers indicate days after treatments started. For treatment abbreviations see legend of Supplemental Figure S1. n = 4-7.

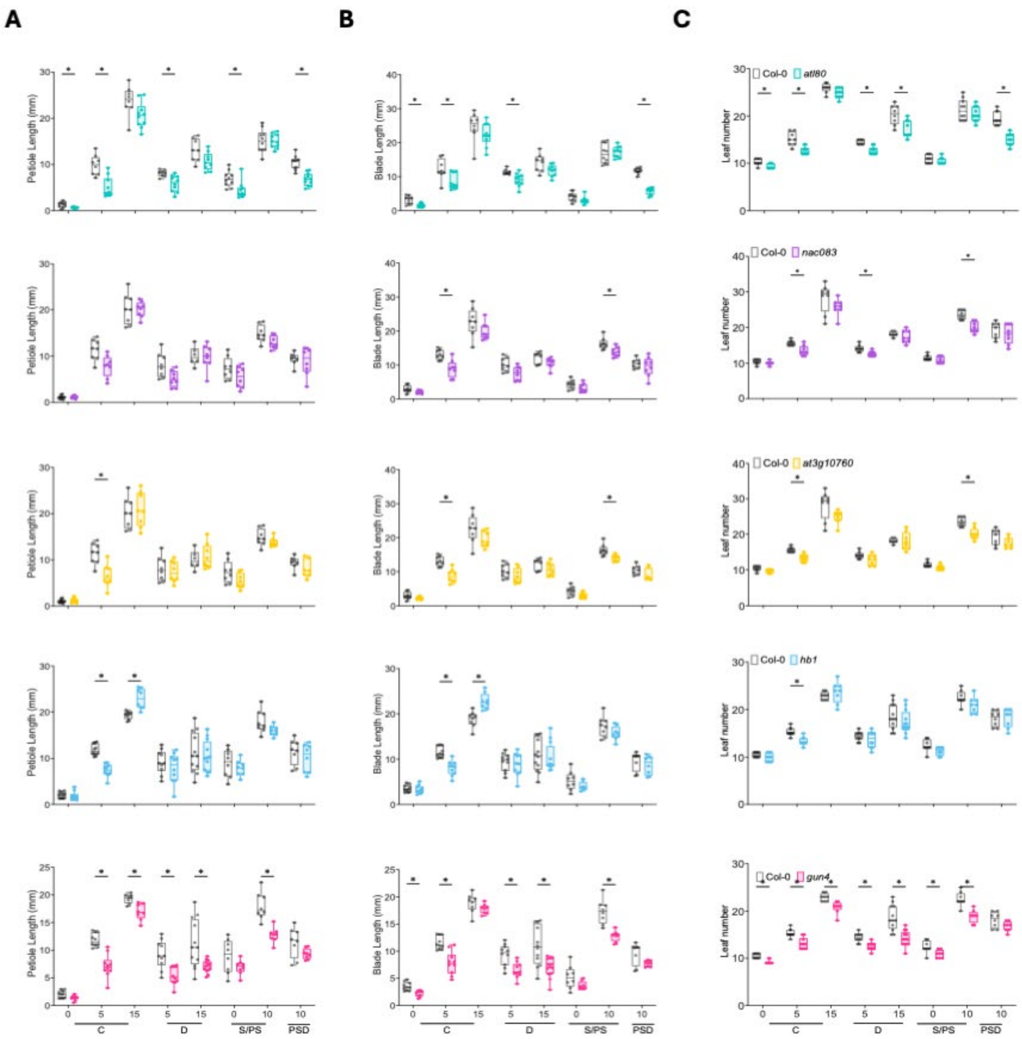


**Supplemental Figure S22.** Effect of sequential and individual stresses on leaf development of selected mutants. **(A, B, C)** Average lengths of petiole (A) and blade (B) of young leaves, and (C) leaf number of Arabidopsis mutants (*atl80* (aqua), *nac083* (purple), *at3g10760* (yellow), *hb1* (azure) and *gun4* (magenta)) and wild-type plants (Col-0; grey). Box plots show the median and boxes indicate boundaries of the second and third quartiles (Q1 and Q3) of the data distribution. Whiskers indicate Q1 and Q4 values within 1.5 times the interquartile range. Asterisks represent significant differences between the mutant and the corresponding wild-type plants within the same timepoint ($p < 0.05$, multiple unpaired t-test with Holm-Šídák correction). Numbers indicates days after treatment started. For treatment abbreviations see legend of Supplemental Figure S1. n = 6-13.

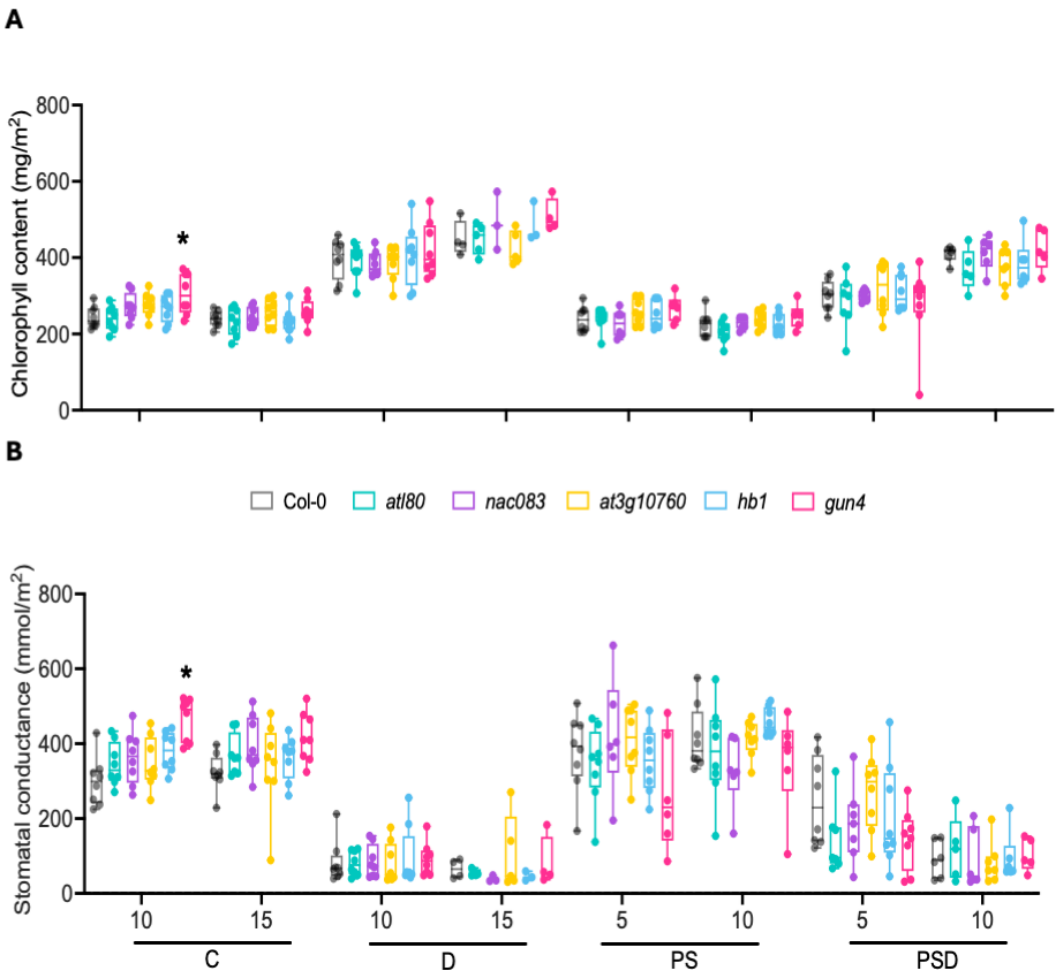


**Supplemental Figure S23.** Effect of sequential and individual stresses on chlorophyll content and stomatal conductance of selected mutants and corresponding wild-type. **(A, B)** chlorophyll content (A) and stomatal conductance (B) of Arabidopsis mutants (*atl80* (aqua), *nac083* (purple), *at3g10760* (yellow), *hb1* (azure) and *gun4* (magenta)) and the wild-type plants (Col-0; gray). Boxes indicate boundaries of the second and third quartiles (Q) of the data distribution. Horizontal bars indicate median and whiskers Q1 and Q4 values within 1.5 times the interquartile range. Asterisks represent significant differences between the mutant and the corresponding wild-type plants within the same time point ($p < 0.05$, one-way ANOVA with Dunnett test). Numbers indicate days after treatment started. For treatment abbreviations see legend of Supplemental Figure S1. n = 3-8.

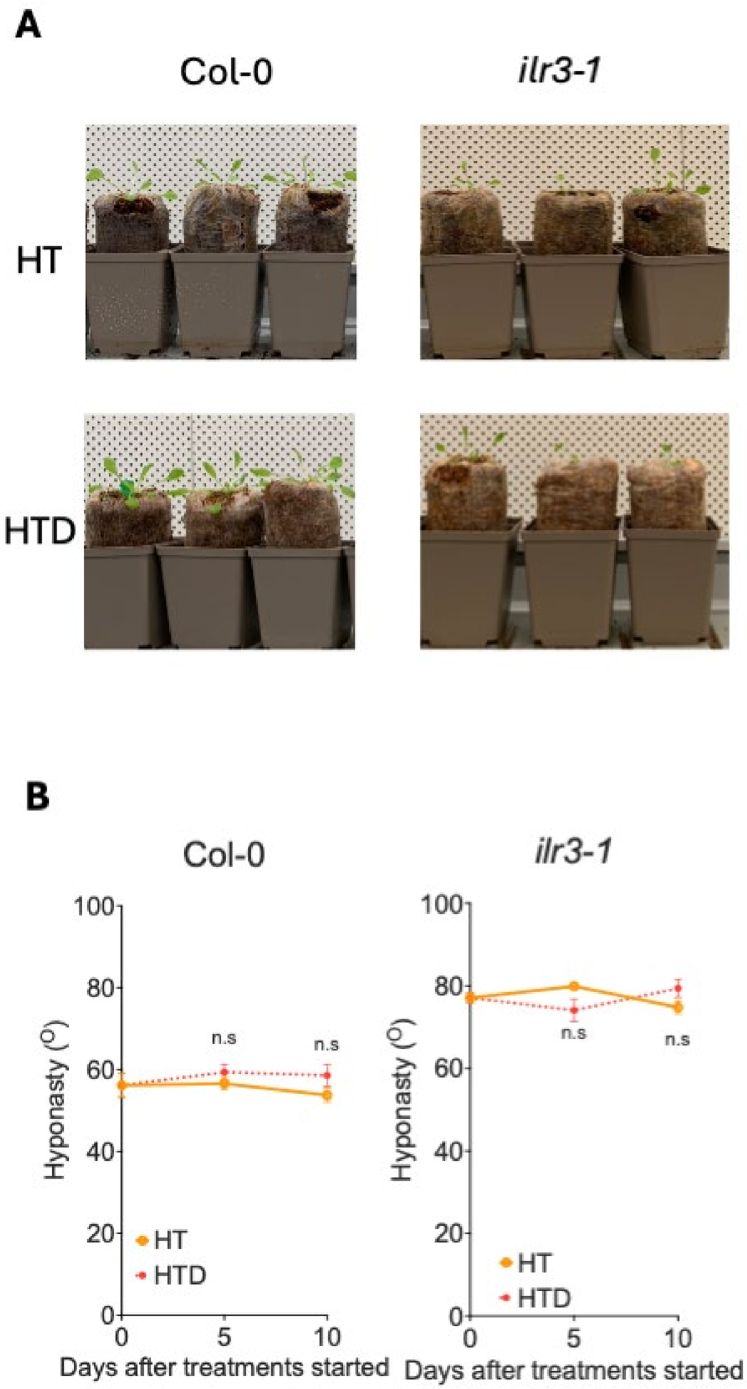


**Supplemental Figure S24.** Typical enhanced leaf hyponasty phenotype of *ilr3* compared to wild-type plants during subjection to high temperature treatments. **(A)** Representative rosette (side) images of Col-0 (left), *ilr3-1* (right) upon 5-day high temperature (HT, upper row) or combined high temperature and drought (HTD, lower row) treatment. **(B)** Average angles of the 2 most hyponastic leaves of individual plants, relative to the horizontal. Red dashed lines represent HTD treatment and solid orange lines indicate HT treatment. The letter 'n.s' represents no significant differences between treatments within the same timepoint ($p>0.05$, unpaired t-test). For treatment abbreviations and used colors, see legend of Supplemental Figure S1. n = 5-10.

**A**

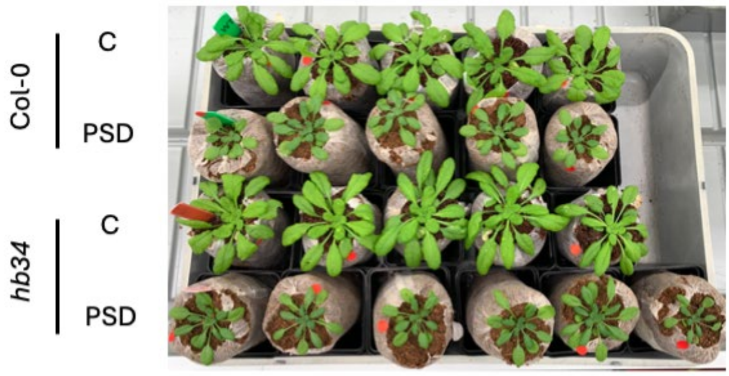


**B**

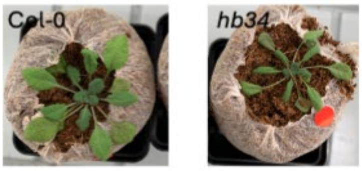


**Supplemental Figure S25.** Representative images of Col-0 (upper two rows) and *hb34* mutant (lower two rows) plants on Jiffy 7c coconut pellet growth substrate, subjected to control (C; 1st and 3rd row) and submergence followed by drought (PSD; 2nd and 4th row) at **(A)** 15 days after treatments started and **(B)** the day of wilting.

### Supplemental Tables

**Supplemental Table S1.** Significantly enriched GO terms ($p < 0.01$) of biological processes in each of the *k-means* clusters under combined high temperature and drought stress (HTD). Indicated are the GO category (GO accession), term description, p-value of GO term enrichment and the *k-means* clusters in which the term was enriched.

| GO accession | Term description | p-value | Cluster |
|---|---|---|---|
| GO:0042254 | ribosome biogenesis | 6.03979E-40 | 1 |
| GO:0042273 | ribosomal large subunit biogenesis | 1.31912E-28 | 1 |
| GO:0002181 | cytoplasmic translation | 4.54098E-20 | 1 |
| GO:0042255 | ribosome assembly | 1.23714E-15 | 1 |
| GO:0042274 | ribosomal small subunit biogenesis | 1.76745E-11 | 1 |
| GO:0006119 | oxidative phosphorylation | 4.70525E-07 | 1 |
| GO:0072344 | rescue of stalled ribosome | 2.7165E-05 | 1 |
| GO:0001510 | RNA methylation | 3.61458E-05 | 1 |
| GO:0032544 | plastid translation | 0.000431133 | 1 |
| GO:0034728 | nucleosome organization | 0.000485249 | 1 |
| GO:0000413 | protein peptidyl-prolyl isomerization | 0.000912284 | 1 |
| GO:0016074 | sno(s)RNA metabolic process | 0.001450317 | 1 |
| GO:0097549 | chromatin organization involved in negative regulation of transcription | 0.001494455 | 1 |
| GO:0043038 | amino acid activation | 0.001783933 | 1 |
| GO:0006091 | generation of precursor metabolites and energy | 0.00011082 | 2 |
| GO:0016192 | vesicle-mediated transport | 0.00157761 | 2 |
| GO:0007030 | Golgi organization | 0.00319522 | 2 |

| | | | |
|---|---|---|---|
| GO:0006403 | RNA localization | 1.20322E-07 | 3 |
| GO:0009657 | plastid organization | 1.30079E-06 | 3 |
| GO:0000375 | RNA splicing, via transesterification reactions | 1.36781E-05 | 3 |
| GO:0030433 | ubiquitin-dependent ERAD pathway | 2.27199E-05 | 3 |
| GO:0034660 | ncRNA metabolic process | 2.84923E-05 | 3 |
| GO:0000373 | Group II intron splicing | 3.26535E-05 | 3 |
| GO:1903311 | regulation of mRNA metabolic process | 0.000139591 | 3 |
| GO:0000959 | mitochondrial RNA metabolic process | 0.000185024 | 3 |
| GO:0006457 | protein folding | 0.000276219 | 3 |
| GO:0009790 | embryo development | 0.000381017 | 3 |
| GO:0006281 | DNA repair | 0.002059724 | 3 |
| GO:0048574 | long-day photoperiodism, flowering | 0.003376124 | 3 |
| GO:0140053 | mitochondrial gene expression | 0.004569125 | 3 |
| GO:0065002 | intracellular protein transmembrane transport | 0.006145542 | 3 |
| GO:0022613 | ribonucleoprotein complex biogenesis | 0.006222077 | 3 |
| GO:0006897 | endocytosis | 0.006450208 | 3 |
| GO:0090305 | nucleic acid phosphodiester bond hydrolysis | 0.007427573 | 3 |
| GO:0042026 | protein refolding | 0.008236804 | 3 |
| GO:0009408 | response to heat | 1.56187E-05 | 4 |
| GO:0016560 | protein import into peroxisome matrix, docking | 3.73094E-05 | 4 |
| GO:0006979 | response to oxidative stress | 0.000104547 | 4 |
| GO:1901607 | alpha-amino acid biosynthetic process | 0.000192445 | 4 |
| GO:0048878 | chemical homeostasis | 0.000337441 | 4 |

| | | | |
|---|---|---|---|
| GO:0009644 | response to high light intensity | 0.000443147 | 4 |
| GO:0006790 | sulfur compound metabolic process | 0.000716216 | 4 |
| GO:0009611 | response to wounding | 0.000954123 | 4 |
| GO:0120253 | hydrocarbon catabolic process | 0.001017665 | 4 |
| GO:0016485 | protein processing | 0.00127862 | 4 |
| GO:0010150 | leaf senescence | 0.001773684 | 4 |
| GO:0046471 | phosphatidylglycerol metabolic process | 0.001807701 | 4 |
| GO:0019684 | photosynthesis, light reaction | 0.00183533 | 4 |
| GO:0042594 | response to starvation | 0.001857114 | 4 |
| GO:0009620 | response to fungus | 0.001947046 | 4 |
| GO:0009743 | response to carbohydrate | 0.002241505 | 4 |
| GO:0006914 | autophagy | 0.003415799 | 4 |
| GO:0006790 | sulfur compound metabolic process | 0.000716216 | 4 |

**Supplemental Table S2.** Significantly enriched GO terms ($p < 0.01$) of biological processes in each of the *k-means* clusters under submergence followed by drought treatment. Indicated are the GO category (accession), term description, p-value of GO term enrichment and the *k-means* clusters in which the term was enriched.

| GO accession | Term description | p-value | Cluster |
|---|---|---|---|
| GO:0034660 | ncRNA metabolic process | 2.91039E-05 | 1 |
| GO:0042793 | plastid transcription | 6.05524E-05 | 1 |
| GO:0000373 | Group II intron splicing | 0.001097608 | 1 |
| GO:0000959 | mitochondrial RNA metabolic process | 0.00110131 | 1 |
| GO:0006281 | DNA repair | 0.003283367 | 1 |
| GO:0065002 | intracellular protein transmembrane transport | 0.003663384 | 1 |
| GO:0006334 | nucleosome assembly | 0.004910088 | 1 |
| GO:0072527 | pyrimidine-containing compound metabolic process | 0.008403109 | 1 |
| GO:0042254 | ribosome biogenesis | 2.10002E-32 | 2 |
| GO:0002181 | cytoplasmic translation | 1.61112E-18 | 2 |
| GO:0042255 | ribosome assembly | 1.16976E-12 | 2 |
| GO:0015995 | chlorophyll biosynthetic process | 4.91082E-09 | 2 |
| GO:0042274 | ribosomal small subunit biogenesis | 5.72558E-08 | 2 |
| GO:0006417 | regulation of translation | 9.99943E-05 | 2 |
| GO:0032544 | plastid translation | 0.00039216 | 2 |
| GO:0009657 | plastid organization | 0.000621101 | 2 |
| GO:0045036 | protein targeting to chloroplast | 0.00114847 | 2 |
| GO:1901259 | chloroplast rRNA processing | 0.003910888 | 2 |
| GO:0015979 | photosynthesis | 1.6229E-06 | 3 |

| | | | |
|---|---|---|---|
| GO:0010258 | NADH dehydrogenase complex (plastoquinone) assembly | 3.7316E-05 | 3 |
| GO:0009110 | vitamin biosynthetic process | 0.00042279 | 3 |
| GO:0009642 | response to light intensity | 0.00202643 | 3 |
| GO:0006778 | porphyrin-containing compound metabolic process | 0.0020781 | 3 |
| GO:0030001 | metal ion transport | 0.00403665 | 3 |
| GO:0070417 | cellular response to cold | 0.00462344 | 3 |
| GO:0016051 | carbohydrate biosynthetic process | 0.00488961 | 3 |
| GO:0001101 | response to acid chemical | 0.00749204 | 3 |
| GO:0051604 | protein maturation | 0.0084925 | 3 |
| GO:0046148 | pigment biosynthetic process | 0.00149185 | 4 |
| GO:0009639 | response to red or far-red light | 0.00677587 | 4 |

**Supplemental Table S3.** Relative expression values of upregulated TFs identified in combined high temperature and drought. Indicated are the the commonly used abbreviation (Gene name), AGI gene locus ID, $Log_2FC$ effect on transcription and p values of the effect per treatment and timepoint.

| | | D5 | | D10 | | HT0 | | HT5 | | HT10 | | HTD5 | | HTD10 | |
|---|---|---|---|---|---|---|---|---|---|---|---|---|---|---|---|
| **Gene Name** | **Locus ID** | **$Log_2FC$** | **p Value** | **$Log_2FC$** | **p Value** | **$Log_2FC$** | **p Value** | **$Log_2FC$** | **p Value** | **$Log_2FC$** | **p Value** | **$Log_2FC$** | **p Value** | **$Log_2FC$** | **p Value** |
| *HSFB2B* | AT4G11660 | 0.425 | 0.210 | 0.243 | 0.782 | 0.220 | 0.353 | 0.358 | 0.540 | 0.675 | 0.260 | 0.977 | 0.037 | 1.600 | 0.015 |
| *HSF4* | AT4G36990 | -0.320 | 0.754 | 0.377 | 0.629 | 0.430 | 0.231 | 1.103 | 0.311 | 0.365 | 0.541 | 0.797 | 0.407 | 1.363 | 0.024 |
| *WHY2* | AT1G71260 | 0.184 | 0.649 | 0.275 | 0.523 | 0.409 | 0.066 | 0.432 | 0.091 | 0.840 | 0.053 | 0.363 | 0.138 | 1.048 | 0.010 |
| *VIP1* | AT1G43700 | 0.264 | 0.303 | 0.175 | 0.480 | 0.113 | 0.732 | 0.845 | 0.045 | 0.571 | 0.039 | 0.712 | 0.033 | 0.924 | 0.002 |
| *ATL66* | AT3G11110 | 0.206 | 0.577 | 0.343 | 0.493 | 0.905 | 0.099 | 0.947 | 0.085 | 0.891 | 0.061 | 1.233 | 0.049 | 0.882 | 0.008 |
| *DEAR3* | AT2G23340 | 0.377 | 0.245 | 0.463 | 0.527 | 0.562 | 0.322 | 1.855 | 0.042 | 0.236 | 0.729 | 1.250 | 0.016 | 0.826 | 0.108 |
| *NF-YB2* | AT5G47640 | 0.420 | 0.150 | 0.779 | 0.481 | -0.124 | 0.809 | 0.184 | 0.571 | 0.158 | 0.808 | 0.601 | 0.031 | 0.665 | 0.258 |
| *ZFP4* | AT1G66140 | 0.104 | 0.779 | 0.623 | 0.392 | -0.156 | 0.683 | 0.355 | 0.497 | 0.059 | 0.826 | 0.587 | 0.095 | 0.617 | 0.019 |
| *TLP7* | AT1G53320 | 0.229 | 0.396 | 0.019 | 0.940 | 0.275 | 0.318 | 0.764 | 0.094 | 0.143 | 0.344 | 0.473 | 0.135 | 0.612 | 0.041 |
| *BZIP28* | AT3G10800 | 0.068 | 0.802 | 0.169 | 0.661 | 0.073 | 0.818 | 0.418 | 0.215 | 0.259 | 0.252 | 0.426 | 0.143 | 0.602 | 0.012 |
| *ERF3* | AT1G50640 | 0.270 | 0.369 | 0.338 | 0.507 | 0.544 | 0.122 | 1.074 | 0.029 | 0.312 | 0.302 | 0.780 | 0.060 | 0.592 | 0.056 |
| *BZIP17* | AT2G40950 | 0.175 | 0.533 | 0.168 | 0.456 | -0.138 | 0.453 | 0.289 | 0.304 | 0.166 | 0.306 | 0.112 | 0.684 | 0.570 | 0.004 |
| *HB1* | AT3G01470 | -0.034 | 0.872 | 0.437 | 0.444 | 0.567 | 0.270 | 0.254 | 0.286 | 0.092 | 0.522 | 0.375 | 0.045 | 0.561 | 0.063 |
| *ATL24* | AT1G74410 | 0.096 | 0.644 | 0.155 | 0.621 | 0.427 | 0.090 | 0.491 | 0.215 | 0.196 | 0.281 | 0.660 | 0.068 | 0.483 | 0.032 |
| *ELF6* | AT5G04240 | 0.355 | 0.111 | 0.226 | 0.462 | 0.143 | 0.584 | 0.334 | 0.235 | 0.152 | 0.373 | 0.323 | 0.131 | 0.444 | 0.018 |
| *SE* | AT2G27100 | 0.260 | 0.076 | 0.243 | 0.310 | -0.201 | 0.554 | 0.001 | 0.995 | 0.240 | 0.079 | 0.127 | 0.147 | 0.412 | 0.024 |

| | | | | | | | | | | | | | | | |
|---|---|---|---|---|---|---|---|---|---|---|---|---|---|---|---|
| *SCL14* | AT1G07530 | 0.375 | 0.173 | 0.261 | 0.452 | -0.160 | 0.505 | 0.292 | 0.408 | 0.048 | 0.752 | 0.248 | 0.361 | 0.407 | 0.008 |
| *CDC5* | AT1G09770 | 0.131 | 0.228 | 0.144 | 0.479 | -0.165 | 0.482 | 0.100 | 0.301 | 0.054 | 0.712 | -0.065 | 0.528 | 0.406 | 0.013 |
| *PCFS4* | AT4G04885 | 0.212 | 0.376 | 0.139 | 0.515 | -0.003 | 0.995 | 0.313 | 0.158 | 0.118 | 0.598 | 0.190 | 0.356 | 0.394 | 0.017 |
| *MYBR1* | AT5G67300 | -0.142 | 0.805 | 0.406 | 0.755 | 0.239 | 0.826 | 1.630 | 0.030 | 0.418 | 0.700 | 0.267 | 0.741 | 0.392 | 0.662 |
| *ARID5* | AT3G43240 | -0.153 | 0.522 | 0.110 | 0.532 | -0.110 | 0.721 | 0.056 | 0.867 | 0.095 | 0.531 | -0.212 | 0.418 | 0.390 | 0.023 |
| *AT1G01930* | AT1G01930 | 0.532 | 0.144 | 0.087 | 0.668 | 0.143 | 0.382 | 0.499 | 0.147 | 0.283 | 0.093 | 0.547 | 0.087 | 0.380 | 0.018 |
| *TCP23* | AT1G35560 | 0.211 | 0.364 | 0.225 | 0.661 | 0.343 | 0.093 | 0.654 | 0.082 | 0.304 | 0.270 | 0.847 | 0.018 | 0.380 | 0.098 |
| *TCP9* | AT2G45680 | 0.037 | 0.944 | 0.328 | 0.702 | -0.316 | 0.409 | 0.783 | 0.025 | 0.856 | 0.164 | -0.078 | 0.766 | 0.330 | 0.530 |
| *AT5G12310* | AT5G12310 | 0.379 | 0.237 | 0.251 | 0.481 | 0.302 | 0.252 | 0.275 | 0.435 | 0.257 | 0.294 | 0.474 | 0.135 | 0.313 | 0.040 |
| *ILR3* | AT5G54680 | -0.108 | 0.472 | -0.035 | 0.786 | 0.107 | 0.576 | 0.322 | 0.082 | 0.157 | 0.195 | 0.271 | 0.084 | 0.265 | 0.028 |
| *ATL13* | AT3G60080 | 0.389 | 0.088 | 0.243 | 0.524 | 0.256 | 0.144 | 0.204 | 0.167 | 0.228 | 0.351 | 0.414 | 0.032 | 0.261 | 0.216 |
| *BEH4* | AT1G78700 | 0.223 | 0.105 | 0.051 | 0.850 | 0.005 | 0.994 | 0.267 | 0.233 | 0.043 | 0.857 | 0.440 | 0.022 | 0.243 | 0.106 |
| *TCP22* | AT1G72010 | 0.209 | 0.382 | -0.031 | 0.900 | 0.011 | 0.931 | 0.229 | 0.611 | 0.051 | 0.769 | 0.168 | 0.498 | 0.235 | 0.028 |
| *SCL1* | AT1G21450 | 0.321 | 0.105 | 0.031 | 0.903 | 0.320 | 0.198 | 0.419 | 0.074 | 0.082 | 0.351 | 0.299 | 0.026 | 0.226 | 0.029 |
| *AT1G80400* | AT1G80400 | 0.218 | 0.237 | 0.113 | 0.507 | 0.233 | 0.219 | 0.227 | 0.181 | -0.009 | 0.948 | 0.418 | 0.042 | 0.215 | 0.033 |
| *HB2* | AT4G16780 | 0.447 | 0.215 | 0.265 | 0.574 | 0.761 | 0.022 | 0.947 | 0.038 | -0.010 | 0.992 | 0.599 | 0.139 | 0.137 | 0.698 |
| *SCL8* | AT5G52510 | 0.319 | 0.153 | 0.422 | 0.361 | 0.113 | 0.458 | 0.264 | 0.286 | -0.093 | 0.764 | 0.379 | 0.041 | 0.130 | 0.508 |
| *BEL1* | AT5G41410 | 0.460 | 0.190 | 0.273 | 0.479 | 0.490 | 0.128 | 0.112 | 0.792 | -0.223 | 0.470 | 0.645 | 0.028 | 0.051 | 0.846 |
| *MYB34* | AT5G60890 | 0.414 | 0.391 | 0.243 | 0.782 | 0.901 | 0.188 | 1.368 | 0.031 | -0.109 | 0.905 | 0.024 | 0.945 | -0.214 | 0.691 |

**Supplemental Table S4.** Relative expression values of upregulated TFs identified in submergence followed by drought. Indicated are the the commonly used abbreviation (Gene name), AGI gene locus ID, $Log_2FC$ effect on transcription and p values of the effect per treatment and timepoint.

| | | D5 | | D10 | | S/PS0 | | PS5 | | PS10 | | PSD5 | | PSD10 | |
|---|---|---|---|---|---|---|---|---|---|---|---|---|---|---|---|
| **Gene Name** | **Locus ID** | **Log2FC** | **p Value** | **Log2FC** | **p Value** | **Log2FC** | **p Value** | **Log2FC** | **p Value** | **Log2FC** | **p Value** | **Log2FC** | **p Value** | **Log2FC** | **p Value** |
| *CGA1* | AT4G26150 | 0.470 | 0.192 | 1.254 | 0.404 | 0.478 | 0.126 | 0.907 | 0.144 | 1.997 | 0.060 | 0.444 | 0.185 | 2.239 | 0.028 |
| *NF-YB2* | AT5G47640 | 0.420 | 0.150 | 0.779 | 0.481 | 0.335 | 0.284 | 0.056 | 0.896 | 0.321 | 0.529 | 0.355 | 0.127 | 1.378 | 0.033 |
| *ZFP4* | AT1G66140 | 0.104 | 0.779 | 0.623 | 0.392 | 1.511 | 0.000 | 0.100 | 0.816 | 0.423 | 0.164 | 0.183 | 0.701 | 1.052 | 0.013 |
| *HB1* | AT3G01470 | -0.034 | 0.872 | 0.437 | 0.444 | 3.098 | 0.000 | 0.138 | 0.843 | 0.398 | 0.048 | 0.171 | 0.309 | 0.904 | 0.011 |
| *NAC083* | AT5G13180 | 0.451 | 0.175 | 0.230 | 0.604 | 0.785 | 0.001 | 0.087 | 0.819 | 0.155 | 0.644 | 0.271 | 0.469 | 0.784 | 0.041 |
| *WRKY17* | AT2G24570 | -0.083 | 0.804 | 0.654 | 0.444 | 0.039 | 0.884 | 0.003 | 0.995 | 1.174 | 0.048 | 0.094 | 0.856 | 0.733 | 0.097 |
| *FBH2* | AT4G09180 | 0.375 | 0.076 | 0.507 | 0.443 | 0.090 | 0.597 | 0.504 | 0.116 | 0.341 | 0.145 | 0.145 | 0.528 | 0.656 | 0.012 |
| *ERF34* | AT2G44940 | 0.282 | 0.506 | 0.395 | 0.478 | 0.279 | 0.402 | 0.351 | 0.528 | 0.119 | 0.635 | 0.410 | 0.613 | 0.641 | 0.037 |
| *RGA1* | AT2G01570 | 0.101 | 0.503 | 0.406 | 0.444 | 0.534 | 0.007 | 0.290 | 0.155 | 0.576 | 0.087 | 0.198 | 0.262 | 0.610 | 0.037 |
| *SCL8* | AT5G52510 | 0.319 | 0.153 | 0.422 | 0.361 | 0.331 | 0.027 | 0.215 | 0.247 | 0.242 | 0.344 | 0.358 | 0.111 | 0.605 | 0.035 |
| *ATL66* | AT3G11110 | 0.206 | 0.577 | 0.343 | 0.493 | 1.166 | 0.014 | 0.652 | 0.223 | 0.351 | 0.155 | 0.518 | 0.195 | 0.594 | 0.041 |
| *DOF2* | AT3G21270 | 0.243 | 0.551 | 0.198 | 0.603 | 1.243 | 0.001 | 0.296 | 0.485 | 0.398 | 0.197 | 0.238 | 0.497 | 0.559 | 0.045 |
| *DOF1* | AT1G51700 | 0.369 | 0.129 | 0.484 | 0.584 | 0.295 | 0.272 | 0.551 | 0.334 | 0.592 | 0.327 | 0.537 | 0.045 | 0.520 | 0.223 |
| *SCL14* | AT1G07530 | 0.375 | 0.173 | 0.261 | 0.452 | 0.692 | 0.001 | 0.015 | 0.971 | 0.098 | 0.316 | 0.129 | 0.679 | 0.438 | 0.013 |
| *HB34* | AT3G28920 | 0.268 | 0.228 | 0.074 | 0.832 | 0.001 | 0.993 | 0.160 | 0.611 | 0.383 | 0.087 | 0.305 | 0.255 | 0.430 | 0.039 |
| *AT3G10760* | AT3G10760 | 0.402 | 0.190 | 0.288 | 0.342 | 1.250 | 0.000 | 0.327 | 0.379 | 0.266 | 0.082 | 0.279 | 0.423 | 0.408 | 0.035 |
| *ATL24* | AT1G74410 | 0.096 | 0.644 | 0.155 | 0.621 | 0.460 | 0.016 | 0.031 | 0.940 | -0.247 | 0.149 | -0.192 | 0.512 | 0.403 | 0.037 |

| | | | | | | | | | | | | | | | |
|---|---|---|---|---|---|---|---|---|---|---|---|---|---|---|---|
| *AL4* | AT5G26210 | 0.217 | 0.237 | 0.277 | 0.426 | -0.033 | 0.827 | 0.108 | 0.735 | 0.274 | 0.158 | 0.118 | 0.419 | 0.402 | 0.042 |
| *CDF3* | AT3G47500 | 0.333 | 0.226 | 0.232 | 0.507 | 0.363 | 0.024 | 0.002 | 0.996 | -0.027 | 0.906 | 0.112 | 0.805 | 0.362 | 0.042 |
| *TCP22* | AT1G72010 | 0.209 | 0.382 | -0.031 | 0.900 | 1.045 | 0.000 | 0.058 | 0.891 | -0.057 | 0.561 | -0.005 | 0.990 | 0.253 | 0.045 |
| *ATL80* | AT1G20823 | 0.126 | 0.792 | 0.284 | 0.684 | 0.117 | 0.662 | 1.232 | 0.150 | 0.569 | 0.410 | 1.021 | 0.024 | 0.209 | 0.618 |

**Supplemental Table S5.** Identified candidate genes mediating acclimation to combined high temperature and drought (HTD) and/or submergence followed by drought (PSD) according to the transcriptomic analysis. Indicated are the the commonly used abbreviation (Gene name), AGI gene locus ID, stress type, and selection criteria.

| Gene Name | Locus ID | Stress type | Selection criteria |
|---|---|---|---|
| *HSFB2B* | AT4G11660 | HTD | Upregulated TF |
| *HSF4* | AT4G36990 | HTD | Upregulated TF |
| *WHY2* | AT1G71260 | HTD | Upregulated TF |
| *VIP1* | AT1G43700 | HTD | Upregulated TF |
| *ATL66* | AT3G11110 | HTD/PSD | Upregulated TF |
| *DEAR3* | AT2G23340 | HTD | Upregulated TF |
| *NF-YB2* | AT5G47640 | HTD/PSD | Upregulated TF/Hub genes in GRN |
| *ZFP4* | AT1G66140 | HTD/PSD | Upregulated TF |
| *TLP7* | AT1G53320 | HTD | Upregulated TF |
| *BZIP28* | AT3G10800 | HTD | Upregulated TF |
| *ERF3* | AT1G50640 | HTD | Upregulated TF |
| *BZIP17* | AT2G40950 | HTD | Upregulated TF |
| *HB1* | AT3G01470 | HTD/PSD | Upregulated TF |
| *ATL24* | AT1G74410 | HTD/PSD | Upregulated TF |
| *ELF6* | AT5G04240 | HTD | Upregulated TF |
| *SE* | AT2G27100 | HTD | Upregulated TF |
| *SCL14* | AT1G07530 | HTD/PSD | Upregulated TF |
| *CDC5* | AT1G09770 | HTD | Upregulated TF |
| *PCFS4* | AT4G04885 | HTD | Upregulated TF |
| *MYBR1* | AT5G67300 | HTD | Upregulated TF |
| *ARID5* | AT3G43240 | HTD | Upregulated TF |
| *AT1G01930* | AT1G01930 | HTD | Upregulated TF |
| *TCP23* | AT1G35560 | HTD | Upregulated TF |
| *TCP9* | AT2G45680 | HTD | Upregulated TF |
| *AT5G12310* | AT5G12310 | HTD | Upregulated TF |
| *ILR3* | AT5G54680 | HTD | Upregulated TF |
| *ATL13* | AT3G60080 | HTD | Upregulated TF |
| *BEH4* | AT1G78700 | HTD | Upregulated TF |
| *TCP22* | AT1G72010 | HTD/PSD | Upregulated TF |
| *SCL1* | AT1G21450 | HTD | Upregulated TF |
| *AT1G80400* | AT1G80400 | HTD | Upregulated TF |
| *HB2* | AT4G16780 | HTD | Upregulated TF |
| *SCL8* | AT5G52510 | HTD/PSD | Upregulated TF |
| *BEL1* | AT5G41410 | HTD | Upregulated TF |
| *MYB34* | AT5G60890 | HTD | Upregulated TF |

| | | | |
|---|---|---|---|
| *CGA1/GNL* | AT4G26150 | PSD | Upregulated TF |
| *NAC083* | AT5G13180 | PSD | Upregulated TF |
| *WRKY17* | AT2G24570 | PSD | Upregulated TF |
| *FBH2* | AT4G09180 | PSD | Upregulated TF |
| *ERF34* | AT2G44940 | PSD | Upregulated TF |
| *RGA1* | AT2G01570 | PSD | Upregulated TF |
| *DOF2* | AT3G21270 | PSD | Upregulated TF |
| *DOF1* | AT1G51700 | PSD | Upregulated TF |
| *HB34* | AT3G28920 | PSD | Upregulated TF |
| *AT3G10760* | AT3G10760 | PSD | Upregulated TF |
| *AL4* | AT5G26210 | PSD | Upregulated TF |
| *CDF3* | AT3G47500 | PSD | Upregulated TF |
| *TCP22* | AT1G72010 | PSD | Upregulated TF |
| *ATL80* | AT1G20823 | PSD | Upregulated TF |
| *ABI5* | AT2G36270 | HTD/PSD | Hub genes in GRN |
| *MYBR1* | AT5G67300 | HTD | Hub genes in GRN |
| *AT1G16080* | AT1G16080 | HTD | Hub genes in GRN |
| *ABF3* | AT4G34000 | HTD | Hub genes in GRN |
| *GBF3* | AT2G46270 | HTD/PSD | Hub genes in GRN |
| *ABF4* | AT3G19290 | HTD | Hub genes in GRN |
| *TLP18.3* | AT1G54780 | HTD | Hub genes in GRN |
| *ABF1* | AT1G49720 | HTD/PSD | Hub genes in GRN |
| *CRL* | AT5G51020 | HTD | Hub genes in GRN |
| *GBF2* | AT4G01120 | PSD | Hub genes in GRN |
| *CCA1* | AT2G46830 | PSD | Hub genes in GRN |
| *PRK* | AT1G32060 | PSD | Hub genes in GRN |
| *AT5G65840* | AT5G65840 | PSD | Hub genes in GRN |
| *AT5G62140* | AT5G62140 | PSD | Hub genes in GRN |
| *GUN4* | AT3G59400 | PSD | Hub genes in GRN |
| *AT1G66130* | AT1G66130 | PSD | Hub genes in GRN |

**Supplemental Table S6.** T-DNA insertion mutants used for HTD in this study. Indicated are AGI gene locus ID, the commonly used abbreviation of the mutants, SALK number or resource publications, primers used for genotyping (including forward, reverse and insertion primers), publications that confirmed the knockout of the corresponding SALK line, status of homozygosity and the insertion position of T-DNA.

| Locus ID | Mutant Name | SALK ID/Resource | Forward primer | Reverse primer | Insertion primer | Studies confirmed knockout function | Homozygous availability | T-DNA insertion position |
|---|---|---|---|---|---|---|---|---|
| AT1G50640 | *erf3* | SALK_200697C | CGATCTGGGCATTAATTTTG | CCTCCTCCTCCTAATCTCCG | CGTCAAGCTCTAAATCGGGG | | No | |
| AT2G23340 | *dear3* | SALK_116495 | TGATCGGCCTTCACTTTTTC | ATGCATCCAACTCGATTTCC | CGTCAAGCTCTAAATCGGGG | | No | |
| AT3G43240 | *arid5* | SALK_111627C | AGCCCACTCTCCACATGATC | TGCCAACCAGAAAAAGATCC | CGTCAAGCTCTAAATCGGGG | (Zhao et al., 2021) | Yes | Exon |
| AT5G54680 | *ilr3-1* | (Rampey et al., 2006) | AGAAATCGCTATGGAATTGTTTATGGTTCT | CATTGTTCAACAGATAACAGTTACGATGAT | - | (Tissot et al., 2019) | Yes | Exon |
| AT2G40950 | *bzip17* | SALK_004048C | GAGAGGATGCTGACGCTAGTG | ATTACCTGCAACACCTTCACG | ATTTTGCCGATTTCGGAAC | y(Cifuentes-Esquivel et al., 2018) | Yes | Exon |
| AT3G10800 | *bzip28* | SALK_132285 | CACCATTAATTTCTTAACCCAAGC | GTTGCCTTAAAGCGACATTCTC | GCGTGGACCGCTTGCTGCAACT | (Gao et al., 2008) | Yes | Exon |
| AT3G10800 /At1g42990 | *bzip28 bzip60* | SALK_132285 (BZIP28), SALK_050203 (BZIP60) | GGAAGAAAAGTCCTCTCGGAG (BZIP60) | CACAGCATCATCGTCTCCTTC (BZIP60) | CGTCAAGCTCTAAATCGGGG | (Sun et al., 2013) | Yes | Exon (Both) |
| AT1G43700 | *vip1* | SALK_080470C | CAAGGGATGCAACAATCATG | CCTCAGGGGAATGGAAATTC | CGTCAAGCTCTAAATCGGGG | | Yes | 3'UTR |
| AT1G78700 | *beh4* | SALK_025953 | TTTTTGGGAAAGCAAGTTGG | AAAAGATTACGGTCGGGGAC | CGTCAAGCTCTAAATCGGGG | | No | |

| | | | | | | | | |
|---|---|---|---|---|---|---|---|---|
| AT1G66140 | *zfp4* | SALK_038923C | TGCCATTTATGTGTCAGCAAC | CTGCTGCTCCGGGTTACTAC | CGTCAAGCTCTAAATCGGGG | (Joseph et al., 2014) | Yes | 5'UTR |
| AT1G01930 | *at1g01930* | - | - | - | - | | No | |
| AT5G04240 | *elf6-3* | SALK_074694C | AGCGCAAAAGCAATAAGACC | TACCCAAGCCATCGAAAAAG | CGTCAAGCTCTAAATCGGGG | (Keyzor et al., 2021; Yu et al., 2008) | Yes | Exon |
| AT2G27100 | *se-1* | SALK_059424 | AGGACGTGGAGGTTATGGTG | CAGAAGGGAAACTGAGAGCG | CGTCAAGCTCTAAATCGGGG | (Lobbes et al., 2006) | Yes | CDS |
| AT4G04885 | *pcfs4-1* | SALK_102934C | GAAAGAGGATGCATGGCTTC | ATGCATCATGTGTCGTTTGG | CGTCAAGCTCTAAATCGGGG | | No | |
| AT5G12310 | *at5g12310* | SALK_123983C | TACACCACCCAAAATTTGGC | ATTTTCTCAATGTCCACCGC | CGTCAAGCTCTAAATCGGGG | | Yes | CDS |
| AT1G74410 | *atl24* | SALK_046988 | TCTTCTCCGACTTGGACCAG | AGTTTCGATGTCATGCATGC | CGTCAAGCTCTAAATCGGGG | | No | |
| AT1G80400 | *at1g80400* | SALK_023687C | AAGGCTGGGTTTCAAGTTTG | TTGGCCTCAATTAATTTGGG | CGTCAAGCTCTAAATCGGGG | | Yes | Exon |
| AT3G60080 | *btl13* | SAIL_1153_F01 | CATCCAACGTTGTTCGTTTG | AATCATCGGTGGAACAGAGC | CTGAATTTCATAACCAATCTCG | | Yes, Col-3 background with multiple loci | 5'UTR |
| AT3G11110 | *atl66* | WiscDsLoxHs207_06D | AAACAACCCAGCTGCTTACG | AGTCTGGTGTTGCCAATTCC | TTTCTCCATATTGACCATCATACTCATTG | | Yes, with multiple loci | 3'UTR |
| AT5G47640 | *nf-yb2* | SALK_025666 | ATCACCAACACCAATGTTCG | AGATCAATTCCAATCCGACG | CGTCAAGCTCTAAATCGGGG | (Sato et al., 2019) | Yes | Exon |

| | | | | | | | | |
|---|---|---|---|---|---|---|---|---|
| AT1G21450 | *scl1* | SALK_102071 | TTTTGGGAGATTACCAAGCG | TCTGGGAAAGAACGGTGAAG | CGTCAAGCTCTAAATCGGGG | | Yes | Exon |
| AT5G52510 | *scl8* | SALK_129947 | GTCCACCTTCTCCGATATCG | GTGGTGGGTCGGATTTTTAC | CGTCAAGCTCTAAATCGGGG | (Ziemer, 2008) | Yes | CDS |
| AT1G07530 | *scl14* | SALK_126931 | GGTCTCACCACACAAAAATAATG | AGCAGCTTGAAGAGCCAAAG | CGTCAAGCTCTAAATCGGGG | (Fode et al., 2008) | Yes | 5'UTR |
| AT3G01470 | *hb1* | SALK_207381C | CCTCGGATCTGAGCTTATCG | GGTATAAAAGACGGCGCTTG | CGTCAAGCTCTAAATCGGGG | (Capella et al., 2015) | Yes | 5'UTR |
| AT4G16780 | *hb2* | SALK_106790C | CTCACGACTCAACGATCTAACC | ATTCCTCTTGAGCCTTGTGG | CGTCAAGCTCTAAATCGGGG | (Stamm et al., 2012) | Yes | Exon |
| AT4G11660 | *hsfb2b* | SALK_137357C | AAAAAGAATACATTTTCCAACTTATCTC | TTCGTTCCACGAGATCAATTC | CGTCAAGCTCTAAATCGGGG | | Yes | Exon |
| AT4G36990 | *hsf4* | SALK_070065C | TCTCTTTTCGTGCGTGTTTG | AAGGTTTTCTCACCAACCCC | CGTCAAGCTCTAAATCGGGG | | Yes | 5'UTR |
| AT5G60890 | *myb34* | WiscDsLox424F3 | TCCGGCGAATTTTCAATAAC | ATGGTGAGGACACCATGTTG | TTTCTCCATATTGACCATCATACTCATTG | (Frerigmann and Gigolashvili, 2014) | Yes | CDS |
| AT5G67300 | *myb44* | SALK_008606 | TTGTCAATTTGTCATGCACTG | TCGCCCATTTATTACCGAAC | CGTCAAGCTCTAAATCGGGG | (Zhao et al., 2016) | Yes | Exon |
| AT1G09770 | *cdc5* | SAIL_207_F03 | TGTACGACCCACAATAGGAGC | ATACTTGAATGCCGAAACCC | CTGAATTTCATAACCAATCTCG | | No | |
| AT2G45680 | *tcp9* | SALK_143587 | TTAAAAATCCCGACGACGAC | GGACCATGACTTAGAGAGGGC | CGTCAAGCTCTAAATCGGGG | | Yes | |
| AT1G35560 | *tcp23* | SALK_203816C | ATATCCGGGCTAACCCCTAG | TCAAGATCACCTAAACCGGG | CGTCAAGCTCTAAATCGGGG | | Yes | 3'UTR |
| AT1G72010 | *tcp22* | SALK_027490 | TGTTGGGGTTCCTATCTTCG | GAGTTAGCTCAGGGTCGTGG | CGTCAAGCTCTAAATCGGGG | (Aguilar Martinez and Sinha, 2013) | Yes | Exon |

| | | | | | | | | |
|---|---|---|---|---|---|---|---|---|
| AT1G53320 | *tlp7* | SALK_120547 | GATTTTCACCCGCGGTATAC | CGAATTTTGTTTGAAGCAACTTC | CGTCAAGCTCTAAATCGGGG | | No | |
| AT1G71260 | *why2* | SALK_016156C | CCATGCACGGGTAAATTTAAC | CGCTTACTTCAAATCCGAGG | CGTCAAGCTCTAAATCGGGG | (Cappadocia et al., 2010) | Yes | CDS |
| AT2G36270 | *abi5-7* | (Nambara et al., 2002) | CGTCAGAGCGAGAAGTAGAG | GCGGGGCGGGGGCACGGGGGGGATTGTTATTATTCTCCTCTGCGAT | - | (Zhao et al., 2020) | Yes | G-to-A mutation at 225 bp |
| AT1G54780 | *tlp18.3* | SALK_109618 | TCGTCCGTTGCTAGTACTGC | TCAAAACCCACCACCTTCTC | CGTCAAGCTCTAAATCGGGG | (Ansari and Lin, 2011) | Yes | Intron |
| AT5G41410 | *bel1* | - | - | - | - | | No | |

**Supplemental Table S7.** List of T-DNA insertion mutants used for PSD in this study. Indicated are AGI gene locus ID, the commonly used abbreviation of the mutants, SALK number or resource publications, primers used for genotyping (including forward, reverse and insertion primers), publications that confirmed the knockout of the corresponding SALK line, status of homozygosity and the insertion position of T-DNA.

| Locus ID | Mutant Name | SALK ID/Resource | Forward primer | Reverse primer | Insertion primer | Studies confirmed knockout function | Homozygous availability | T-DNA insertion position |
|---|---|---|---|---|---|---|---|---|
| AT5G26210 | *al4* | SALK_087658 | TACTGGTCCGAAAAACCCTG | GCGTCAAGGAAAAGCATCTC | CGTCAAGCTCTAAATCGGGG | | No | |
| AT2G44940 | *erf34* | SALK_020979C | TTAAGAGGTGACGCACATGC | CCGAAGAAAACAACTGAGCC | CGTCAAGCTCTAAATCGGGG | | No | |
| AT4G09180 | *fbh2* | SALK_063665C | TTGCATTTTGCAGACATACG | AATCTCGAGAACTTGGTCAGC | CGTCAAGCTCTAAATCGGGG | | Yes | Exon |
| AT1G51700 | *dof1* | SALK_065359 | TGGCTAAAACATTTGACAGGTG | AGGCAAAAGCATGGAATTTG | CGTCAAGCTCTAAATCGGGG | | No | |
| AT3G21270 | *dof2* | SALK_097322C | GAAAAATGGAACCAAAACAAAG | TTCTCAAAAACCGGATTTGG | CGTCAAGCTCTAAATCGGGG | | Yes | |
| AT3G47500 | *cdf3* | GK-808G05 | AATCATCTCCATTCTCTACCCGAG | CATAGTTCCAGTCTTGTGTTACCA | ATATTGACCATCATACTCATTGC | (Corrales et al., 2017) | Yes | CDS |
| AT4G26150 | *gnl (cga1)* | SALK_003995 | CACCGCAACAAAATTTCATG | AGGCAAAAGCATGGAATTTG | ATTTTGCCGATTTCGGAAC | (Chiang et al., 2012) | Yes | Intron |
| AT4G26150/ AT5G56860 | *gnlgnc* | SALK_003995(GNL), SALK_001778 (GNC) | TTTGATCTTGCACTTTTTGGC | AGGCAAAAGCATGGAATTTG | ATTTTGCCGATTTCGGAAC | (Chiang et al., 2012) | Yes | Exon & Intron |
| AT1G66140 | *zfp4* | SALK_038923C | TGCCATTTATGTGTCAGCAAC | CTGCTGCTCCGGGTTACTAC | CGTCAAGCTCTAAATCGGGG | (Joseph et al., 2014) | Yes | 5'UTR |
| AT3G11110 | *atl66* | WiscDsLoxHs207_06D | AAACAACCCAGCTGCTTACG | AGTCTGGTGTTGCCAATTCC | TTTCTCCATATTGACCATCATACTCATTG | | Yes, with multiple loci | 3'UTR |
| AT1G74410 | *atl24* | SALK_046988 | TCTTCTCCGACTTGGACCAG | AGTTTCGATGTCATGCATGC | CGTCAAGCTCTAAATCGGGG | | No | |
| AT1G20823 | *atl80* | SALK_088190 | AGCCATATTTTGTTACGTGAAAG | CTTCTTCAGGCCTTTGTTCG | CGTCAAGCTCTAAATCGGGG | | Yes | Promoter |

| | | | | | | | | |
|---|---|---|---|---|---|---|---|---|
| AT5G47640 | *nf-yb2* | SALK_025666 | ATCACCAACAC CAATGTTCG | AGATCAATTCCAATCCGACG | CGTCAAGCTCTAAA TCGGGG | (Sato et al., 2019) | Yes | Exon |
| AT3G10760 | *at3g10 760* | SALK_011176C | CCTGAAGGTGCT TGAGCTTC | TCAGACTGGTTCGCAAGATG | CGTCAAGCTCTAAA TCGGGG | | Yes | Exon |
| AT1G07530 | *scl14* | SALK_126931 | GGTCTCACCAC ACAAAAATAATG | AGCAGCTTGAAGAGCCAAAG | CGTCAAGCTCTAAA TCGGGG | (Fode et al., 2008) | Yes | 5'UTR |
| AT5G52510 | *scl8* | SALK_129947 | GTCCACCTTCTC CGATATCG | GTGGTGGGTCGGATTTTTAC | CGTCAAGCTCTAAA TCGGGG | (Ziemer, 2008) | Yes | CDS |
| AT2G01570 | *rga1* | SALK_137951 | AAACCTTTTTCAT GAAATTATCGC | ACCGTGGATTCGTTGCTAAC | CGTCAAGCTCTAAA TCGGGG | | Yes | |
| AT3G01470 | *hb1* | SALK_207381C | CCTCGGATCTGA GCTTATCG | GGTATAAAAGACGGCGCTTG | CGTCAAGCTCTAAA TCGGGG | (Capella et al., 2015) | Yes | 5'UTR |
| AT5G13180 | *nac08 3* | SALK_143793 | AATGACCATATT GCCCTTGG | ATCGGTTCTTGAGCCATGAG | CGTCAAGCTCTAAA TCGGGG | (Yang et al., 2011) | Yes | Exon |
| AT1G72010 | *tcp22* | SALK_027490 | TGTTGGGGTTCC TATCTTCG | GAGTTAGCTCAGGGTCGTGG | CGTCAAGCTCTAAA TCGGGG | (Aguilar Martinez and Sinha, 2013) | Yes | Exon |
| AT2G24570 | *wrky17* | SALK_142377 | CAAGAAGCTGC ATCACAAGG | CTTACCGCCGGTACTCTCAC | CGTCAAGCTCTAAA TCGGGG | | Yes | Intron |
| AT3G28920 | *hb34* | SALK_085482C | GAAGACGACGA GGCGTTTAC | GCAACATTCATCAACCAGAGC | CGTCAAGCTCTAAA TCGGGG | (Lee et al., 2022) | Yes | Exon |
| AT2G36270 | *abi5-7* | - | CGTCAGAGCGA GAAGTAGAG | GCGGGGCGGGGGCACGGGGGG GATTGTTATTATTCTCCTCTGCGAT | | | Yes | |
| AT3G59400 | *gun4* | SALK_011461 | GACTCTGCCCAT GTGCCTAG | AGGTGAAAACAATCTCCCCC | CGTCAAGCTCTAAA TCGGGG | (Fölsche et al., 2022) | Yes | Exon |

**Supplemental Table S8.** Relative values of measured phenotypic traits of Arabidopsis mutants compared to wild-type plants under HTD. Indicated are genotypes (mutants or wild-type plants) and the values of measured traits relative to the corresponding wild-type plants. Asterisks represent significant differences between mutants and the corresponding wild-type plants for the particular trait ($p < 0.05$, unpaired t-test).

| Genotype | Relative total leaf area | Relative petiole length | Relative blade length | Relative leaf number | Dry weight at wilting | Leaf number at wilting | Day to wilting | %SWC at wilting |
|---|---|---|---|---|---|---|---|---|
| Col-0 | 1.000 | 1.000 | 1.000 | 1.000 | 1.000 | 1.000 | 1.000 | 1.000 |
| *abi5-7* | 0.631* | 1.720* | 1.245 | 0.865* | 0.615* | 0.842* | 0.953 | 1.084* |
| *elf6-3* | 0.783 | 0.969 | 0.986 | 0.881* | 0.863 | 0.782* | 0.884* | 1.148* |
| *at5g12310* | 0.811 | 1.182 | 1.023 | 0.958 | 1.342 | 1.088 | 0.913* | 1.041 |
| *vip1* | 1.011 | 1.735 | 1.133 | 0.997 | 1.114 | 0.998 | 0.902* | 1.040 |
| *zfp4* | 0.978 | 1.069 | 0.981 | 0.969 | 0.912 | 0.924 | 0.896 | 1.107* |
| Col-3 | 0.931 | 0.793 | 0.861 | 0.992 | 1.240 | 1.045 | 0.959 | 1.025 |
| *btl13* | 0.868* | 1.883* | 1.490* | 0.979 | 0.745* | 0.923* | 1.081 | 0.945 |
| *bzip17* | 0.737* | 0.766 | 0.742* | 0.892* | 1.076 | 1.013 | 0.980 | 1.020 |
| *hb2* | 1.215* | 1.714* | 1.411* | 1.022 | 1.015 | 0.988 | 1.062 | 0.964 |
| *nf-yb2* | 1.059 | 2.218* | 1.363* | 0.902* | 0.780 | 0.934 | 0.976 | 1.071 |
| *ilr3-1* | 0.749* | 1.898 | 0.995 | 0.878 | 0.512* | 0.868 | 0.967 | 0.990 |
| *why2-1* | 0.679* | 1.598 | 1.159 | 0.996 | 1.077 | 1.086* | 1.026 | 1.010 |
| *se-1* | 0.636* | 1.028 | 0.907 | 0.979 | 0.920 | 0.906* | 0.987 | 1.035 |
| *atl66* | 1.285* | 1.241 | 1.384* | 1.005 | 0.982 | 0.988 | 0.992 | 1.055 |
| *tlp18.3* | 1.480* | 2.956* | 1.362 | 0.982 | 0.967 | 0.987 | 1.055 | 0.994 |
| *bzip28bzip60* | 0.910 | 1.098 | 1.152 | 0.935* | 0.755* | 1.023 | 1.011 | 1.066 |
| *bzip28* | 1.072 | 1.041 | 0.934 | 0.946* | 0.965 | 1.085* | 0.978 | 1.020 |
| *hb1* | 0.713* | 0.960 | 0.934 | 0.991 | 1.050 | 0.986 | 1.044 | 0.989 |
| *hsfb2b* | 0.749* | 1.348 | 0.948 | 0.993 | 0.964 | 0.917 | 1.008 | 0.962 |
| *myb34* | 0.828* | 0.859 | 0.920 | 0.939 | 1.016 | 0.936 | 0.969 | 1.014 |
| *tcp22* | 1.238 | 1.658* | 1.194 | 1.011 | 1.090 | 0.990 | 0.959 | 1.022 |
| *tcp9* | 1.099 | 1.754 | 1.109 | 0.936* | 1.192 | 0.979 | 0.934 | 1.030 |
| *scl14* | 1.005 | 1.841 | 1.114 | 0.930* | 0.853 | 1.004 | 1.021 | 1.029 |
| *at1g80400* | 1.185 | 1.290 | 0.860 | 0.957 | 1.369* | 1.035 | 0.965 | 0.994 |
| *arid5* | 1.220 | 1.229 | 1.089 | 0.975 | 1.571* | 1.025 | 0.946 | 0.991 |
| *scl1* | 0.895 | 1.142 | 0.913 | 0.949 | 1.240 | 1.014 | 0.982 | 1.018 |
| *scl8* | 0.964 | 1.079 | 0.907 | 1.004 | 1.035 | 1.070 | 1.000 | 1.015 |

| | | | | | | | | |
|---|---|---|---|---|---|---|---|---|
| *hsf4* | 1.003 | 1.200 | 1.164 | 0.986 | 0.911 | 0.949 | 1.022 | 0.997 |
| *myb44* | 1.178 | 1.286 | 1.011 | 0.935 | 1.196 | 0.967 | 0.996 | 1.000 |
| *tcp23* | 0.840 | 1.199 | 0.966 | 0.934 | 1.190 | 0.966 | 0.970 | 1.012 |

**Supplemental Table S9.** Relative values of measured phenotypic traits of Arabidopsis mutants compared to wild-type plants under PSD. Indicated are genotypes (mutants or wild-type plants) and the values of measured traits relative to the corresponding wild-type plants. Asterisks represent significant differences between mutants and the corresponding wild-type plants for the particular trait ($p < 0.05$, unpaired t-test).

| Genotype | Relative total leaf area | Relative petiole length | Relative blade length | Relative leaf number | Dry weight at wilting | Leaf number at wilting | Day to wilting | %SWC at wilting |
|---|---|---|---|---|---|---|---|---|
| Col-0 | 1.000 | 1.000 | 1.000 | 1.000 | 1.000 | 1.000 | 1.000 | 1.000 |
| *nac083* | 1.117 | 1.411* | 1.469* | 0.929 | 0.856 | 0.885 | 0.947 | 1.138* |
| *hb1* | 1.570* | 1.366 | 1.276* | 1.020 | 0.876 | 1.008 | 0.988 | 1.127* |
| *gun4* | 1.294 | 1.104 | 1.188 | 0.962 | 0.482* | 0.764* | 0.881 | 1.194* |
| *abi5-7* | 1.369* | 1.354 | 1.057 | 0.992 | 0.842 | 0.943 | 0.888* | 1.198* |
| *at3g10760* | 0.867 | 1.176 | 1.045 | 0.895* | 0.978 | 0.948 | 0.898* | 1.169* |
| *atl80* | 0.985 | 0.858 | 0.755 | 0.942 | 0.720 | 0.928 | 0.868* | 1.255* |
| *dof2* | 1.088 | 1.093 | 1.250 | 0.920* | 0.649* | 0.815* | 0.889 | 1.129 |
| *scl*14 | 1.386* | 0.973 | 1.002 | 1.000 | 0.848 | 0.912* | 0.981 | 1.025 |
| *hb34* | 1.016 | 0.631* | 0.760 | 0.904* | 0.830 | 0.933 | 0.993 | 0.999 |
| *tcp22* | 1.231 | 1.189 | 1.346* | 0.923* | 1.093 | 0.966 | 0.965 | 1.057 |
| *nf-yb2* | 1.004 | 1.132 | 1.352* | 0.875* | 0.947 | 0.930 | 0.985 | 1.061 |
| *zfp4* | 1.030 | 0.868 | 1.120 | 0.884* | 0.911 | 0.880* | 0.963 | 1.094 |
| *gnl*gnc | 0.943 | 0.908 | 0.992 | 0.976* | 0.690* | 0.899 | 1.017 | 0.984 |
| *atl66* | 1.498* | 1.121 | 1.159 | 1.062 | 1.135 | 1.024 | 1.003 | 1.003 |
| *cdf3* | 1.233 | 1.160 | 1.368* | 0.998 | 0.856 | 0.927 | 0.990 | 1.089 |
| *fbh2* | 0.958 | 0.997 | 1.242 | 0.887* | 0.776 | 0.917 | 0.986 | 1.073 |
| *rga1* | 1.090 | 0.970 | 1.344 | 0.924* | 0.824 | 0.892 | 0.993 | 1.066 |
| *scl8* | 1.067 | 0.786 | 0.777 | 0.988 | 0.975 | 0.993 | 0.964 | 1.045 |
| *wrky17* | 0.950 | 0.948 | 1.048 | 1.027 | 1.013 | 1.033 | 1.031 | 0.899 |
| *erf34* | 0.816 | 0.793 | 1.108 | 0.900 | 0.983 | 0.905 | 0.945 | 1.091 |
| *gnl* | 1.171 | 1.066 | 0.977 | 1.026 | 0.930 | 0.970 | 0.947 | 1.096 |
